# In-silico Exploration of Mouse Brain Dynamics by Stimulation explains Functional Networks and Sensory Processing

**DOI:** 10.1101/512871

**Authors:** Andreas Spiegler, Javad Karimi Abadchi, Majid Mohajerani, Viktor K. Jirsa

## Abstract

Sensory and direct stimulation of the brain probes its functional repertoire and the information processing capacity of networks. However, a systematic exploration can only be performed in silico. Stimulation takes the system out of its attractor states and samples the environment of the flow to gain insight into the stability and multiplicity of trajectories. It is the only means of obtaining a complete understanding of the healthy brain network’s dynamic properties. We built a whole mouse brain model with connectivity derived from tracer studies. We systematically varied the stimulation location, the ratio of long- to short-range interactions, and the range of short connections. Functional networks appeared in the spatial motifs of simulated brain activity. Several motifs included the default mode network, suggesting a junction of functional networks. The model explains processing in sensory systems and replicates the in vivo dynamics after stimulation without parameter tuning, emphasizing the role of connectivity.

## INTRODUCTION

Although many structural and functional components of brain networks are known across spatial and temporal scales, there is yet an integration to be performed enabling us to link brain activity to function in cognition, perception and action. The debate on representation of behavior is old and on-going (see for recent discussions, Huys et al., 2014; Krakauer et al., 2017; Pillai & Jirsa, 2017), thereby converging towards acknowledging the dynamic nature of behavior, typically manifest in its capturing as mathematical objects suited to represent dynamics (Structured Flows on Manifold (SFM, see Pillai & Jirsa, 2017), Dynamical Causal Modeling (DCM, Friston et al 2003), Heteroclinic Cycles (Laurent & Rabinovich, 2008). This is less evident for brain dynamics, in which the field remains dominated by static representations of brain activity such as brain states (sleep, awake, seizure, behavior (sniffing, etc.)), prone to neo-phrenological interpretations and localizationist perspectives on the representation of function in brain activations (see, for instance, Dehaene et al., 2015; Calvert et al., 2004 and the discussion therein). To go beyond the sampling of brain states and localization of their activity focus, brain dynamics outside its attractor states needs to be systematically measured and evaluated from a dynamics perspective. Attempts in this domain have been made by groups quantifying resting and task dynamics through sequences of brain activation patterns (micro states: Van de Ville et al., 2010; Michel and Koenig, 2018), fit low-dimensional dynamic systems to event-related potentials (for instance in generative Bayesian models via DCM: David et al., 2006) and estimators of various forms of causality. Such model inversion relies in all cases on the data features selected for the inversion and model identifiability remains a principle source of uncertainty (Raue et al., 2009). *In-silico* brain modeling avoids these issues around model inversion and allows creating brain network dynamics under the constraints of biologically realistic connectivity. When the modeled brain source activity is placed in an appropriate neuroinformatics framework (see for instance http://www.thevirtualbrain.org), then biophysical forward solutions can be computed linking the source signals to functional neuroimaging signals as observed in real world situations. In-silico modeling of stimulated brain activity thus offers an excellent means of exploring brain dynamics inside and outside of attractor states, with a direct link to measurable brain signals.

Brain imaging techniques using electrodes (Buzsáki et al., 2012), two-photon microscopy (Denk et al., 1994; Svoboda & Yasuda, 2006), and voltage-sensitive dye (VSD) imaging (Mohajerani et al., 2013) provide little (but thoroughly precious) direct insight into brain dynamics on the large scale of an individual brain because of the restriction on recordings (e.g., limited number of intraoperative electrodes and limited access to dyeable brain tissue in a single individual). Because the access is limited both to (i) the interventions on the living tissue, and to (ii) the measurements of its activity, systematic *in-vivo* explorations of brain dynamics are difficult to accomplish, especially not feasible in individuals within a given period of time. Brain models however allow performing experiments in silico extensively and systematically. Protocols in such ‘virtual’ experiments can mimic and thus accompany those used in-vivo, for instance, for validation purposes (e.g., behavior tasks, brain stimulation as well as resting state protocols). In addition to that, in-silico experiments allow for manipulations that are lethal in vivo and simply not practical in individual (e.g., removing brain tissue, cutting connections, changing physiology). However, in-silico experiments have the potential to give insights if and only if experimental data is amply incorporated in the modeling (models are biologically informed) and missing information is discussed and tested in terms of assumptions. To perform systematic exploration of the brain dynamics via stimulation in-silico we exploited the Allen Brain Atlas (ABA), which compiles tracer studies from thousands of mice to construct an anatomical model of an average mouse brain, see Figure 1. Such a model is biologically informed concerning its geometry and the connectivity of its parts the brain areas. The structure of the brain constrains the brain function and dysfunctions (and vice versa) that are reflected in brain signals (measured, for example, by electrophysiology, VSD imaging, functional MRI). Our modeling approach of the brain assumes that (i) the brain structure given by the ABA is stable within a period of the measurements (e.g., for one hour), and that (ii) local dynamics at each node of the brain network (forming spatially extended brain areas in the model) provides the brain with activity. The assumption about the brain anatomy is realistic because the structure (i.e., geometry and connectivity) is experimentally derived and concerns the large scale of a brain. The assumption about the dynamics is less grounded because less sufficient evidence is available to infer the local dynamics from measurements. For instance, by using electrophysiology local activity (in a distinct brain structure) can be recorded. However, recordings from a limited number of sites do provide information about characteristic changes in space and time but do not clarify if an activation pattern related to function or dysfunction (e.g., epileptic discharges) originated locally or emerged throughout the actions in the network. Brain stimulation techniques are here an appropriate method to infer local characteristics (e.g., cortical excitability) by probing the formation of characteristic patterns in the brain activity (e.g., spreading and dissipation of activity from the stimulation site). The scope of this study is to probe the mouse brain structure by stimulation. For this purpose we assume that the dynamics at each network node is identical. This assumption is rather idealizing but it allows for equally probing the influence of connectivity and each of its parts by stimulation. That means that differences in the organization of brain activity in the model do not originate from the differences in the local dynamics but solely originate from the large-scale brain network of the average mouse brain derived from the experimental data provided by the ABA. In this sense, we have provided each network node with simple dynamics that allows it to respond to stimulation and incoming input from other brain regions with a damped oscillation in the gamma range (see Figure 2). Rhythmic activity at each node is assumed to allow the transmission times of the connections between brain areas to be effective. For this purpose the rhythms need to be fast (e.g., faster than 40 Hz) and the conduction speed needs to be slow (e.g., less than 1 m/s) in consideration of the length of axonal fibers connecting brain areas (e.g., mean length of about 6.25 mm). We assume a natural frequency of 42 Hz at each network node and 1 m/s of conduction speed - both within biologically plausible ranges (see section ‘Conduction speed and transmission delays’ and section ‘Local areas and their dynamics’ in the Results). By this simple dynamics at each node we probe the experimentally derived large-scale brain network to produce functionally relevant organizations in the activity by stimulation. Dynamics of simple systems (e.g., planar systems can show smooth and rhythmic responses to stimulation) are often included in systems that are more complex (e.g., Jansen-Rit model within certain ranges for each parameter). However, complex behavior (e.g., occurrence of spike-wave complexes (Jirsa et al., 2014) or epileptic discharges in distant brain areas (Proix et al., 2014)) indicates the complexity of systems and therefore cannot be embedded in simple systems. This strategy provides a discussion about the integration and emergence of function and dysfunctions in the brain, especially at the large scale of entire brains (for a review see Deco et al., 2011; and Breakspear, 2017). If large-scale models are able to provide plausible brain dynamics than the large-scale network provides entry points for interventions, useful for diagnostics and therapy among others.

**Figure 1.**
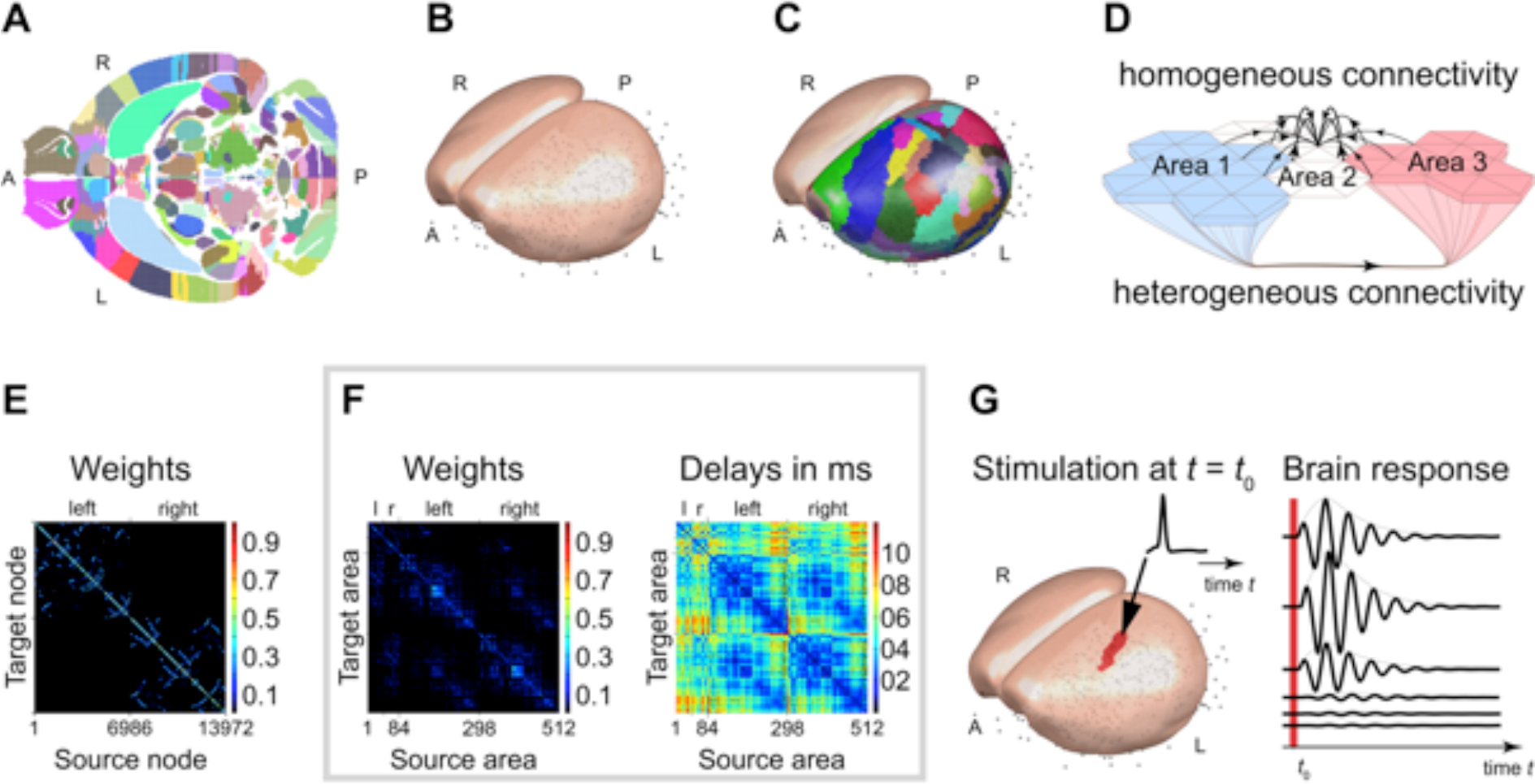
Modeling the mouse brain and simulating the mouse brain’s response to focal stimulation. The ABA database is isotropically sampled with a resolution of 50 μm resulting in a total number of 3,300,706 voxels (1,653,785 voxels for left side), which encode the injection by its spatial location and connectivity among 512 brain areas. The 512 brain areas are listed in Table 2 and Table 3. Panel **A** shows an axial slice of the sampled database. Each dot indicates the center of a voxel and the color indicates the different brain areas. To model large-scale activity of the cortex such as recorded using VSD imaging, the surface of the isocortex is reconstructed from the ABA database (slice shown in panel **A**) using a regular triangle mesh of 7,020 vertices for the left and 6,952 vertices for the right hemisphere. The quality of the mesh is quantified in Figure 11. Panel **B** shows the reconstructed isocortical surfaces. All other areas are lumped in their spatial extent (in the ABA) to point masses (centroids) indicated by the grey circles in panels **B**, **C**, and **G**. Panel **C**: each vertex in the isocortical meshes is assigned to an area by the five closest voxels to the normal vector extended inside the volume of the ABA (slice shown in panel **A**). The areas of the isocortex and the number of vertices per area are listed in Table 1. Panel **D**: The model distinguished two types of structural connectivity. Heterogeneous SC and corresponds to the connections extracted from the ABA such as tracts connecting brain areas over long distances. Homogeneous SC corresponds to gray matter fibers in the isocortex, with short-range connections within a given isocortical area, but also enabling some communication over short distances between neighboring areas. Although in the ABA, cortical area 2 is not connected to areas 1 and 3, it is weakly linked to both areas via a set of short-range SC. Panel **E**: Homogeneous SC matrix for the 13,972 vertices of the isocortex. The synaptic weights are color-coded. The diagonal describes in warm colors the strong SC of adjacent nodes. SC decreases with distance, which is shown in cold colors. SC of nearby nodes are scattered (e.g., blue dots) in panel **E** because each cerebral hemisphere is described by a surface, which makes it impossible to cluster nodes locally along both axes. Note the absence on interhemispheric short-range SC. Panel **F**: Heterogeneous SC for the 512 (84 isocortical plus 428 subcortical) brain areas for weights (left) and time delays (right). Within one hemisphere, the 214 sub-isocortical areas mostly project to the 42 isocortical areas. Some connections between subcortical areas can also be seen. The 42 cortical areas project heavily to both cortical and subcortical areas. Some interhemispheric connections can also been seen. Note also the presence of relatively large time delays. Panel **G**: The stimulation paradigm. The dynamics are simulated without noise. For that reason the activity is flat before stimulation. The brief stimulation causes activity in the targeted brain area, in panel **G** it is the unassigned left primary somatosensory area, and in the entire mouse brain model. The spread and the dissipation of the activity caused by the stimulation depend on the SC and the transmission delays. Each of the 512 cortical and subcortical brain areas is stimulated and the set of characteristic spatial activations of the brain response is extracted and investigated.

**Figure 2.**
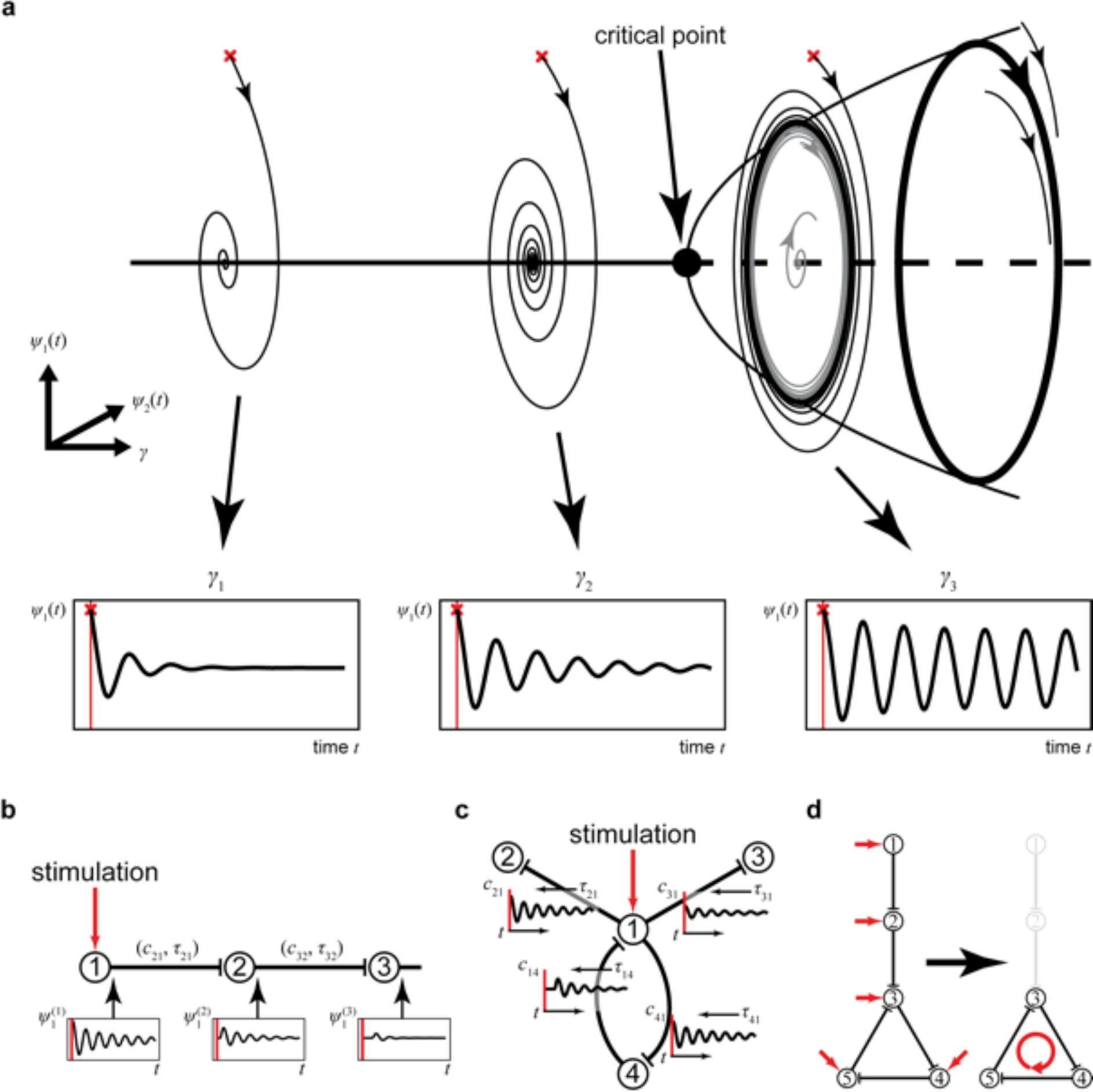
The large-scale brain model works near criticality. Panel **A**: Each node in the model is parameterized by *γ* to operate intrinsically at the same distance from the critical point if unconnected. A node shows zero activity or oscillation (42 Hz) in response to stimulation (red crosses). The activity at each node is described by two time-dependent variables, *Ψ*_1_(*t*) and Ψ_2_(t). The closer a node operates to the critical point, the larger and the longer lasting is the oscillation (compare *γ*_1_ and *γ*_2_). When the critical point is reached, the node intrinsically performs a rhythm of constant magnitude. The model, however, is set so that the critical point is never exceeded. Panel **B**: Principles of activity spreading after stimulation. The damped oscillation generated in the stimulated node (1) is sent via its efferent connections to its target node (2), triggering there, in turn, a damped oscillation with weaker amplitude and faster decay, which then propagates to the next node. Activity *Ψ*^(*j*)^_1_(*t*) of node (*j*) is scaled by *c_ij_* transmitted to node (*i*) via homogeneous and heterogeneous connections (SCs), delayed by *τ_ij_* in the latter case. In such a chain, activity would decay fast. Panel **C**: In the large-scale brain model, multiple activity re-entry points can be found. At any time point, the dynamics of a node is influenced by all incoming activity. The response of the node to stimulation (1) is relayed to linked nodes (2–4), which may be fed back to 1 via 4 and may allow the induced activity to dissipate on a much longer time scale. The network response thus depends upon the SC and allows the network to operate near criticality. Panel **D**: Activation of dynamically responsive networks. Activity after stimulating a node (1 or 2) in a series connection decays fast (as in panel **B**). However, activity may circulate and thus decays slower in a feedback network (of nodes 3–5). Such remaining activity after the initial stimulation decay reveals the so-called dynamically responsive networks (DRNs).

We have used focal stimulation to render the brain dynamics visible in the brain response. The stimulation is brief and, to be effective, defined by the characteristic time of the intrinsic dynamics (gamma activity). We systematically stimulated every single brain area and investigated the subsequently induced activity in the brain responses. We simulated the whole-brain mouse model on a computer using the large-scale brain network simulation engine The Virtual Brain (TVB). For exploring the mouse brain dynamics in silico, we systematically varied three model parameters, namely (i) the ratio to which extend brain areas can interact over short-and long-ranges, (ii) the range of short connections that are homogeneous on the isocortex, and (iii) the location of the focal stimulation. The extent to which information is processed over short-or long-range SC is unclear and focal stimulation is closely related to direct electrical, sensory and optical stimulation. We identified consistent spatial motifs of mouse brain activity after stimulation called dynamically responsive networks (DRNs) using dimension reduction methods. We found the experimentally known functional networks back in the DRNs. Interestingly, the salience and the default mode network spanned several motifs indicating the salience and default mode network to be a junction for functional networks. To test whether the mouse brain activity can be exclusively inferred from the underlying topology, we performed graph theoretic analysis of the model structure. We furthermore investigated the organization of brain activity following sensory pathways. The model structure reflects the sensory systems, namely: the auditory pathways, the visual pathways, the whisker pathways and the pathways for the limbs. Finally, with the developed large-scale mouse brain network model we had, for the first time, the chance to systematically investigate peripheral pathways, especially, the processing of sensory data. We showed that the in-silico brain responses resemble in-vivo brain dynamics after stimulation.

**Table 1.** Mapping of brain areas in the mouse model, that is, areas to brain structures following the division of the Allen Brain Atlas (ABA; http://connectivity.brain-map.org). Because of a total of 512 brain areas in the model, only the number of areas is listed. The names are listed in Table 2 and Table 3.

| Structure | Division | Areas |
| --- | --- | --- |
| Cortex | Isocortex | 42 |
|  | Olfactory areas | 11 |
|  | Hippocampal formation | 10 |
|  | Cortical subplate | 7 |
| Cerebral nuclei | Striatum | 13 |
|  | Pallidum | 8 |
| Thalamus | Sensory-motor-related | 13 |
|  | Polymodal association cortex related | 8 |
| Hypothalamus | Periventricular zone | 3 |
|  | Periventricular region | 11 |
|  | Medial zone | 12 |
|  | Lateral zone | 7 |
| Midbrain | Sensory related | 6 |
|  | Motor related | 17 |
|  | Behavior-state related | 7 |
| Pons | Sensory related | 3 |
|  | Motor related | 7 |
|  | Behavior-state related | 7 |
| Medulla | Sensory related | 9 |
|  | Motor related | 23 |
| Cerebellum | Cortex | 12 |
|  | Nuclei | 3 |

**Table 2.**
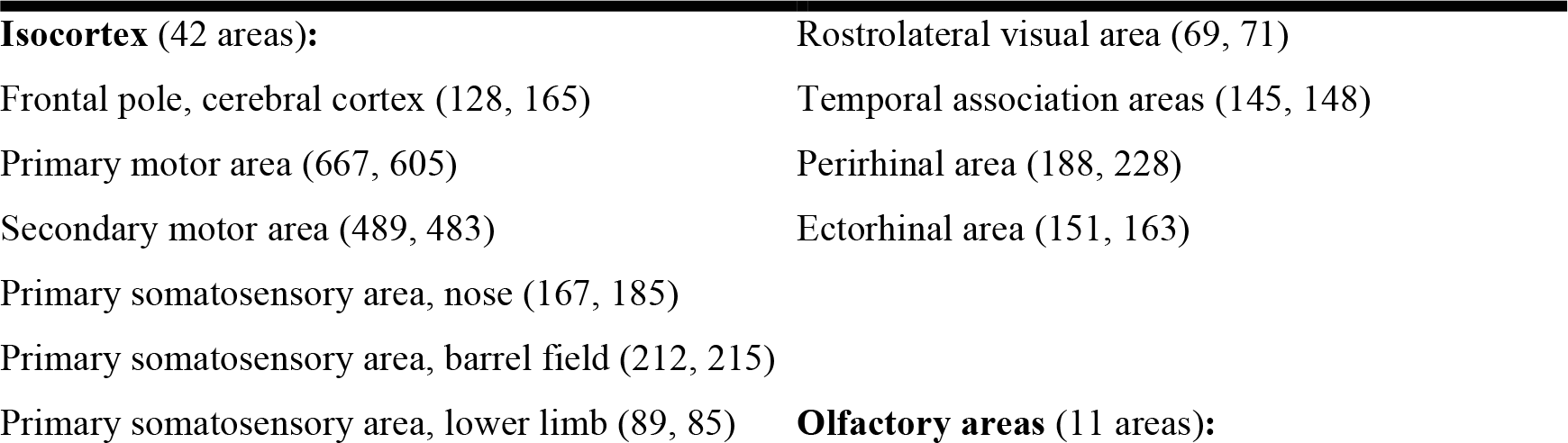
Cerebral brain structures and their division. Number of nodes per area in brackets (left, right).

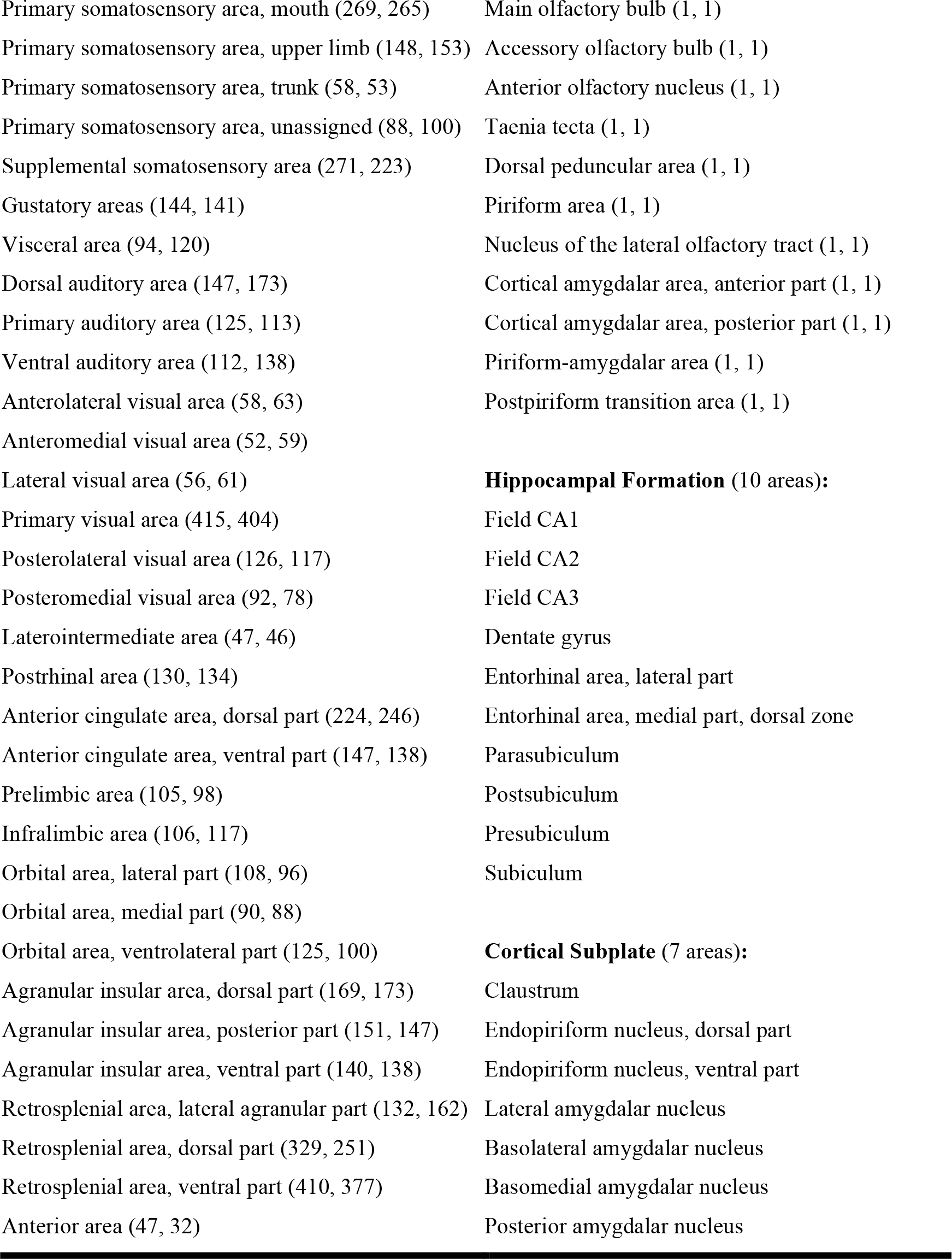

**Table 3.**
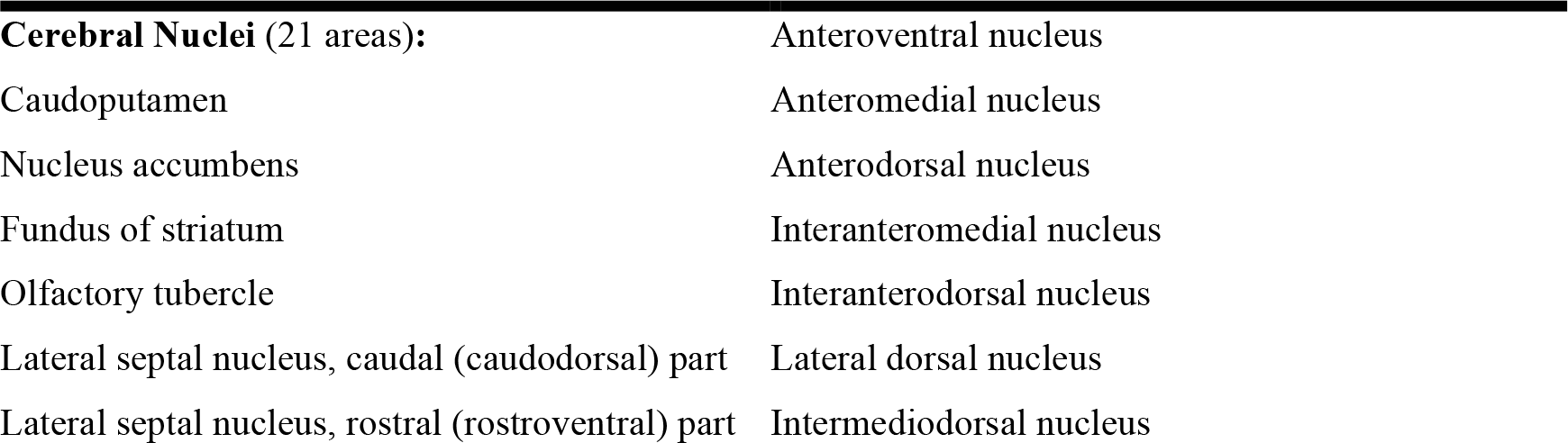
Subcortical brain structures and their division. One network node per area.

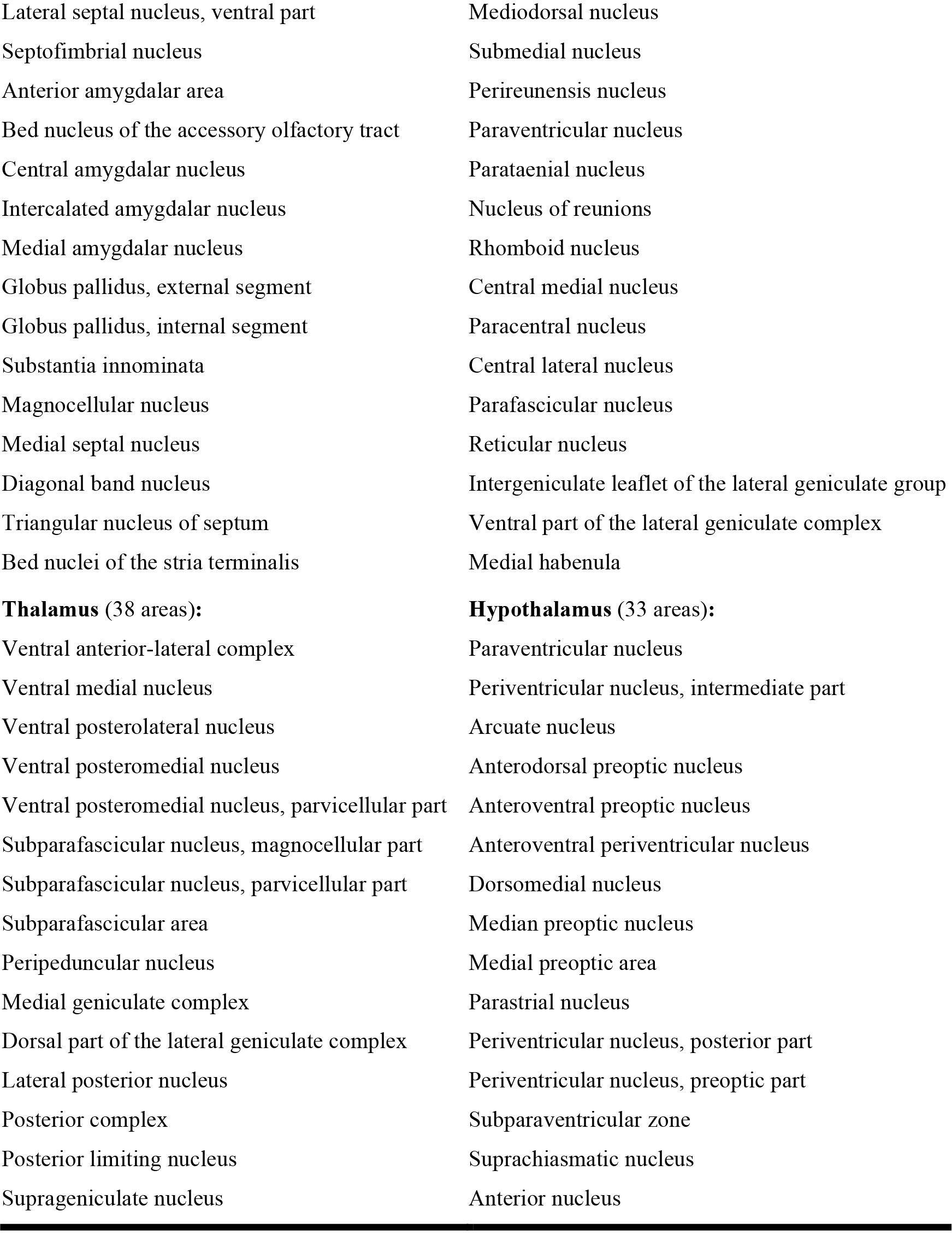

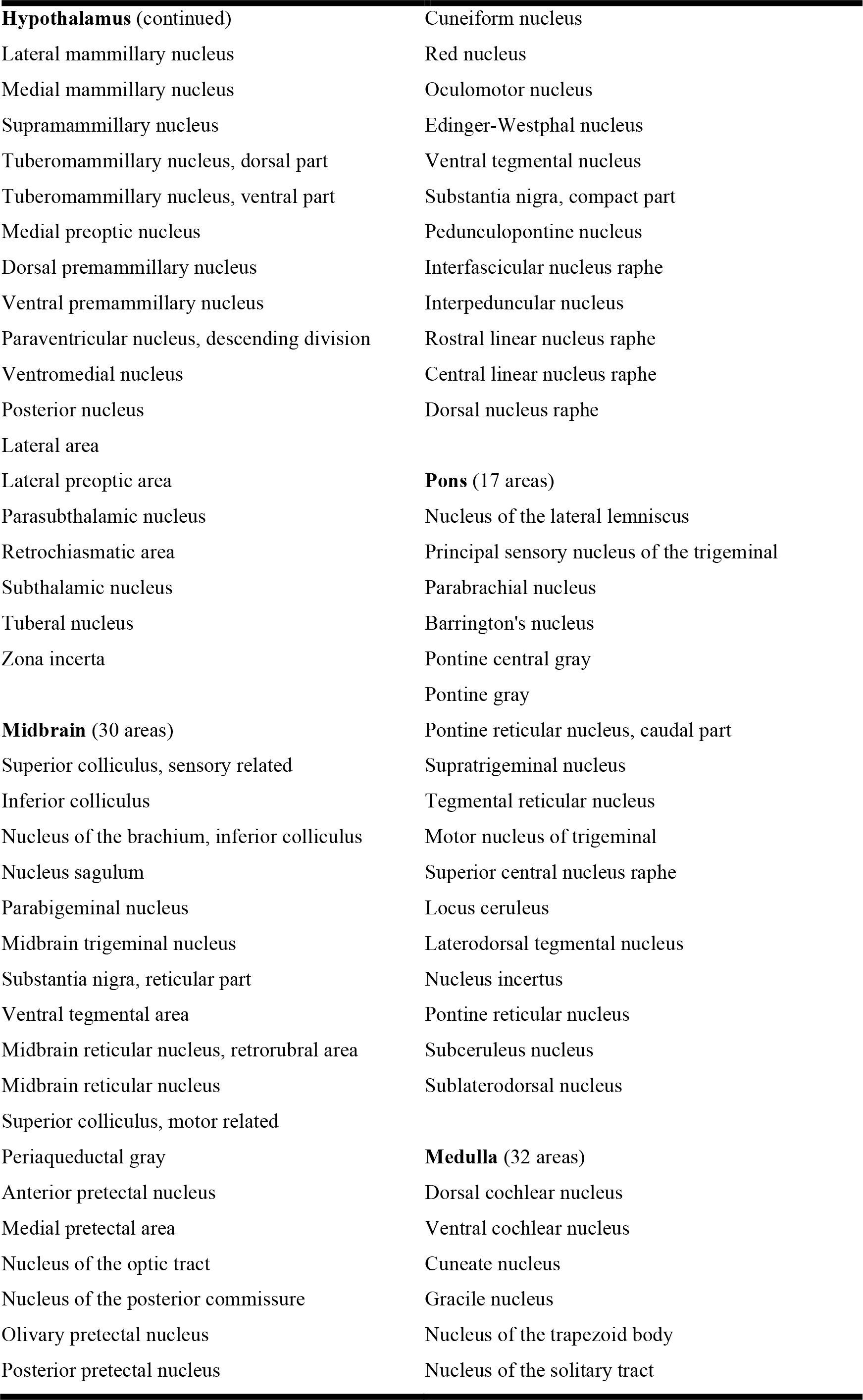

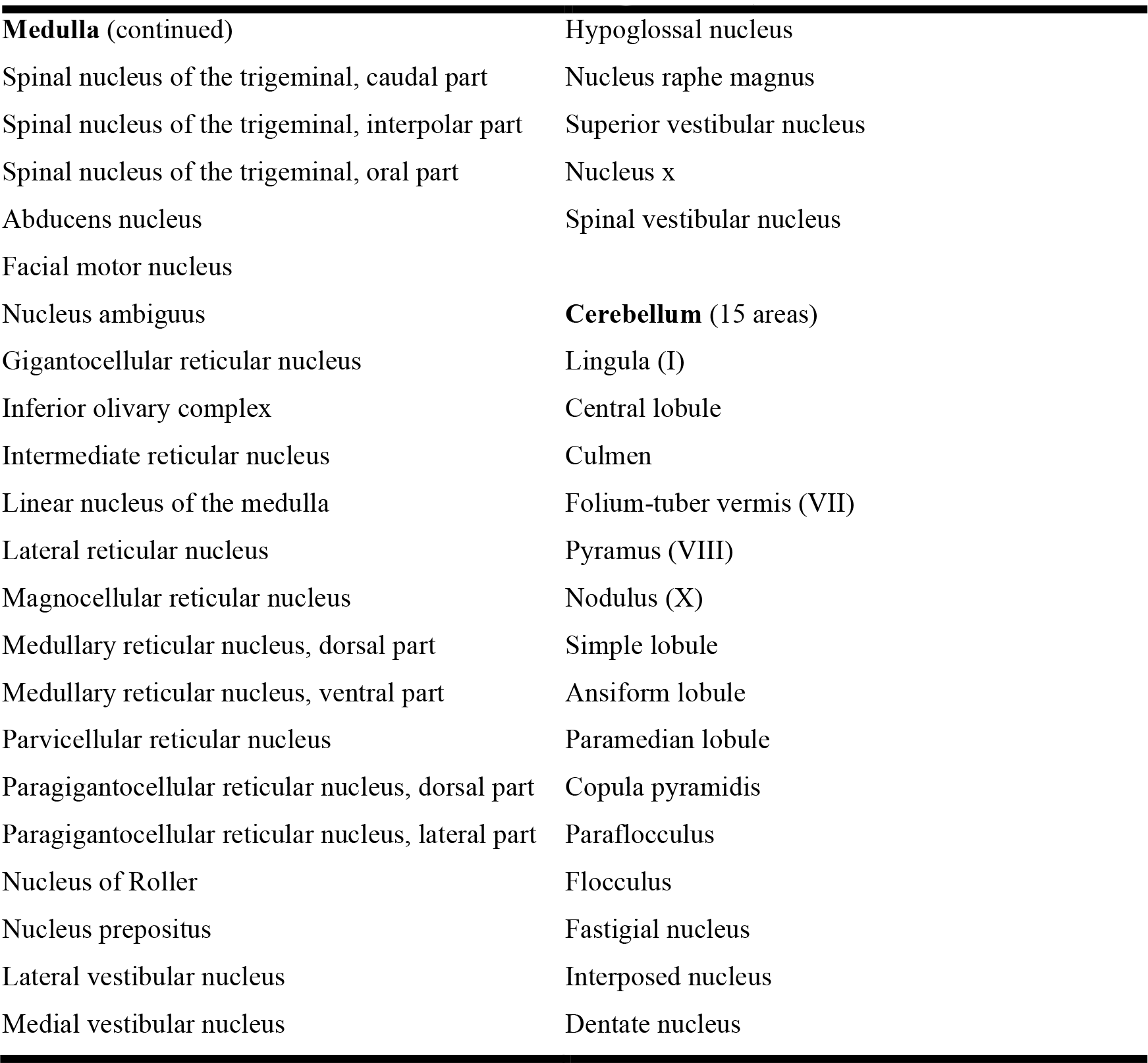

## RESULTS

The result section is divided into two parts. The first part contains the results of the in-silico exploration study by means of focal stimulation. The second part contains the results of compiling the whole mouse model. This modeling work is fundamental to the in-silico exploration but it is also suitable for many more diverse studies, and in this sense, the modeling work stands for itself. The modeling work of the mouse brain was thus not solely driven by the goal of performing in-silico brain stimulation but is rather driven by the objective of obtaining a systemic and holistic description of the brain. We have used brain stimulation as an appropriate concept and the results emphasize the potential of dynamic models to study the functional repertoire of the brain and the information processing capacity of networks.

### In-silico exploration of mouse brain dynamics by means of stimulation

The model allows each brain areas to communicate through short- and long-range connections with other nearby or distant areas. The local dynamics (at each brain area) distinguish both types of structural connectivity in the mathematical description, where a model (order) parameter α sets the ratio and the weight of short-and long-range connections. The parameter α simply scales all long-and all short-range connectivity weights (regarding absolute values) with no effect on the relative weights (between short-range connections or between long-range connections).

We consider this ratio of homogeneous SC to heterogeneous SC as a degree of freedom and performed a parametric study. The ratio has been estimated. For instance, Braitenberg and Schüz (1998) assessed that pyramidal cells have synapses in equal shares from long-range and local axons. However, the ratio of homogeneous SC to heterogeneous SC mainly depends on the resolution of the used geometrical model of the cortex, and with that the representation of the SC, and the local dynamic model (e.g., canonical model, neural mass model), which is able to incorporate local connectivity (for more detail, see Spiegler and Jirsa, 2013). At the extremes, (1) *α* = 1, that is, 0% of heterogeneous SC (thus, 100% of homogeneous SC gives two unconnected hemispheres where only nodes within the isocortices are locally and homogeneously connected) only allows activity to propagate locally from an isocortical stimulation site, and (2) *α* = 0, that is, 100% of heterogeneous SC (thus 0% of homogenous SC gives 512 purely heterogeneously connected brain areas with locally unconnected nodes) only allows activity to travel distances with time delays. The connectivity scaling factor *α* is = {0.0 0.2 0.4 0.6 0.8 1.0}.

Furthermore, since the spatial range of homogeneous SC is of μm up to mm (Braitenberg and Schüz, 1998), we also consider it as a model parameter varying between 500 μm and 1000 μm (step size 100 μm). We then systematically stimulate each of the 512 areas with a large range of connectivity parameter values (for the ratio and the spatial range), resulting in a total number of 18,432 simulation trials.

The dynamically responsive networks (DRNs) are extracted by (i) determining the induced component of generated brain activity after stimulation (subtracting the evoked response at the stimulation site, that is, an isolated and unconnected node), (ii) then decomposing the time series (of the 14,400 neural masses) of stimulation-induced activity (in the time period between 250 ms and 750 ms after stimulation onset *t*_0_) into principal components (eigenvectors and corresponding eingenvalues) using the principal component analysis (PCA), (iii) correlating the eigenspaces spanned by the components that cover 99% of the variance in the brain activity, (iv) clustering the eigenspaces, and (v) lumping the clusters after rotating the eigenspaces onto a common basis.

#### Effect of structural connectivity with time delays on the brain response to stimulation

Considering the scaling of structural connectivity at both extremes, that is, 0% of heterogeneous SC and 100% of heterogeneous SC, the formation of dynamically responsive networks (DRNs) due to stimulation is found to be different (see Figure 3). The DRNs of both extremes change in the communication via short-range and long-range connections. The results indicate that both extremes merge in the mix of the two connectivity types rather than reorganize (e.g., due to time delays in the long-range connections). One explanation for this behavior is that the short-range homogeneous connectivity pertains to a minority of 84 areas in the model whereas the long-range heterogeneous connectivity pertains simply all of the 512 areas. The DRNs are most similar when the 512 areas are only linked via the heterogeneous connectivity. The overall similarity decreases with scaling the homogeneous connectivity up (consequently the heterogeneous connectivity down) and clusters of similar DRNs emerge and become clearer (see Figure 3). The clusters due to different stimulation sites are however the same throughout. Within a cluster structures may appear for higher scaling of the homogeneous connectivity, but clusters do not move, split nor merge (as observed in a similar study for human; Spiegler et al., 2016). The behavior in the similarity of DRNs is unaffected by the spatial range of the local connections. Merely the similarities increase with increasing the range of local connections for higher proportions of the short-range connectivity (e.g., 80%). All in all, both SC types of short- and long-range connections in the mouse model act as spatiotemporal filters of local dynamics in the emergent functional network. The results (Figure 3) do not support evidence for a reorganization of dynamics due to interplay between both types of SC in the mouse as compared to human models (Spiegler et al., 2016).

**Figure 3.**
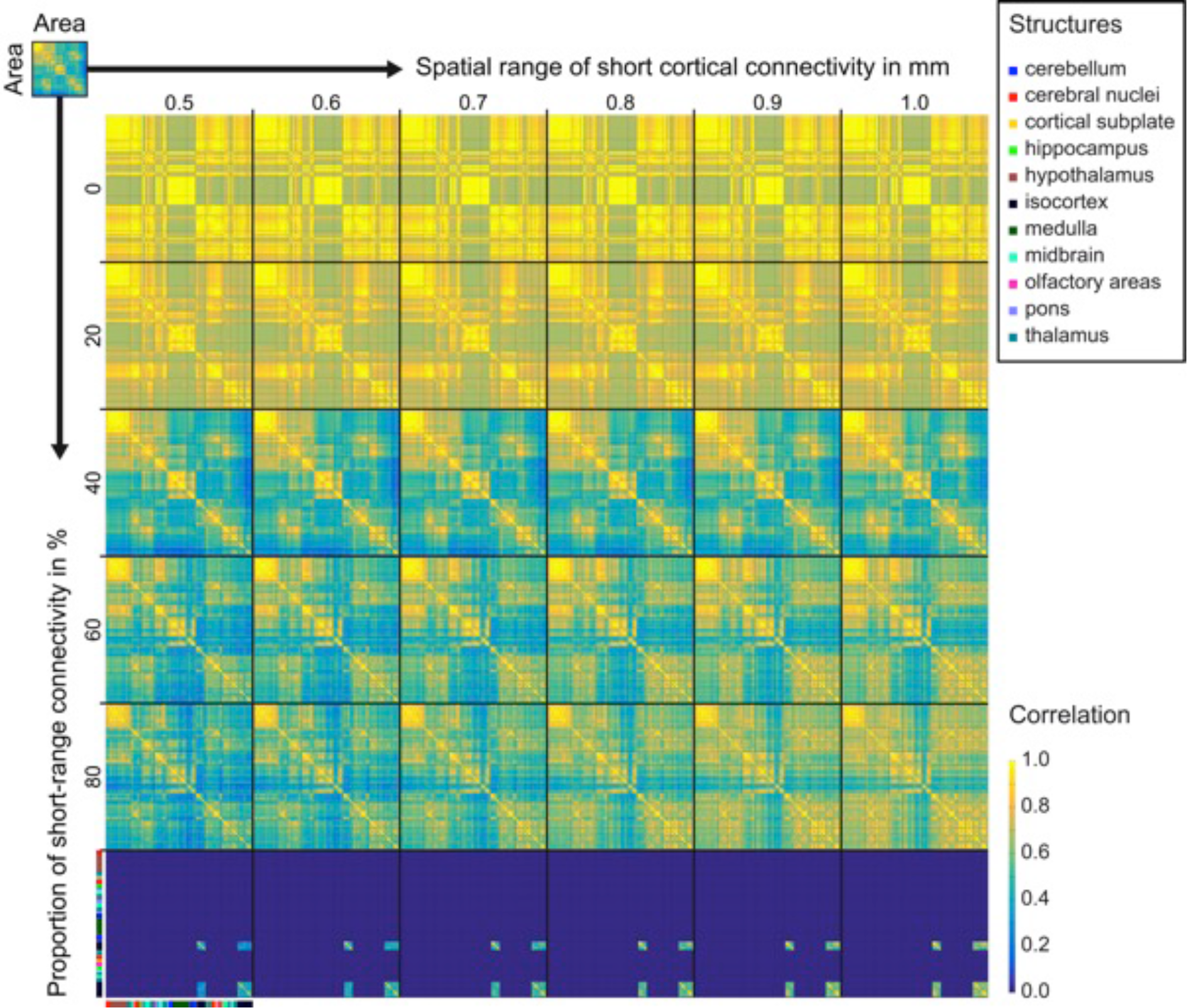
Similarity matrices of the dynamically responsive networks (DRNs) between stimulated brain areas reveal a consistent formation of spatially similar brain activity in the mouse as a function of the proportion of short-and long-range connectivities in the communication between brain areas and the length of short-range connections. Structural connectivity forms clusters of functional networks that are responsive to sets of different stimulation sites. This is summarized in a similarity matrix for a given proportion of homogeneous short-range to heterogeneous long-range connectivities in the model (varied along rows) and for a given length of short-range connections (varied along columns). The similarity of DRNs increases with warmer colors (yellow indicates maximum similarity). All similarity matrices are equally sorted and the color bar presented on the axes of the matrix for a spatial range of 500 μm and 100% of short-range homogeneous connectivity (i.e., no long-range heterogeneous connectivity) indicates the brain structures (e.g., cerebral nuclei) to which a brain area is part of (e.g., nucleus accumbens and fundus of striatum are cerebral nuclei in the mouse model). The results indicate that long-range heterogeneous and short-range homogeneous connections in the mouse shape functional networks but changes in the structural connectivities not necessarily cause reorganization of networks. The similarity matrices of the DRNs to stimulation were almost unchanged with changes in the length of short-range connections (see the similarity matrices along the columns). The short-range heterogeneous connectivity (see the row of 0% short-range homogeneous connectivity) supported DRNs that differ from the DRNs supported the short-range homogeneous connectivity (see the row of 100% short-range homogeneous connectivity). Both similarity matrices merge in the mix of short-and long-range connections (see the similarity matrices along the rows).

Whereas the modeling work of the human brain mainly comprises cortical areas and their connectivities (i.e., local homogeneous short-range connections and connections via long-range white matter fiber tracts), more weight is given to subcortical areas in this mouse model. The findings indicate that for the mouse the homogeneous short-range SC in the isocortex is not sufficient for observing reorganizations of functional networks due to changes in the structural connectivities. Furthermore, the transmission time delays between areas are found to be important in the emergence of the DRNs but less important in the filter action and the interplay of the two connectivity types.

#### Dynamically responsive networks (DRNs)

The dynamically responsive networks (DRN) convey 99% of the stimulation-induced energy in the brain composed by three principal components. By correlating the DRNs of different stimulation sites and performing a permutation test, we identified twelve clusters that are consistent throughout the systematic parameter exploration. The 14,400 time series were used for extracting and correlating the DRNs. On this fine-grained level of 14,400 nodes, the isocortex is overrepresented (with 13,972 nodes), whereas on the coarse-grained level of the 512 brain areas the sub-isocortical areas (e.g., thalamus, cerebral nuclei) form the majority.

To match the DRNs with the voltage-sensitive dye (VSD) imaging data, which mainly record brain activity from the isocortices, the clustering of the responsive network to stimulation of different sites was performed on the fine-grained level of the nodes.

The in-silico exploration provides a spatiotemporal map of consistent formation of a sequence of spatially similar brain activity and a map indicating which brain areas need to be stimulated (Figure 4, Panel **A**: the similarity matrix) to ‘push’ the brain in a particular state (see Figure 5 and Figure 4, Panels **B**, **C**: DRNs of the clusters 1 and 2 in Panel **A**). Stimulations specifically bias widespread functional networks. Spatial proximity of stimulation does not necessarily predict the similarity of induced activities. The same spatial organization of brain activity can be induced by stimulation of several sites scattered all over the brain.

**Figure 4.**
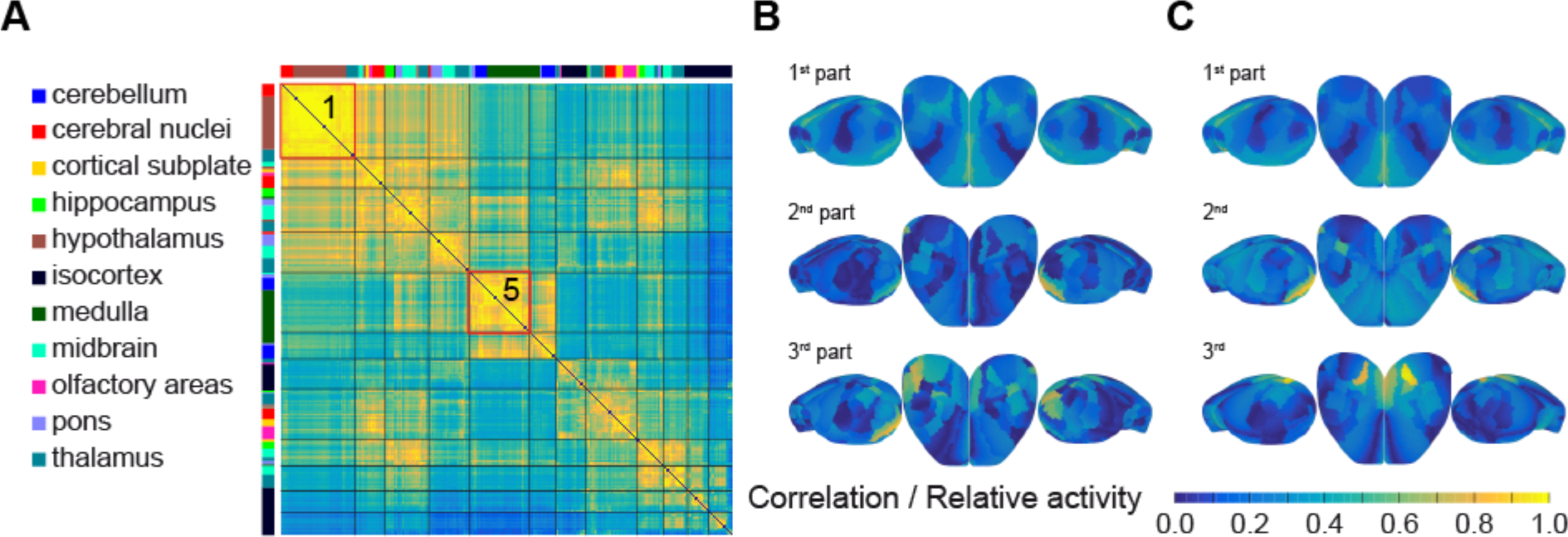
Specific focal stimulations activate similar networks. Panel **A** is the similarity matrix of the dynamically responsive networks (DRNs) for stimulation of the 512 different subareas. The stimulation sites and the DRNs form 12 clusters (separated by the black lines). This suggests that similar functional networks are activated in the mouse brain in response to stimulation of the different sites within a cluster. The color bar on the *x*-and on the *y*-axis indicates the structures to which a brain area is part of. The similarity matrix in panel **A** is for a spatial range of 500 μm and 40% of short-range homogeneous connectivity (i.e., 60% of long-range heterogeneous connectivity). Similarity of DRNs is color coded (warmer color means more similarity). Panels **B** and **C** show the spatial organization of brain activity in the isocortex that corresponds to the clusters 1 and 5 as an example from the left, top, and right view. Three parts compose the motif of each DRN. The blue to yellow scale gives the relative contribution of areas to the response. The complete set of motifs, that is, the averaged brain activity of DRNs in each of the 12 clusters in panel **A** (numbered up from top to bottom) is shown in Figure 5.

**Figure 5.**
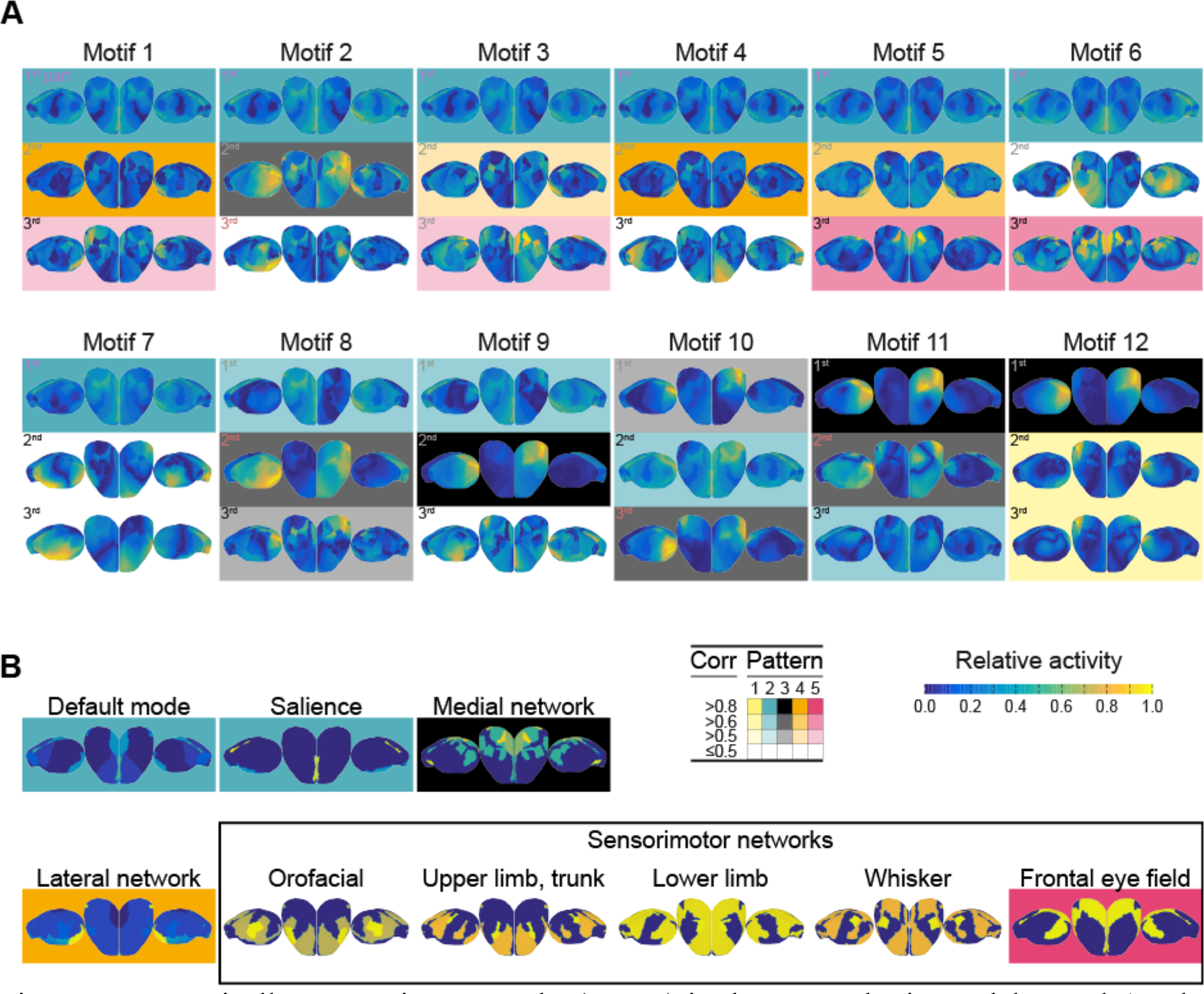
Dynamically responsive networks (DRNs) in the mouse brain model. Panel **A**: The responses of the mouse brain to focal stimulation dissipate in 12 motifs, that is, the DRNs. Each motif proceeds from similar network activity after systematic stimulation of different sites (see Figure 4). Each dynamically responsive network is composed of three components (parts) covering 99% of the activity induced by the stimulation. Each motif, in panel **A**, shows the averaged spatial activity (topography) for each part (1 to 3) across the similar network responses for the isocortices from the left, top, and right view. Similar parts among the motifs are highlighted. The five distinct patterns in the parts entangle the different motifs. The different shade indicates the correlation level (Corr) of the components (no shade, that is, white for not significant correlations and shades: dark, medium, light for significant correlation values greater than 0.8, 0.6, 0.5). Six components are unique (i.e., parts in motifs 4, 6, 7, and 9) and not correlated to other motif parts. This makes motif 7 the most unique one and motif 1 the most entangled motif (followed by the entanglements in motifs 5, 11, and 12). In order to interpret the DRNs, the components were correlated to functional networks. Panel **B**: Experimentally known resting state and functional networks (see description in the Materials and Methods) were compared to each part of each motif. The organization of brain activity in the isocortex is shown in Panel **B** for each of the nine functional networks from the left, top, and right view. Four experimentally known functional networks correlate with the patterns in the parts of the motifs and are highlighted in accordance with Panel **A**. Note that the motifs do not cover the isocortical organization of sensory motor networks (neither left/right symmetric nor lateral) except for the frontal eye field network. The activity of the frontal eye field network within one hemisphere (lateralized organization) corresponds to pattern 4, which is highlighted in magenta. The unique parts in motifs 2, 4, 6, 7, and 9 (see parts in Panel **A** that are not highlighted) are not correlated with one of the nine specific functional networks. Moreover, pattern 1 (highlighted in yellow) in motif 12 does not correspond with any of the nine functional networks. The default mode and the salience networks show similarities at the isocortical level. Both comprised most of the motifs (cyan pattern 2 in motifs 1 to 11). Stimulation responses do activate to a lesser extent the lateral network (orange pattern 4 in motifs 1, and 3 to 5) and the medial network (gray pattern 3 in motifs 8 to 12), where the activity of both networks within one hemisphere (lateralized organization) is reflected. The patterns (1 to 5) in the parts link the motifs (1 to 12). The default mode network (and the salience network) is thus related in the model to the most flexible and sensitive pattern (cyan pattern number 2) that can activate and involve most of the motifs (1 to 11). In both panels, **A** and **B**, the blue to yellow scale gives the relative contribution of isocortical areas to the response. The order of the clusters is consistent with

#### Statistics on DRNs

We have performed statistics on the catalogue of DRNs (Figure 5) to (i) clarify whether the motifs are entangled in their individual parts, and to (ii) discuss how the formation of brain activity due to stimulation does follow the structure. For the first purpose we pair-wise correlated each of the three parts of each of the 24 motifs, that is, twelve motifs by stimulation of left brain areas and twelve motifs by stimulation of right areas (makes a total of 72 parts composing the 24 motifs). The significance level of 1% was Bonferroni corrected by the total number of components. In addition, three thresholds are applied to the correlation values *C* to (*C* > 0.8), good (0.6 < *C* ≤ 0.8), and moderate agreements (0.5 < *C* ≤ 0.6) among the 72 parts of the 24 motifs. Please note that for the rest of the manuscript we refer to the twelve motifs without indicating the stimulation side (left/right) because of the bilateral symmetry (in the spatial organization of brain activity due to left/right stimulation). The analysis reveals a total of five significantly distinct spatial patterns in the motifs of the DRNs, indicating their entanglements. The distinct spatial activation patterns could possibly be the structural links for the transfer of energy, that is, brain activity that may transmit information from one motif to another at rest over time. The reason for that conjecture is the occurrence of the five spatial patterns in the 12 motifs (see the highlighted parts in Figure 5), which, importantly, also allow an activation of more than one motif at a time (i.e., phrasing). The analysis also reveals six components that are rather unique (i.e., parts in motifs 2, 4, 6, 7, and 9 in panel **A** in Figure 5) and are not correlated to other motif parts. Considering the parts of the motifs and the occurrence of patterns, the motif with the number 7 (see, panel **A** in Figure 5) is the most unique one and the motif 1 is the most entangled one (followed by the entanglements in motifs 5, 11, and 12) among all motifs. The pattern with the number 2 is most present in the motifs (part of 11 out of 12 motifs), whereas pattern 1 occurs only within motif 12. The most present pattern, that is, number 2 is also the pattern with the strongest correlations (out of a total of 11 parts, 7 parts are strongly correlated and 4 parts show good correlation), whereas pattern 1 shows only good correlations. Before interpreting the themes of the DRNs by relating the parts of the motifs (and thus the patterns) to experimentally known functional networks, we tested to which extent the topology in the structural connectivity predicts the brain activity due to stimulation.

The brain activity in response to stimulation follows the structural connections from the stimulation sites. Amongst the 13 graph theoretic measures on the structural long-range network (ABA), the direct projections from the site of stimulation best explain the brain response to stimulation though the correlation is rather moderate. The performance of the other twelve graph-theoretic measures (see Materials and Methods) is rather poor, that is, less correlated, weakly predictive and fewer significant configurations. The best prediction resulted from comparing the rank (Kendall’s tau) of the activity at the network nodes after stimulation with the rank of connectivity strengths of the direct projection from the stimulation site. The rank correlation is weak but significant for most connectivity parameter configurations of the ratio *α* of long-range to short-range connections, and the spatial spread *σ* of short-range connections. By utilizing this simple measure on the structural large-scale network, a high correlation value means that the energy induced by the stimulation spreads into areas of the brain that are directly connected to the stimulation site. We corrected the significance level of 5% (Bonferroni). The values of significant correlations are however fairly low with an average maximum value of 221.542 × 10^−3^ across all parts of all motifs and all connectivity parameter configurations (maximum average correlation value of 253.889 × 10^−3^ all 1^st^ parts, 255.936 × 10^−3^ for all 2^nd^ parts, and 338.724 × 10^−3^ for all 3^rd^ parts across all motifs and all connectivity parameter configurations). The percentage of parameter configurations with significant rank correlation is: 48.981% across all parts and all motifs (49.706% for the 1^st^, 56% for the 2^nd^, and 82.576% for the 3^rd^ parts). This low rate of significant configurations and the overall low correlation indicates that most of the activity induced by the stimulation spreads over the network on short and long ranges in a specific ways. The “direct network response” to stimulation seems to be captured by the 2^nd^ and 3^rd^ parts of the motifs because correlations are generally stronger for higher-order components, that is, 2^nd^ and 3^rd^ parts than the 1^st^ parts (and across the entire motifs). The first parts of each motif reflect the reorganization of the network activity caused by the stimulation. Because of the reorganization, these spatial formations of brain activity are however not necessarily traceable to the site, at which energy was introduced by stimulation.

When considering stimulation-induced activities in the entire brain model (i.e., including all substructure), the maximum correlation is the lowest but the number of parameter configurations with significant correlations is the highest. This emphasizes the specificity of responses to substructures of the brain (e.g., thalamus, isocortex). For instance, the activity in the cerebellum shows the highest maximum correlation with the direct projections from the stimulation sites, whereas the number of configurations with significant correlation is fairly low. It is also interesting to mention that activity in the isocortical subplate shows no significant correlation with the underlying structure at all.

#### Comparison of dynamically responsive and functional networks

We compared the DRNs with distinct functional networks. The spatial organization of the nine experimentally known functional networks derived from Sforazzini et al., (2014) and Zingg et al., (2016) is shown in Figure 5, panel **B** for the isocortex. Four of the nine functional networks are correlated with the five patterns occurring in the parts of the motifs (see Figure 5). Interestingly, the sensory motor networks are not reflected in the motifs of the DRNs. Merely the network associated with the frontal eye field shows significant but moderate correlations with four parts of four motifs (see magenta pattern in Figure 5). Here, the lateral activity of the frontal eye field network (i.e., activity within one hemisphere) corresponds to pattern 4 (see Figure 5). The unique parts in motifs 2, 4, 6, 7, and 9 (see parts in Panel **A** that are not highlighted) are not correlated with one of the nine specific functional networks. Moreover, pattern 1 (highlighted in yellow) in motif 12 does not correspond to any of the nine functional networks. The default mode and the salience networks show similarities at the isocortical level. Both comprised most of the motifs (cyan pattern 2 in motifs 1 to 11). Stimulation responses do activate to a lesser extent the lateral network (orange pattern 4 in motifs 1, and 3 to 5) and the medial network (gray pattern 3 in motifs 8 to 12), where the activity of both networks within one hemisphere (lateralized organization) is reflected. The default mode network (together with the salience network) is thus related in the model to the most flexible and sensitive pattern (cyan pattern number 2) that can activate and is involved in most of the motifs (1 to 11).

#### Organization of brain activity following sensory pathways

Stimulation of a brain area induces activity and traces out a spatiotemporal trajectory while dissipating energy into specific parts of the network. The trajectory is characteristic and leaves a particular fingerprint of responses. With our large-scale mouse brain network model we have for the first time the chance to systematically investigate sensory pathways and the processing of sensory data. We focus on five systems, which are: the auditory pathways, the visual pathways, the whisker pathways and the pathways for the limbs. For each of the sensory system we extracted network descriptions according to the scientific consensus from the literature (see Materials and Methods). Figure 6 shows the networks of the sensory pathways. We then tested whether the experimentally derived large-scale brain network contains the textbook descriptions of each of these sensory pathway networks. The sensory networks (visual, auditory pathways and the pathways for the whisker, as well as fore and hind limb) are statistically well represented in the used ABA connectome. The sensory pathways in Figure 6 suggest that sensory processing follows specific paths of activations towards the primary somatosensory cortices. Activation of sensory processing can be ipsilateral to the sensory organ, such as for the whisker follicles up to the level of the midbrain (see Figure 6, panel **D**). The sensory processing usually crosses to the contralateral side of the sensory organ, indicating the integration of the left and right sensory organ of the same modality (see Figure 6), such as for the auditory pathways showing directed processing to the contralateral side (projections from the medulla to the pons) and then a processing mainly in the contralateral side through the midbrain and the thalamus towards the primary somatosensory areas in the isocortex (see Figure 6, panel **B**). Because our systematic exploration of brain dynamics via stimulation provides the brain responses to focal stimulation, we can investigate the brain responses and the DRNs to stimulation of areas involved in sensory processing. We can thus trace the activity and the dynamics in each of the sensory pathways. Note that such an investigation is possible in silico but hard to accomplish for an individual brain in vivo. The network model extracted from the ABA significantly contains the sensory pathways. The sensory pathways as shown in Figure 6 do reflect the primary paths of sensory processing and are thus a detail of a bigger picture, that is, the large-scale brain network that spans the entire brain. The brain areas that are known to be involved in sensory processing may thus be connected to other brain areas and within the large-scale brain network and consequently involved in one or several functional networks. For this reason, it is evident that the entire brain dynamics needs to be considered in the sensory pathways. The brain areas in the sensory pathways as shown in Figure 6 provide guidance and prior knowledge for which the brain response to stimulation of exactly those areas and the occurring DRNs should be consistent and evolve. However, due to the large-scale network of the brain and its organization, it is not trivial that the dynamics in the model are consistent across a sensory pathway of a specific modality (e.g., auditory system). Because the statistics on the DRNs, see section ‘Statistics on DRNs’ in the Results, indicate that the response are (although weak) best predicted by the strengths of the directly outgoing connections from the stimulation site, we analyzed the spatial spread of the energy in the brain responses (integral of the power over time, that is, .2s to .8 s after stimulation). To test whether activity is focal or spread over the brain (or within a structure), we determined the brain areas that contain more than 95% of the total energy in a brain response (entire brain and within a brain structure). The results show that after stimulation of brain areas in the auditory pathways, the activity across all brain structures was most centered within the cerebellum in the flocculus and the copula pyramidis. In only one case (left and right stimulation of the nucleus of the brachium of the inferior colliculus of the midbrain) the activity was clearly centered in isocortex (laterointermediate area). The activity was mostly cumulated in the cerebellum. The distribution of activity was spreads and more complex in all other structures, namely olfactory areas, hippocampus, cortical subplate, cerebral nuclei, thalamus, hypothalamus, midbrain, pons, and medulla). In contrast to the auditory pathways, after stimulation of areas in the visual pathways the activity was most centered within the cerebellum but more moderate across stimulation sites. In only one case (left and right stimulation of the dorsal part of the lateral geniculate complex of the thalamus) the activity was clearly centered in isocortex around the laterointermediate area (better pronounced that for the case in the auditory pathway, that is, left and right stimulation of the nucleus of the brachium of the inferior colliculus of the midbrain). Activity after stimulation areas in the whisker pathways showed a clear centered distribution within the thalamus and within the cerebellum. The distribution of activity was more complex in all other structures. Furthermore, the results show that after stimulation of brain areas in the pathways of the limbs, the activity across all brain structures was less spread within the isocortex, the thalamus and the cerebellum. To clear whether activity spreads laterally we determined the brain areas that contain more than 95% of the total energy in a brain response in the entire brain and compared the location (left/right). The accumulation of activity is symmetric after stimulation of a given structure in the left and its complement in the right hemisphere. We found only weak asymmetries, which in turn point more towards a similar activation after left and after right stimulation. Most of the symmetric patterns are lateralized. Activity mostly cumulates either ipsilaterally or contralaterally to the stimulation site stimulation site (e.g., left or right LGN). However, non-lateralized symmetries are found in the medulla across all modalities. The pathways of the limbs show most non-lateralized symmetries across structures (olfactory areas, hippocampus, cerebral nuclei, pons, and medulla). Stimulation of areas in the auditory pathways indicates that most of the activity patters are lateralized and symmetric to left/right stimulation. The cerebral nucleus and the medulla show non-lateralized symmetries. Stimulation of areas in the visual pathways indicates that most of the activity patters are lateralized and symmetric to left/right stimulation. The medulla shows non-lateralized symmetries. Stimulation of areas in the pathways of the whiskers indicates that most of the activity patters are lateralized and symmetric to left/right stimulation. The pons and the medulla show non-lateralized symmetries. Stimulation of areas in the pathways of the limbs indicates that the activity patters are lateralized and symmetric to left/right stimulation. However, olfactory areas, hippocampus, cerebral nuclei, pons, and medulla show non-lateralized symmetries. To test whether the brain response to stimulation shows such symmetries as bilateral activations, we correlated the activity in the left areas with the activity of the corresponding areas on the right. A symmetric response means that the activity in the brain areas on the left side is similar to the ones on the right. An anti-symmetric response is indicated by a negative correlation and means that the activity on the left side is the reverse of that on the right. Not significant correlations (significance level of 5%) do not support either type of symmetry. The results suggest dividing the activity in the isocortex from the rest. Consistently across all stimulation of all brain areas involved in the sensory pathways (across all modalities and left and right areas), the responses in the subisocortex are mostly symmetric, whereas isocortical patterns are more variable. Dependent on the stimulation the response in the isocortex shows no symmetry (sign for lateralization), anti-symmetry (left/right inverted activity) or symmetries (bilaterally symmetric). This is consistent for the auditory, visual, whisker and limb pathways. In addition to investigating the distribution of the response we can analyze the energy to determine the areas that are most active. The energy in the brain responses (integral of the power over time, that is, .2s to .7 s after stimulation) indicates that the areas in the hypothalamus are the most active ones throughout stimulations, sites, and modalities. Their activation is lateralized and, in the 1st (dominant) part of a motif, contralateral to the stimulation sites in most of the cases (except vision, that is not lateralized). The most dominant areas in the hypothalamus are: the intermediate part of the periventricular hypothalamic nucleus, the lateral mammillary nucleus, and the arcuate hypothalamic nucleus. Ordering the areas to their strength of activity and then comparing between left structures and the complement on the right indicate a lateral symmetry. The brain areas that are most active after stimulation of areas across the auditory pathways are: the arcuate nucleus, the dorsal cochlear nucleus, the intermediate part of the periventricular nucleus, the abducens nucleus, the primary auditory area, and the laterointermediate area. The most active brain areas after stimulation across the visual pathways are: the laterointermediate area, the arcuate nucleus, and the intermediate part of the periventricular nucleus. Stimulation across the pathways of the whisker shows the following most active areas: the intermediate part of the periventricular nucleus, the primary somatosensory area of the mouth, the arcuate nucleus, the supplemental somatosensory area, and the ectorhinal area. The brain areas that are most active after stimulation of areas across the pathways of the limbs are: the intermediate part of the periventricular nucleus, the primary somatosensory area of the lower limb, the abducens nucleus, the gracile nucleus, the nucleus ambiguus, the laterointermediate area, the visceral area, and the lateral mammillary nucleus.

**Figure 6.**
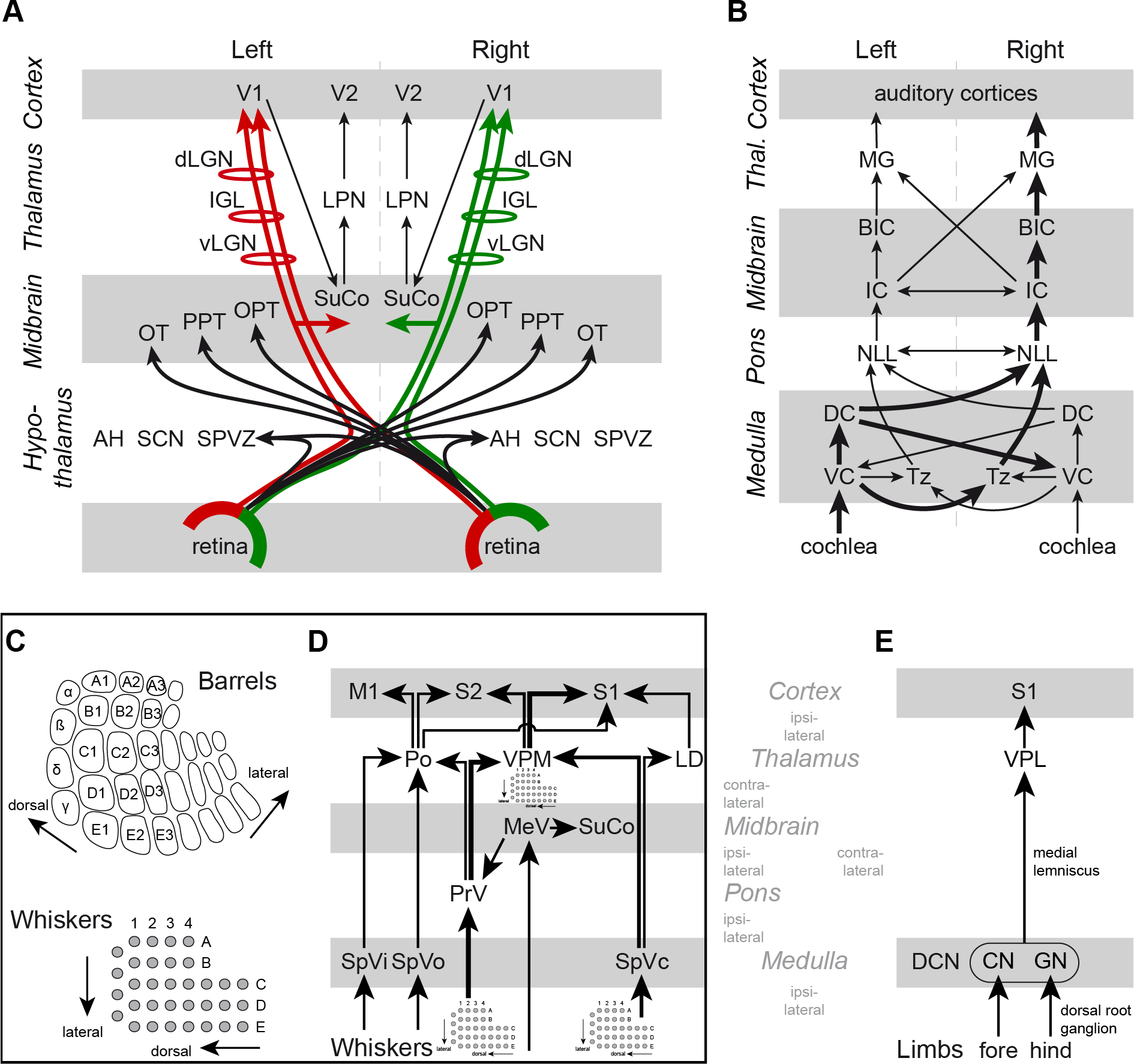
Networks of sensory pathways. Panel **A**: The visual pathways run from the retina through the hypothalamus, midbrain, and thalamus to terminate in the primary visual cortices. The retinal ganglion cells bilaterally connect to the hypothalamus and all of them project to the SuCo according to the ipsilateral visual field (colored links). The SuCo projects to the to areas of the cerebral cortex that are involved in controlling eye movements through the LPN of the thalamus. Note that the SuCo also receives activity due to whisker movements. The retinal ganglion cells have ramifications to contralateral structures in the midbrain (OT, PPT, OPT) and ramifications that run into the primary visual cortex through the thalamus (vLGN, IGL, dLGN) according to the ipsilateral visual field. The projections from the visuals fields through the retinal ganglion cells are highlighted in panel **A** by red and green colored links. The arrowheads indicate the termination areas of the axonal fibers. Panel **B**: The auditory pathways originate from the cochleae through the medulla, pons, midbrain, and thalamus to the primary auditory cortices. The thickness of the links in panel **B** indicates the strength of connectivity. Panels **C**–**D**: The pathways from the whiskers to the barrel fields run trough the medulla, pons, midbrain, and thalamus to terminate in the isocortex. Panel **C**: The somatotopic map of the whiskers where each individual whisker is represented in a discrete anatomical unit, that is, a barrel. The map shows the correspondence of the topographical organization of the major facial whiskers through the trigeminal nerve in the nuclei in the medulla (SpVc), the pons (barrelets), and through the thalamus (barreloids in VPM) to the somatosensory cortices (barrel cortex). Panel **D**: The pathways from the facial whiskers to the isocortex. The somatotopy of the whisker follicles mainly comes to the somatosensory cortices (i.e., S1, S2) via the trigeminal nerve nuclei (PrV in the pons and SpVc in the medulla) and the thalamus (VPM). The thickness of the links in panel **D** indicates the strength of connectivity. Whisker movements activate the SuCo in the midbrain through the MeV. The SuCo is also related to the visual system (see panel **A**) and in particular to eye movements. Panel **E**: The pathways from the fore and hind limbs pass via dorsal root ganglion to the ipsilateral dorsal column nuclei (DCN) in the medulla (CN and GN). The upper limb, and especially the forelimb, connects through the CN whereas the lower limb, and especially the hind limb, connects through the GN in the medulla. The nuclei of the medulla connect via the medial lemniscus to the contralateral thalamus (VPL), which has ipsilateral ramifications into the primary somatosensory cortex (S1). The sensory networks in panels **A**, **B**, **D**, **E** are based on textbook descriptions (e.g., Watson et al., 2011) and include the relevant structures given by the ABA. The sensory pathways can be described using the ABA. However, the ABA needs refinements to distinguish lower and upper limb (especially hind and forelimb), see panel **E**. The ABA does not include a division of the barrel field of the primary somatosensory areas, see panels **C**–**D**. The networks in panels **A**, **B**, **D**, **E** indicate information flows to areas that do not necessarily terminate in the primary sensory areas of the isocortex, such as the hypothalamic and midbrain targets of the retinal ganglion cells in panel **A,** and the midbrain nuclei related to whisker movements in panel **D**. These connections are well known and usually not discussed regarding sensory processing in textbooks. As a result, visual and whisker system meet in the SuCo regarding eye and whisker movements. Note that the model is agnostic about the (sensory) information, meaning that nuclei may be activated but the object of processing (information) is less defined. Meaning is attached by the vast amount of studies about the physiology of the nuclei. For instance, the hypothalamic nuclei play a role in the circadian timing system, and the nuclei in the midbrain (OT, PPT, OPT) receive input from the retina (see panel **A**) and are known to be involved in eye movement coordination and reflexes. The ABA includes the following areas involved in sensory processing. The nuclei in the medulla are DC: dorsal cochlear nucleus, CN: cuneate nucleus, GN: gracile nucleus, SpVc: caudal part of the spinal nucleus of the trigeminal, SpVi: interpolar part of the spinal nucleus of the trigeminal, SpVo: oral part of the spinal nucleus of the trigeminal, Tz: nucleus of the trapezoid body, and VC: ventral cochlear nucleus. The hypothalamic nuclei are AH: anterior hypothalamic nucleus, SCN: suprachiasmatic nucleus, and SPVZ: subparaventricular zone. Involved structures in the pons are NLL: nucleus of the lateral lemniscus, and PrV: principal sensory nucleus of the trigeminal. The sensory related nuclei of the midbrain are BIC: nucleus of the brachium of the inferior colliculus, IC: inferior colliculus, MeV: trigeminal nucleus, OT: nucleus of the optic tract, OPT: olivary pretectal nucleus, PPT: posterior pretectal nucleus, SuCo: superior colliculus, sensory related. The thalamic nuclei involved in the sensory pathways are dLGN: dorsal part of the lateral geniculate complex, IGL: intergeniculate leaflet of the lateral geniculate complex, LD: lateral dorsal nucleus, LPN: lateral posterior nucleus, MG: medial geniculate complex, Po: posterior complex, vLGN: ventral part of the lateral geniculate complex, VPL: ventral posterolateral nucleus, and VPM: ventral posteromedial nucleus. The sensory brain areas in the isocortex are A1: primary auditory area, V1: Primary visual area, V2: secondary visual area, S1 primary somatosensory cortex, S2: secondary somatosensory cortex, and M1: primary motor cortex. Note that the brain areas are listed in Table 2 and Table 3.

So far we have analyzed the brain response to stimulation of brain areas in sensory pathways independently. To clarify whether the brain responses are consistent for a certain sensory pathway summarized in Figure 6, for example, the auditory pathways from the cochlea to the primary auditory areas, we made use of the correlation of the responses and the clustering into characteristic motifs of the dynamically responsive networks (DRNs) as described in section ‘Statistics on DRNs’ in the Results. The catalogue of dynamically responsive networks and its stimulation sites (see Figure 4 and Figure 5) reveal whether the responses due to stimulation along the pathways belong to the same cluster. The catalogue shows that the responses are consistent and alter along the auditory pathways (bottom-up). The formation from the cochlea to the primary auditory areas can be subdivided into three motifs (from cochlea to cortex through medulla, pons, midbrain, and thalamus): (1) the motif with number 5 in Figure 5 due to stimulation in the medulla (dorsal and ventral cochlear nucleus), (2) the motif with number 9 in Figure 5 shared by stimulation of the nucleus of the trapezoid body of the medulla, the nucleus of the lateral lemniscus of the pons, and the nucleus of the brachium of the inferior colliculus of the midbrain, and (3) the motif with number 10 in Figure 5 shared by stimulation of the inferior colliculus (midbrain) and the medial geniculate complex of the thalamus. The catalogue shows that the responses are consistent throughout the visual pathways and have the same motif, that is, motif with number 10 in Figure 5. The catalogue shows that the responses are consistent and alter along the pathways of the whiskers (bottom-up). The formation from the whisker follicles to the barrel field of the primary somatosensory areas can be subdivided into three motifs (from follicles to cortex through medulla, pons, midbrain, and thalamus): (1) motif with number 6 in Figure 5 shared by caudal, interpolar, and oral part of the spinal nucleus of the trigeminal in the medulla as well as by the principal sensory nucleus of the trigeminal in the pons, (2) motif with number 6 in Figure 5 shared by stimulation in the midbrain (of the superior colliculus, sensory related) and in the thalamus (of the lateral dorsal nucleus), (3) the two thalamic areas, that is, the ventral posteromedial nucleus and the posterior complex of the thalamus share the same motif that has the number 8 in Figure 5. The response to stimulation of the nuclei of the medulla involved in the sensory pathways of the limbs, that is, cuneate nucleus and gracile nucleus are correlated and have the same motif (number 6 in Figure 5). The responses due to stimulation of the nuclei in the thalamus, that is, the ventral posterolateral nucleus of the thalamus, and the parvicellular part of the ventral posteromedial nucleus of the thalamus differ and belong to two motifs with the number 8 and 7 (for the parvicellular part) in Figure 5.

#### Comparison of DRNs to in-vivo brain dynamics after stimulation

To compare the simulated and empirical spatiotemporal dynamics of the brain activity after focal stimulation of the isocortex, we used wide-field voltage-sensitive dye (VSD) imaging combined with optogenetic focal stimulation of the cortical areas (Figure 7, panel **A**). This technique allowed us to capture the brain activity over a large portion of the cortical mantle (Figure 7, panel **B**) while an arbitrary point on the cortical surface was stimulated (Figure 7, panel **C**). We qualitatively observe a large similarity between simulated and empirical patterns of brain activity evoked by the focal stimulation of the retrosplenial (RSC) and visual areas (see Panels **A** in Figure 8 to Figure 10). This similarity was described by the overlap of the maximally activated patterns of cortical regions after focal stimulation (see Panels **B** in Figure 8 to Figure 10). The spatial organization of these patterns is not trivially linked to the stimulation focus and shows characteristic and reproducible large-scale activations across the measured spatial window. Both empirical and simulated data suggested that the brain activity evoked by the focal stimulation could be predicted by the structural connectivity between regions. For example, focal stimulation of the RSC led to the activation of the visual and midline areas, which are strongly connected to RSC. However, our empirical and simulated data did not agree when we compared the simulated and empirical spatiotemporal patters of brain activity evoked by the stimulation of sensory areas other than visual ones. Here we did not observe a great amount of overlap in the set of activated cortical regions after focal stimulation of such areas.

**Figure 7.**
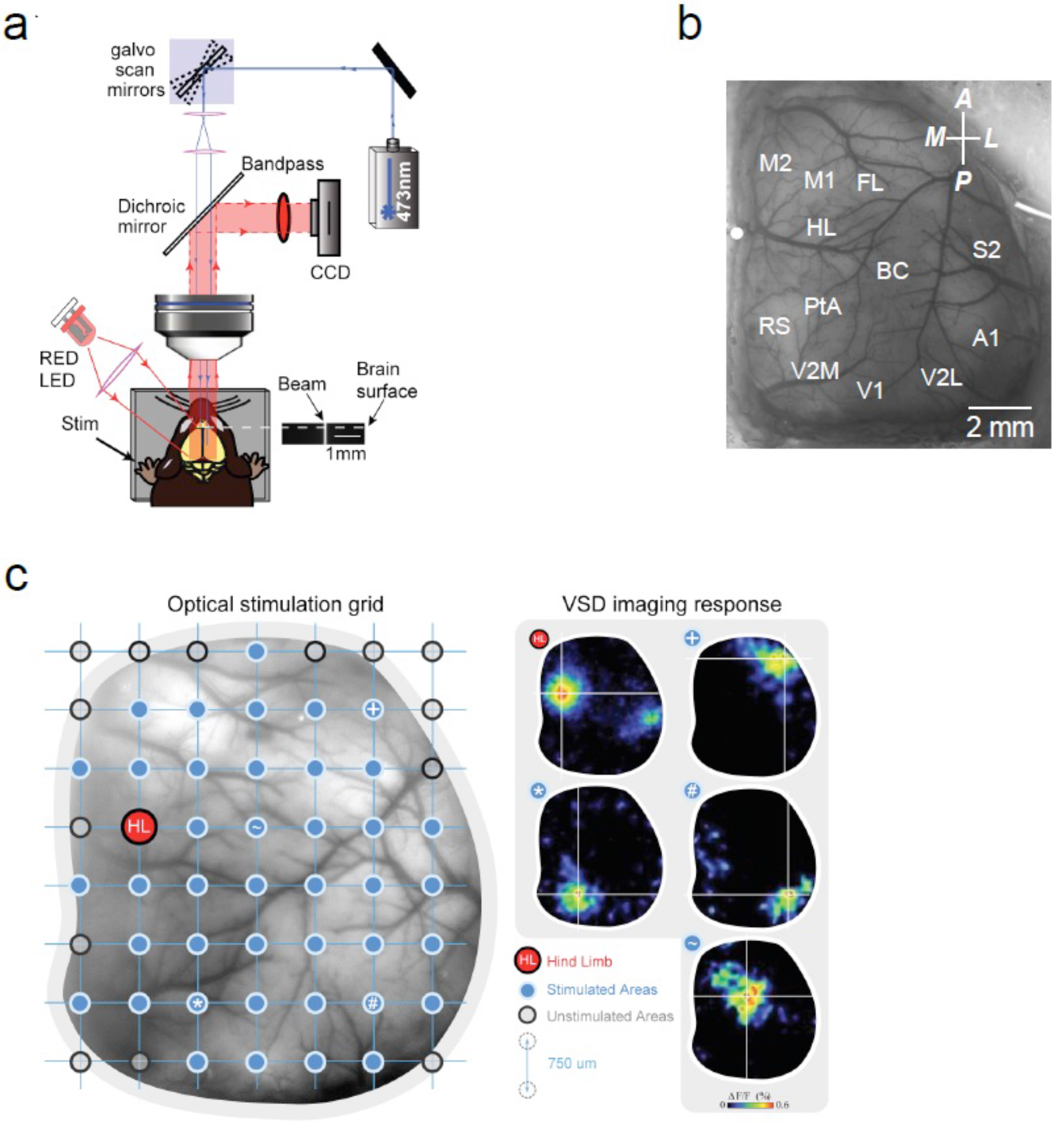
Experimental apparatus and method to investigate brain response to stimulation. Panel **A**: The schematic of the experimental set-up. This set-up consists of a 473 nm laser generator whose beam is directed by galvo scan mirrors to a specific point on the cortical surface. By changing the configuration of the galvo scan mirrors, we can change the coordinates of the stimulated point. The CCD camera captures the voltage activity of the brain. Panel **B**: An example of a cranial window over the mice cortex prepared for VSD imaging. The cortical regions are labeled based on their coordinates with respect to the bregma, marked by white circle. Panel **C**: An example of the grid points mapped on the cortical surface by galvo scans mirrors. The laser can stimulate each one of these points. Examples of the voltage activity right after stimulation of some of these points are presented on right.

**Figure 8.**
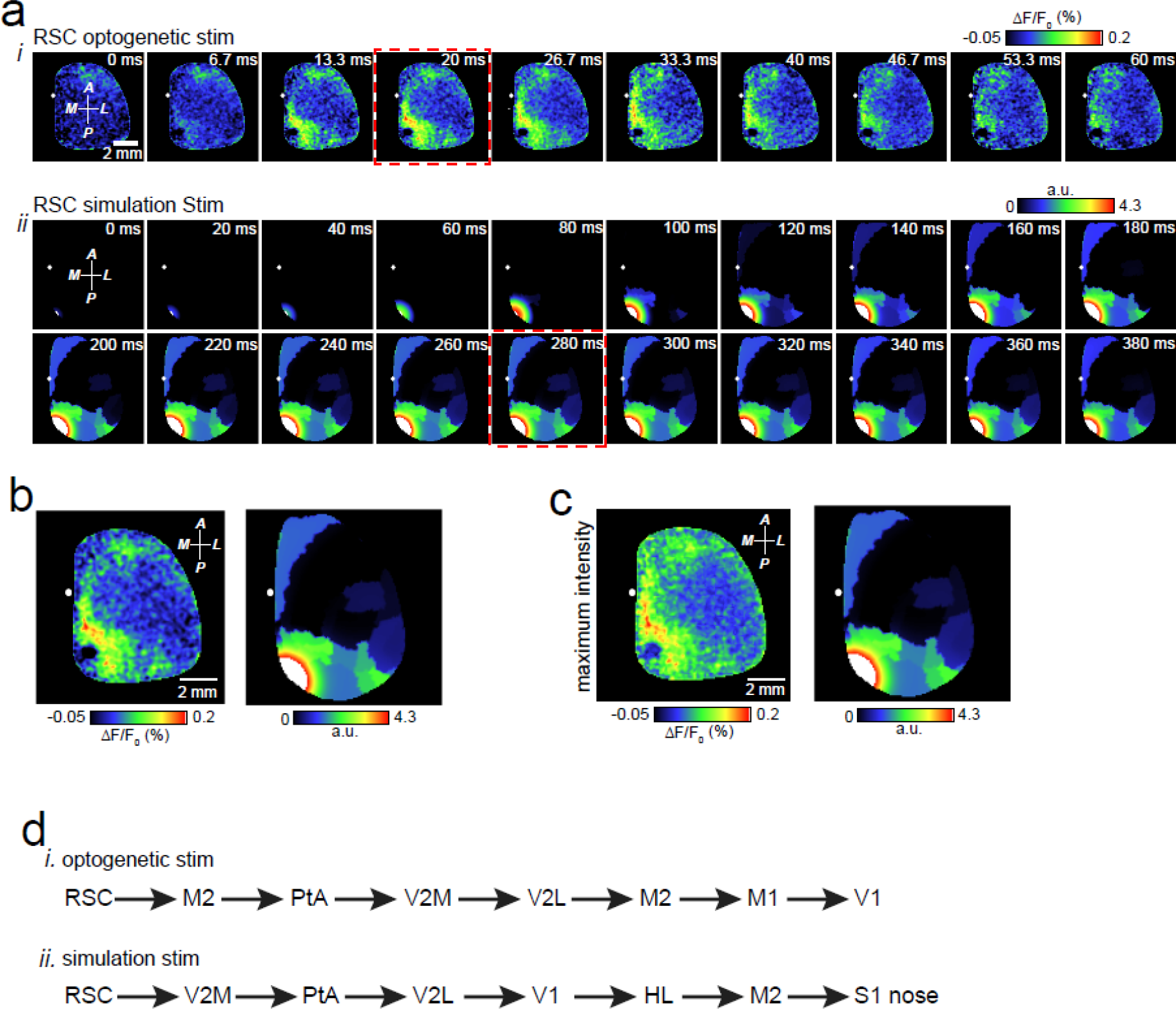
Brain response after stimulation. Panel **A**: The spatiotemporal pattern of voltage activity after (i) optogenetic and (ii) simulated stimulation of RSC. Panel **B**: The enlarged frames, highlighted by a red rectangle in Panel **A**, juxtaposed for easier comparison. Panel **C**: The spatial distribution of the post-stimulus maximum voltage activity Panel **D**: The temporal order of activation of cortical regions after optogenetic (i) and simulated (ii) stimulation. Activation occurs when the voltage level of a given region surpasses 20% of its peak activity after stimulation was applied.

**Figure 9.**
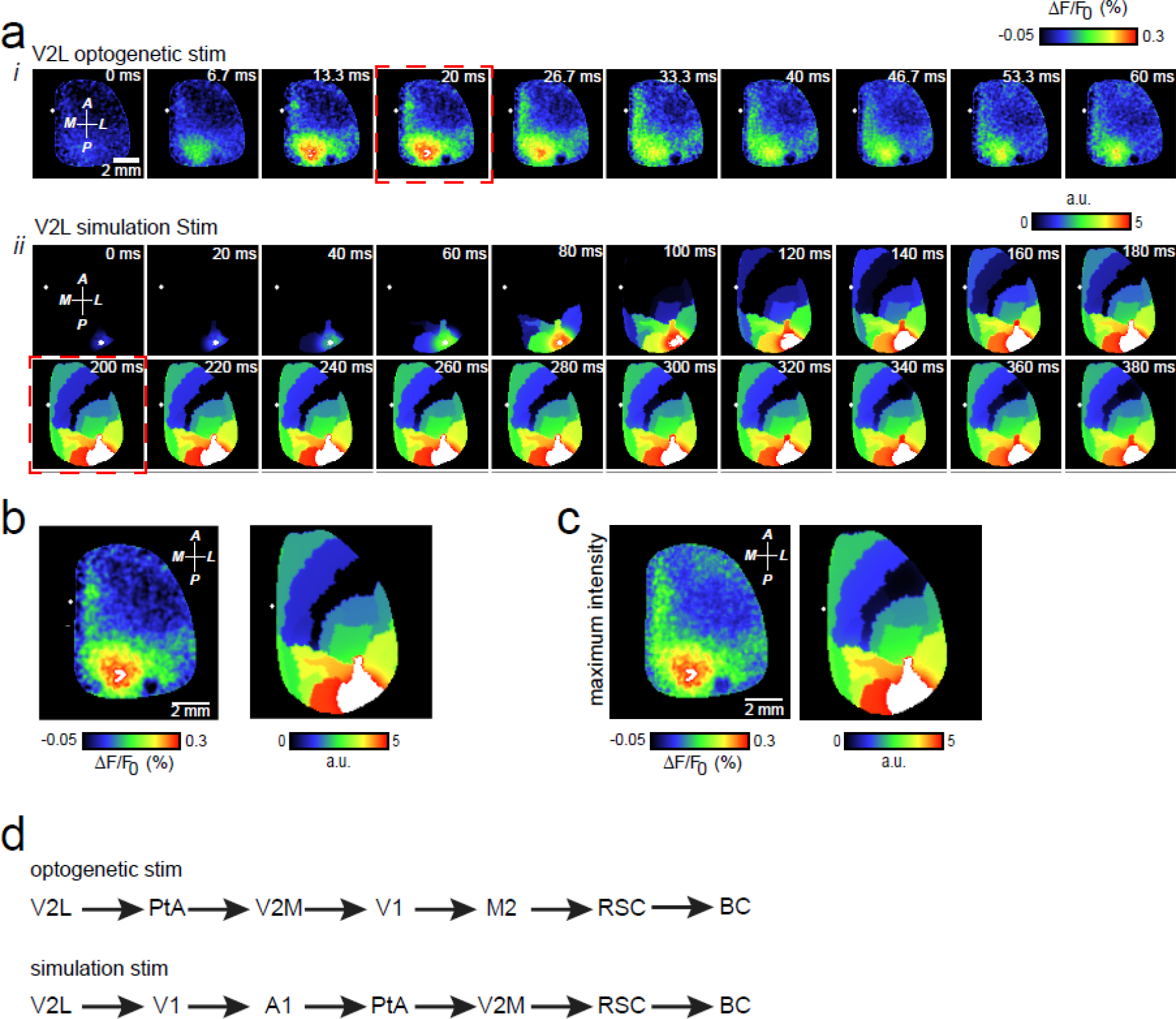
Panel **A**: The spatiotemporal pattern of voltage activity after optogenetic (i) and simulated (ii) stimulation of V2L. Panel **B**: The enlarged frames, specified by a red rectangle in Panel **A**, juxtaposed for easier comparison. Panel **C**: The spatial distribution of the post-stimulus maximum voltage activity. Panel **D**: The temporal order of activation of cortical regions after optogenetic. (i) and simulated (ii) stimulation. Activation occurs when the voltage level of a given region surpasses 20% of its peak activity after stimulation was applied.

**Figure 10.**
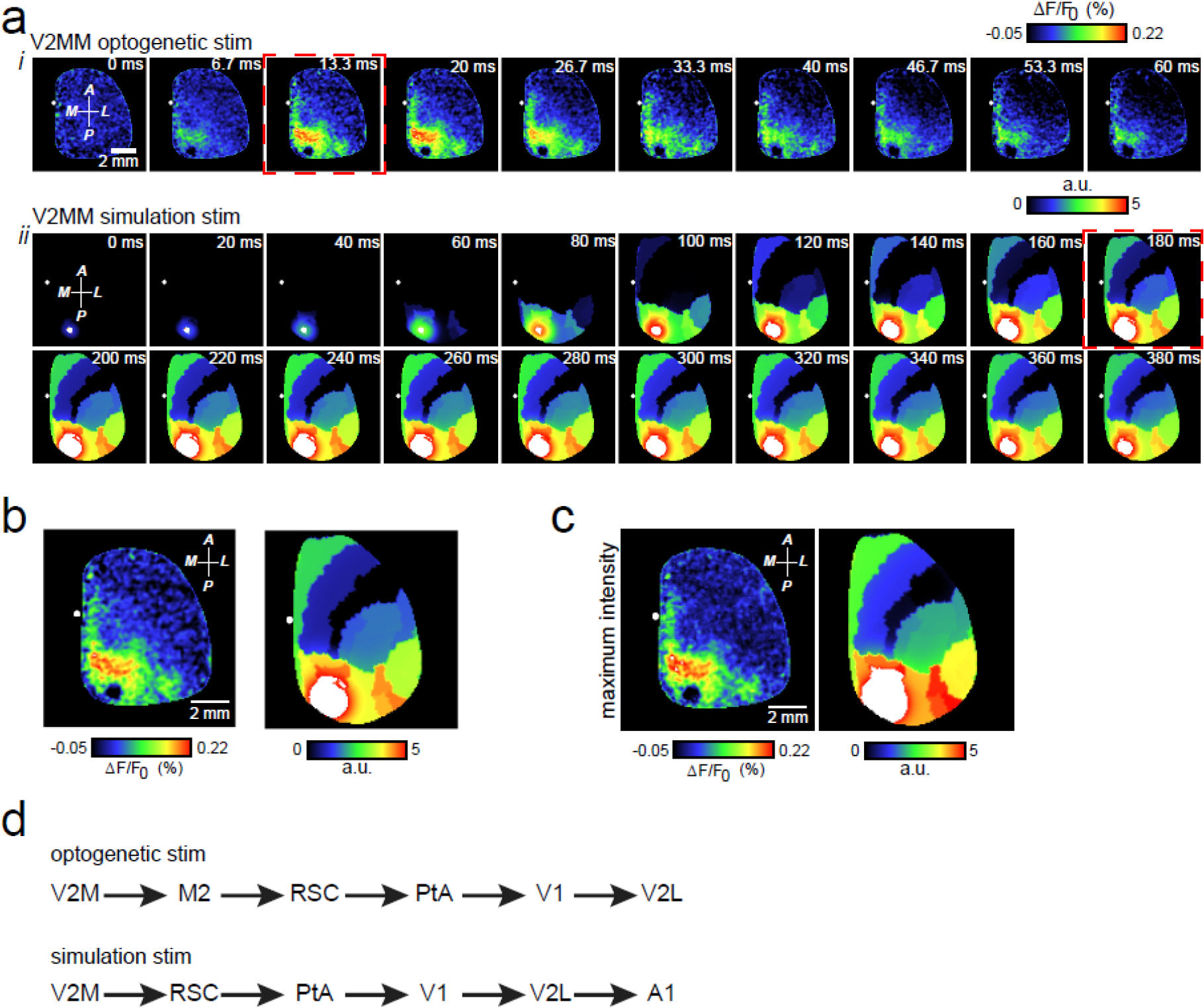
Panel **A**: The spatiotemporal pattern of voltage activity after optogenetic (i) and simulated (ii) stimulation of V2M. Panel **B**: The enlarged frames, specified by a red rectangle in Panel **A**, juxtaposed for easier comparison. Panel **C**: The spatial distribution of the post-stimulus maximum voltage activity. Panel **D**: The temporal order of activation of cortical regions after optogenetic (i) and simulated (ii) stimulation. Activation occurs when the voltage level of a given region surpasses 20% of its peak activity after stimulation was applied.

Moreover, the temporal order of activation cortical regions after focal stimulation showed some level of similarity (see Panels **D** in Figure 8 to Figure 10, and Supplementary Figure 1 to Supplementary Figure 3). We observed that the temporal order of activation in the simulated data was preserved up to one or two transpositions compared to the empirical data. It suggests that our model assumptions can fairly explain the flow of information in the cortical networks.

### Modeling the mouse brain

#### Structure and division of the mouse brain model

The model is based on the Allen Brain Atlas (ABA), which compiles tracer injections, selectively sampling mice brain volumes, to map axonal projections of different areas within the adult mouse brain (see Figure 1). The database of the Allen Brain Atlas was spatially sampled with a resolution of 50 μm (voxel of 50 (μm)^3^ isotopically sample the volume). In this way, the model distinguishes between 512 distinct brain areas. The smooth surface of the isocortex was reconstructed from the ABA, approximated by a regular mesh of 27,554 triangles with 41,524 edges (88 μm is the length of the shortest edge, 190 μm is the longest edge, with a mean edge length of 125 μm) between 13,972 vertices (Figure 11), and spatially divided into 84 cortical areas given by the ABA. The division of the reconstructed isocortical surface into areas, that is, the parcellation was performed by (i) extending the normal vectors of each vertex to about 200 μm inside the volume of the isocortex, (ii) determining the 5 closest injection sites, and (iii) evaluating their classification to an area of the isocortex in the ABA. The isocortical parcellation was furthermore corrected for holes and isolated/mismatched vertices. The 512 distinct brain areas contain the 84 isocortical areas (42 per hemisphere) and 428 (non-isocortex) areas composing the olfactory areas, the hippocampal formation, the cortical subplate, the cerebral nuclei, the thalamus, the hypothalamus, the cerebellum as well as nuclei in the midbrain, the pons, and the medulla (see Table 1 for the division, Table 2 and Table 3 for a complete list of area names). These 428 sub-isocortical areas are lumped in the model to a point in physical space (i.e., centroid).

**Figure 11.**
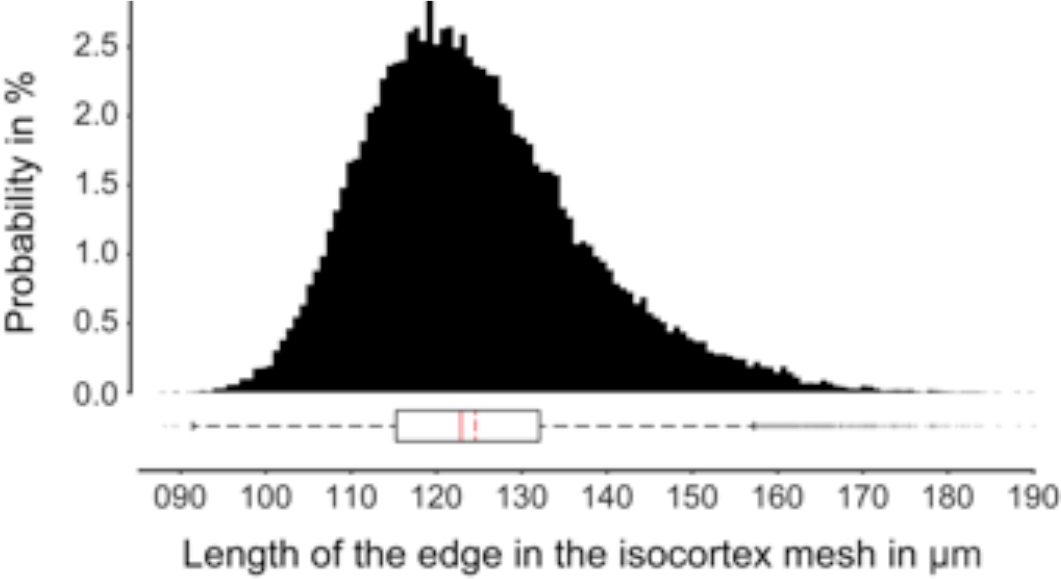
The sharp unimodal distribution of edge lengths indicates the quality of the reconstructed surface of the isocortex using a regular-triangle mesh of 27,554 triangles with 41,524 edges. The expectation and the standard deviation is (124.5786 ± 13.0144) μm. The edge lengths are positively skewed (0.7164) due to positive definite length (minimum and maximum lengths are 88.1525 μm and 190.0629 μm).

#### Short-range structural connectivity

The short-range connectivity comprises axonal projections within the isocortex (e.g., in a brain region) over distances of μm up to mm on the surface (Braitenberg and Schüz, 1998). These short-range connections were assumed to be homogeneous throughout the isocortex, which connectivity decreasing with distance (Braitenberg and Schüz, 1991). In the model, the so-called homogeneous structural connectivity (SC) resembles short-range connections linking vertices of the isocortical mesh within an area, and between areas if they are spatially close to one another with a connection probability following a Gaussian function and decreasing with distance. Note that the homogeneous SC concerns the isocortex. Other brain areas are not directly linked via the homogeneous SC. Because the homogeneous SC represents an approximation of short-range connectivity, the spatial range of the Gaussian kernel was systematically varied by its standard deviation σ between 500 μm and 1000 μm in steps of 100 μm. The spatial cut-off range for short-range connections was eight times the standard deviation of the connectivity kernel 8σ (Spiegler and Jirsa, 2013). This gives six configurations of the homogeneous SC.

#### Long-range structural connectivity

The long-range connectivity comprises axonal projections of distant brain areas. This type of structural connections is estimated by the ABA between the 512 distinct brain areas from adult mouse brains. The weights of the connectivity are obtained by the ratio of the projection density to the injection site volume (Oh et al., 2014). Note that the ABA provides directional information connectivity in the connectivity. The connectivity of the lateral geniculate nucleus to the primary visual area, for instance, weighs more than the connectivity of the primary visual area to the lateral geniculate nucleus. In the model, the so-called heterogeneous structural connectivity (SC) resembles long-range connections linking the brain areas. The heterogeneous SC links all the vertices of an area with the vertices of another area. Note that the 428 non-isocortical areas represent a node in the large-scale brain network. Connections between these nodes are point-to-point connections. Connections between an area of the isocortex and on of the 428 non-isocortical areas are links between all vertices of the isocortical area and the node of the non-isocortical area. Note that within the isocortex, neighboring areas are able to exchange information via the homogeneous SC within the cortex and via the heterogeneous SC leaving and entering the isocortex. At each local node (brain area) activity arrives and exits via afferent and efferent projections with an experimentally derived strength that indicates how strong a node collects and distributes activity - in other words, how well node is embedded in the network. The graph-theoretic measures, that is, in-strength (afferent projections and arriving activity) and out-strength (efferent projections and exiting activity) are computed given the experimentally derived large-scale heterogeneous SC. In addition, the in-degree (number of afferent connections) and out-degree (number of efferent connections) are computed. Figure 12 and Figure 13 list the top 10 brain areas, indicating that the hypothalamus is the most embedded brain structure in the mouse brain model based on the ABA. The hypothalamus receives the strongest input in the mouse brain model and the cerebellum the weakest. The graph-theoretic measures also indicate that the subcortical structures are weakly connected with the isocortex tough ample connections exist. The graph-theoretic measures also indicate that the subcortical structures are on average weakly connected with the isocortex (see Figure 12) though ample connections exist. This does not necessarily preclude the sensitivity of the isocortex to subcortical input but rather emphasizes that the isocortex can be functionally autonomous as well under particular circumstances (certain brain states).

**Figure 12.**
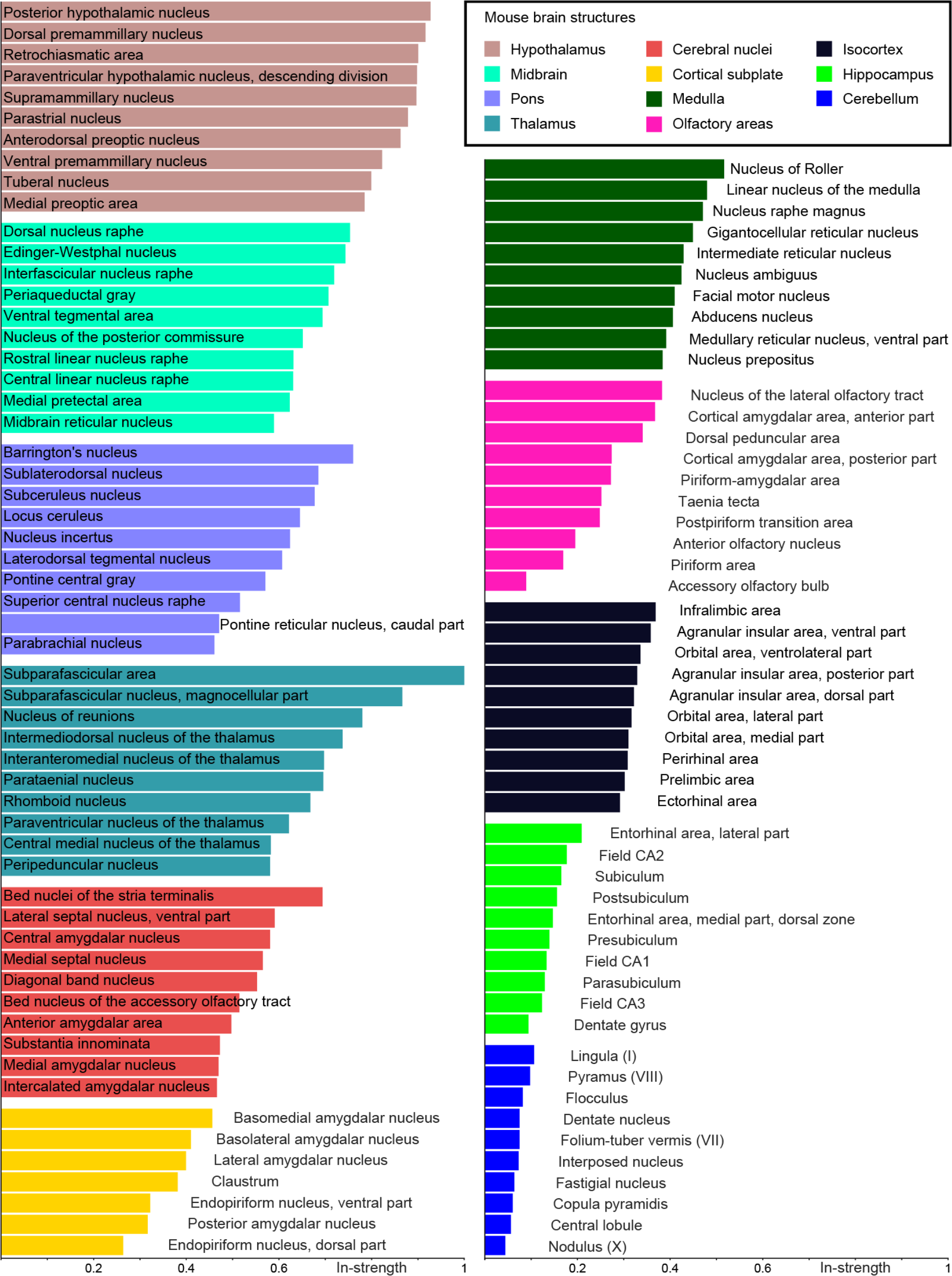
In-strength indicates the income of a brain area. The hypothalamus receives the strongest input in the mouse brain model and the cerebellum the weakest. The isocortex is listed 9 out of 11 structures. The structures are ordered by their mean in-strength. This indicates that the subcortical structures are highly interdependent and the isocortex might be sensitive to subcortical input but autonomous as well. For each structure (e.g., thalamus) 10 areas are listed in order of the highest in-strength. Note that the cortical subplate is divided in seven areas.

**Figure 13.**
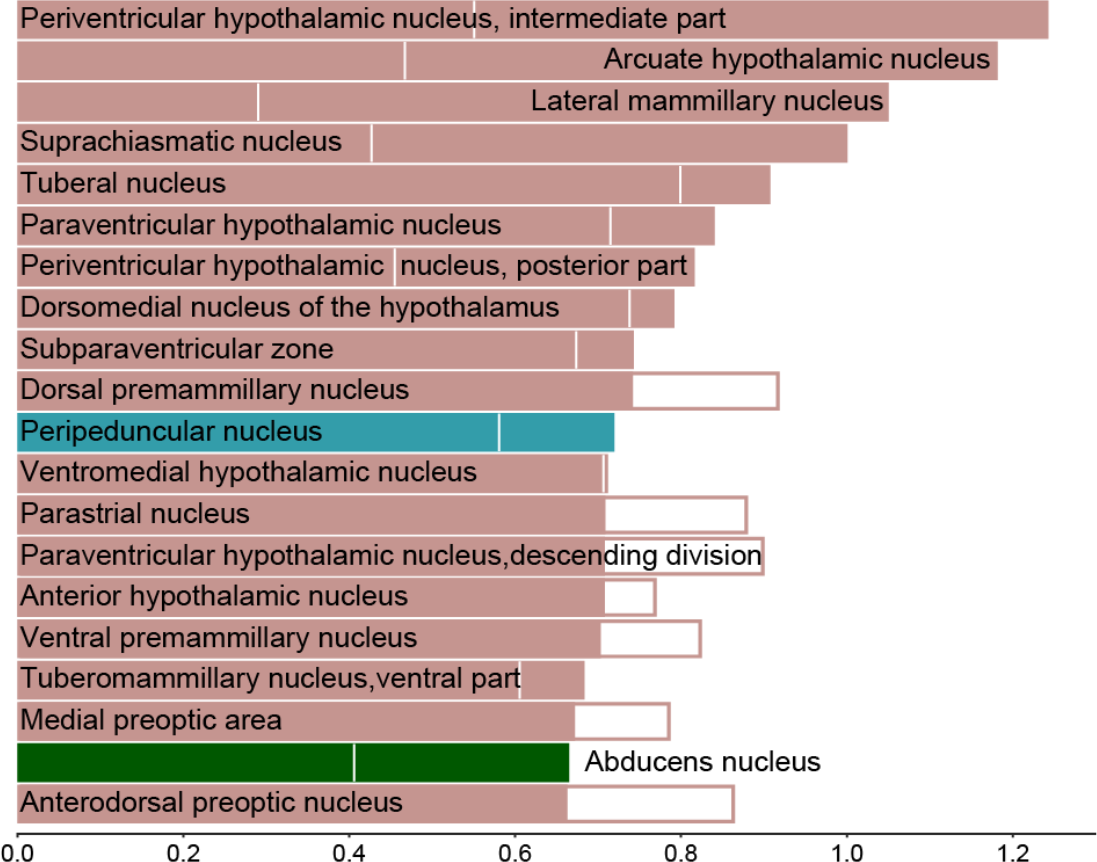
Out-strength indicates that areas in the hypothalamus project the most in the mouse brain model. The color-code for the structures is the same as in Figure 12: brown indicates areas in the hypothalamus, dark cyan the thalamus, and dark green medulla. The white vertical lines in a bar and the white filled bars indicate the in-strength of an area.

Though the tracer injection techniques underlying the ABA allow for directional information of connectivity, the analysis of the weight matrix of the ABA structural connectivity reveals a high symmetry, thus dominant bi-directionality of connectivity. Two measures were used for estimating the symmetry in the connectivity weights (see Materials and Methods). Both measures, *Q*_0_ and *Q*_1_ ranging between 0 (if symmetric) and 1 (if asymmetric and anti-symmetric respectively) and indicate a high symmetry in the long-range structural connectivity reconstructed from the ABA with *Q*_0_ = 238.2871 × 10^−3^ and *Q*_1_ = 226.6417 × 10^−3^. The tracer experiments compiled in the ABA mostly relate the spatial diffusion of the tracer to the injection site in the right brain. The connectivity is consequently left/right symmetric in the mouse model. For that reason, results are shown for one brain site though the model simulations where performed using both left and right hemispheres and the interhemispheric connections. For a sanity check, the simulated activity was correlated between the same area in the left and right brain and confirmed the left/right symmetries imposed by the area.

#### Conduction speed and transmission delays

The axonal conduction speed is determined by the axonal diameters as well as the degree of myelin. Both the axonal diameter and the degree of myelination vary throughout the mice brain (Wang et al., 2008; Foran and Peterson, 1992). Compared to larger mammalian brains as in humans, axons are less myelinated in the brains of mice, and if so, the axons are shorter (see, for example, Wang et al., 2008). The axonal conduction speed in mice is in the range of 0.2 m/s (Wang et al., 2008) for unmyelinated axons with an average axonal diameter in mice of 0.3 μm (Braitenberg and Schüz, 1998). The axonal conduction speed for myelinated axons is in range of 0.3 m/s and 3.5 m/s (Wang et al., 2008; Swadlow and Waxman, 2012; Salami et al., 2003). Length and the conduction speeds of axons cause time delays in the transmission of information processing. In contrast to human brains, the conduction via myelinated axons in mice is slower and more similar to unmyelinated axons (Wang et al., 2008). Moreover, the ratio of myelinated axons to unmyelinated axons is smaller in mice brains compared to human (Wang et al., 2008). Myelinated axons are shorter in mice brain compared to human (Wang et al., 2008). Note that the injection techniques underlying the ABA do not provide an estimate of the axonal lengths. The Euclidean distances between brain areas could be used as a proxy to estimate axonal lengths. Another possibility is given by the availability of diffusion tensor MRI data, from which the axonal tracts can be estimated and a connectivity matrix, similar to the ABA can be compiled. In contrast to the ABA, the diffusion tensor MRI however does not provide information about directionality but mouse individual connectivity information. With the availability of diffusion tensor MRI data of mice, the ABA connectivity could be co-register with the diffusion tensor connectivity, and thus provide information about axonal lengths. Following the reasoning of Buzáki and Wang (2012) the local brain activity, that is, of neural populations shows gamma oscillations (30-90 Hz) with cycle durations between 11 ms and 33 ms. Considering that the average length of axonal tracts between lateral areas of one isocortex and contralateral isocortical areas in mice ranges between 20 mm and 40 mm (Schüz et al., 2006) and assuming that the time scales of transmission delays have to be similar to the characteristic time scales of local brain activity, result in an effective conduction speed smaller than 3.6 m/s for the extreme, that is, the longest axon (40 mm) and the fastest local activity (11 ms). This estimation is in line with the range of conduction speed found in myelinated visual cortical neurons through the corpus callosum in mice (Swadlow & Waxman, 2012). Most of the cortico-cortical homolateral connections in mice are shorter than 10 mm (e.g., Braitenberg and Schüz, 1998, Chapter 26, report 7 mm for the longest distance to the tracer injection site in a planar view on the isocortices with two modes, at about 2 mm and at about 3.5 mm). Taking the same approach, this results in effective conduction speed values smaller than 1 m/s for a maximum length of 11 mm divided by 11 ms to affect fast (90 Hz) gamma oscillation (the upper bound of the speed for the longest distance to the tracer injection site and the two modes of distance to the tracer injection site, see, Braitenberg and Schüz 1998, Chapter 26, are 0.6364 m/s = 7 mm / 11 ms, 0.1818 m/s = 2 mm / 11 ms, and 0.31818 m/s = 3.5 mm / 11 ms). Transmission delays are assumed to hardly affect intrinsic local activity, that is, gamma waves, which are most apparent at a frequency of >40 Hz across the brain. If we assume a modest spread of 3 m/s and 10 mm length, then this results in time delays of up to 3.333 ms. This is short and may not play a role in the mouse, but remains limit, especially if the speed can be slowed down by a fact of 2-3. Considering harmonic oscillations in the brain, the time delay can be effective on a scale smaller than the characteristic time constant of the intrinsic frequency (that is the period of a cycle) and expected to be most effective around a quarter of a full cycle (and multiple of half a cycle) because of the gradients, which results in effective time delays around 6.25 ms for 40 Hz and 2.77 ms for 90 Hz. This means that a conduction speed of about 1 m/s is effective for 40 Hz oscillations if the majority of connections is about 6.25 mm long (and 2.77 mm for 90 Hz). Short time delays affect mainly fast oscillations but not necessarily slow oscillations and long time delays affect slow oscillations more than fast oscillations. In this sense, we have provided each network node with simple dynamics that allows it to respond to stimulation and incoming input from other brain regions with a damped oscillation in the gamma range (at about 42 Hz), see Figure 2.

The conduction speed for white-matter fiber tracts is assumed in the model to be 1 m/s. Given the distances between the brain areas from the ABA, the transmission delays result from the division of the mean connection lengths by the conduction speed (see Figure 14). First of all, the histograms in Figure 14 do not give indications for bimodality as reported by Braitenberg and Schüz, 1998 in Fig.62, Chapter 26, though the technique is similar and the distance measurement is slightly different. However, after dividing the entire mouse brain into areas (e.g., left/right isocortex) and subareas (e.g., left/right barrel cortex; following the ontology of ABA) and investigating intra-hemispheric connections (e.g., among left isocortical areas) and inter-hemispheric connections (e.g., between left and right isocortical areas) we can confirm the second peak in the histogram whose significance is obscure (see Figure 14 panel **B**). This second peak basically reflects the long commissure connecting the two hemispheres such as the corpus callosum. Because the number of connections within the isocortex is similar to those between left and right isocortical areas (Table 4), the effect of the two peaks in the histogram for intra-and inter-hemispheric connection lengths is not visible in the histogram of all isocortical connection lengths. The same is true for all the connections in the entire mouse brain model (see Figure 14 panel **A**). The descriptive statistics of the connection lengths are summarized in Table 5.

**Table 4.** The number of connections among areas in the isocortex is outnumbered by other areas in the mouse brain model. The table lists the number of connections of the mouse brain model and its division into isocortex and sub-isocortex. The number of different areas is given in brackets.

| Area | Structure |  |  |
| --- | --- | --- | --- |
|  | Mouse brain (512) | Isocortex (84) | Sub-Isocortex (428) |
| Intra | 13,0560 | 003,444 | 091,592 |
| Inter | 13,1072 | 003,528 | 091,164 |
| All | 26,1632 | 006,972 | 182,756 |

**Table 5.** Statistics of connection lengths in the mouse brain model and its subdivision. Note the difference between intra-and inter-iscocortical connections in terms of expectation, median (i.e., 50% quantile), and variance. All values are in mm, except the variance that is in mm^2^ and the skewness that is a bare number.

| Parameter | Mouse brain |  | Isocortex |  | Sub-isocortex |  |
| --- | --- | --- | --- | --- | --- | --- |
|  | Intra | Inter | Intra | Inter | Intra | Inter |
| Minimum | 0.1085 | 0.1083 | 0.4441 | 0.2352 | 0.1085 | 0.1083 |
| Maximum | 11.6731 | 11.8715 | 8.5598 | 10.0594 | 11.6731 | 11.8715 |
| Expectation | 4.0562 | 4.9721 | 3.7661 | 6.3171 | 3.6806 | 4.4273 |
| Variance | 3.3677 | 3.4961 | 2.8349 | 3.5418 | 3.0860 | 2.9773 |
| Skewness | 0.5133 | 0.2307 | 0.2042 | -0.5012 | 0.6632 | 0.3889 |
| Quantile-05% | 1.3444 | 2.0139 | 1.1605 | 2.7377 | 1.1909 | 1.7948 |
| Quantile-50% | 3.8858 | 4.9148 | 3.6685 | 6.5294 | 3.4521 | 4.3090 |
| Quantile-95% | 7.2747 | 8.1711 | 6.5422 | 9.0240 | 6.8353 | 7.3476 |
| Quantile-99% | 9.0197 | 9.4866 | 7.7075 | 9.5383 | 8.4382 | 8.7810 |

**Figure 14.**
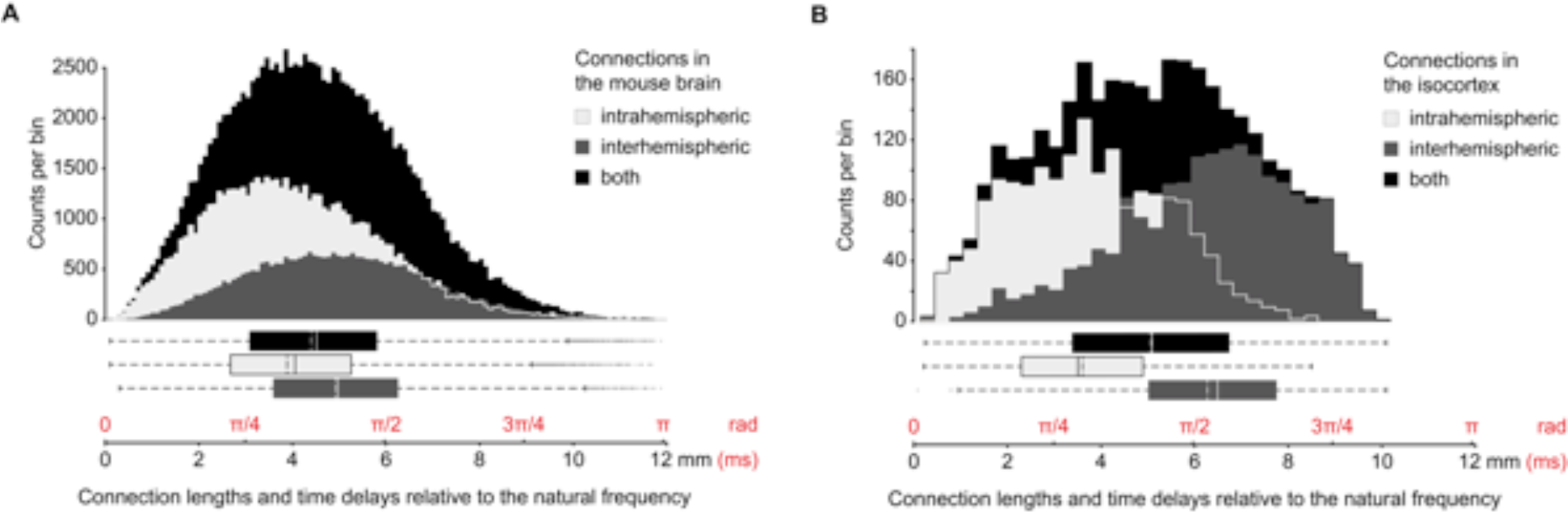
Histograms of the distances between brain areas from the ABA used to approximate the tract lengths of long-range connections. Panel **A**: The distribution of connections in the entire mouse brain is slightly skewed but unimodal with a mean and median of about 4.511 mm and 4.394 mm. The time delays due to the signal transmission shift the local activity by half a period via the longest connections and mostly by about 3π/8 for medium-length connections. The distances directly translate into time delays by assuming an average conduction speed for white matter axonal fibers of 1 m/s. The connections appear of similar length regarding the entire mouse in panel **A**, the connections differ regarding the isocortex in panel **B**. The intrahemispheric connections are shorter than the connections between the isocortical hemispheres. This result confirms the report in Braitenberg and Schüz, 1998, Chapter 26, of a second peak in the histogram whose significance is obscured. The isocortical connections are outnumbered, thus their quantitative effect on the entire mouse, in A, is marginal. Time delays translate into shifts of transmitted local brain activity. The local activity in the network model of the mouse brain is assumed to primarily convey the natural frequency at each brain area (about 42 Hz). The time delays translate into a phase shift in local activity throughout its transmission. The number of bins is 119 with a bin width of 0.0889 mm in panel **A** and 17 bins of 0.3056 mm width in panel **B**. The number of bins in the histogram, *n*_bins_, was calculated according to *n*_bins_ = exp(0.626 + 0.4 log (*n* − 1)), with the number of connections *n* (Otnes and Enochson, 1972), which is listed in Table 4. The statistics are given in Table 5.

#### Local areas and their dynamics

Each of the 512 brain areas is assumed to intrinsically perform an oscillation with a natural frequency in the gamma range (at about 42 Hz). This rhythm in the gamma band accounts for local activity, such as a coordinated interaction of excitation and inhibition (Buzsáki and Wang, 2012), which is not physiologically, but phenomenologically modeled here (Palmigiano et al., 2017). Having said that, the damping factor *γ* determines the local excitability (i.e., the critical distance to the Andronov-Hopf bifurcation), and is thus a surrogate for the ratio of excitation and inhibition at each brain area. A two dimensional (2D) model (i.e., consisting of two state variables: mean postsynaptic potential and current) is used to provide the temporal dynamics of each brain area. In case of the isocortex, where brain areas, that is, isocortical areas are spatially extended, each mesh vertex holds a 2D-element. The 2D-model is equally parameterized for all brain areas by *γ* = 1.21, *ε* = 12.3083. This means that without any links between the brain areas, local behavior is identical. Without any stimulation (no noise, no time-variant input) the activity is constantly zero. The local response to stimulation is a damped oscillation in the gamma frequency range. Because the long-range connections in the structural network of the mouse brain are heterogeneous (see Figure 14), subsequently emerging brain function (and dysfunction) is shaped by the brain structure through the network interactions and, hence, the local response can be highly specific to a brain area though the areas are equally parameterized.

#### State system and its parameterization

The modeling approach to generate brain activity in The Virtual Brain (TVB) involves three interacting systems, namely, concerning the inputs, the dynamics and the observations. Where the *input system* allows for interventions and the *observer system* specifies the measurement modalities, the *dynamical system* generates the brain activity and constitutes brain states. The physical and spatiotemporal properties of actions to brains determine the input to TVB (compare, for example, focal brief direct electrical stimulation, transcranial direct current stimulation and retinal stimulation). In general, models are agnostic about the action and hence have the potential to integrate and moreover to discuss the effects of different modalities of action such as of sensory, direct electrical and transcranial stimulation but also blocked arteries and changes of myelin sheaths. The input system relates an action to the state variables of TVB (e.g., postsynaptic potentials and currents) causing a set of events (over a relative short time scale, for example, event-related potentials (e.g., Shah et al., 2004) and event-related (de)synchronization (Pfurtscheller & Lopes da Silva 2001), and long-term effects such as necrosis and changes in apoptosis and autophagy). Such events may be symptomatic for medical conditions and diseases such as ischemic stroke (as a result of blocked artery) and multiple sclerosis (due to damage of myelin sheath). The dynamical system comprises the variables with which the brain activity is modeled. The model is a network of 14,400 nodes (13,972 vertices for the 84 spatially extended areas on the isocortical surface, and 428 nodes, each for a non-isocortical brain area). Links between brain areas via the homogeneous SC (short-range connections) and the heterogeneous SC (long-range connections) are assumed to be excitatory. Inhibition is implicitly present in the gamma oscillator representing local activity of each brain area, which is distinguishable in the action (excitatory or inhibitory) of their projections. To our best knowledge, however, neither a complete nor an extensive classification is currently available for the characterization of all 512 considered brain areas in the model (e.g., to excite or to inhibit synaptically connected area). The weights of connections are normalized to maximum unity in-strength of a brain area so that the network is segregating and integration information, where the energy of activities decays in the network. In other words, the network cannot amplify activity by connectivity weights greater one. The local dynamics are described by the 2D-model (see, for example, Spiegler et al., 2016). Consequently, the state of the system is spanned by 28,800 state variables. Note that the local behavior was set to a subcritical working point as suggested by many authors within the resting state literature (Deco et al., 2011; Fagerholm et al., 2015). Subcriticality is here represented by a stable focus close to instability, showing damped oscillations in response to perturbations. The specific connections to a given network node (brain area) and the energy that they transfer from connected nodes change the operating point of the local node towards the critical point (i.e., Andronov-Hopf bifurcation) at which the decay time of a stimulation response increase and the node starts performing a self-sustained oscillations (in absence of any stimulation). The brain model is however parameterized that none of the applied stimulation led to a self-sustained oscillation (i.e., limit cycle). Consequently, the brain network segregates and integrates information, where the energy is dissipated by the local intrinsic dynamics. The model parameters are the six configurations for the spatial range of the homogeneous SC on the isocortical surface and the ratio of homogeneous to heterogeneous SC. The brief action of the stimulation is related to the set of nodes spanning a given brain area by the input system. The input system adds the input describing brief focal stimulation of brain areas to the postsynaptic potentials (PSPs).

##### Observer system for voltage-sensitive dye (VSD) imaging

The physical forward solution is captured by means of an observer system links, which links the variables of the state system (here, the mean postsynaptic potential and current) for the 2D-model at each network node, to an experimentally observable measure. To compare the activity in the isocortex due to stimulation in the model and in the experiment during the voltage-sensitive dye (VSD) imaging (Mohajerani et al., 2013) the simulated brain activity is projected onto the plane that reconstructs the focal plane from which the camera setup captures the VSD images. VSD imaging measures the emitted light from the superficial layers of the isocortex stained with VSD molecules as a function of the neuronal membrane potential. The results show that the temporal fluctuations in the light emission are correlated with the electrophysiological measures such as local field potentials (Mohajerani et al., 2013). This suggests that VSD imaging reflects postsynaptic potentials (PSPs) of neural populations. Consequently, we establish a direct link between the state variables of neural activity and the VSD imaging data by equating the first variable of the 2D-model with the PSP of a local population and the second variable, which is the temporal derivative of the PSP, with the current. The PSP of a local population can be linked to the VSD imaging. However, the VSD imaging projects only the brain activity within a limited spatial range around the focal plane (i.e., the depth of field) intersecting the isocortex onto a plane (image sensor) regardless of the curvature of the isocortex in physical space. In the VSD imaging data, isocortical brain areas were identified by the early response to different sensory and peripheral stimulation (i.e., whisker, auditory, forelimb, hindlimb, and visual). These locations were used to find a projection plane (size given by the camera sensor used for the VSD imaging) in the model that best corresponds to the VSD imaging data (see Figure 15). This mapping provides then a direct link of the local PSP in the model to the measured VSD images. The isocortical areas, captured by the VSD imaging setup (see, for more details, Mohajerani et al., 2013), are listed in Table 6. Simulated and measured activity can thus be compared on the imaging level as well as by the region mapping.

**Table 6.** The experimental voltage sensitive dye (VSD) imaging captures activity of isocortical brain areas that are given in the virtual mouse brain model by the ABA. The numbering (#) of the 26 listed areas (out of 84 isocortical areas) in the isocortex (total number of 512 brain areas) captured by the VSD imaging technique corresponds to the labels in Panel **B** of Figure 15.

| # | Areas captured by the VSD imaging | # | Areas captured by the VSD imaging |
| --- | --- | --- | --- |
| 1 | Frontal pole | 14 | Primary auditory area |
| 2 | Anterior cingulate area, dorsal part | 15 | Primary somatosensory area, trunk |
| 3 | Secondary motor area | 16 | Anterior area |
| 4 | Primary motor area | 17 | Rostrolateral visual area |
| 5 | Primary somatosensory area, mouth | 18 | Anterolateral visual area |
| 6 | Supplemental somatosensory area | 19 | Retrosplenial area, dorsal part |
| 7 | Primary somatosensory area, upper limb | 20 | Retrosplenial area, lateral agranular part |
| 8 | Primary somatosensory area, nose | 21 | Anteromedial visual area |
| 9 | Ventral auditory area | 22 | Posteromedial visual area |
| 10 | Primary somatosensory area, lower limb | 23 | Primary visual area |
| 11 | Primary somatosensory area, unassigned | 24 | Lateral visual area |
| 12 | Primary somatosensory area, barrel field | 25 | Laterointermediate area |
| 13 | Dorsal auditory area | 26 | Temporal association areas |

**Figure 15.**
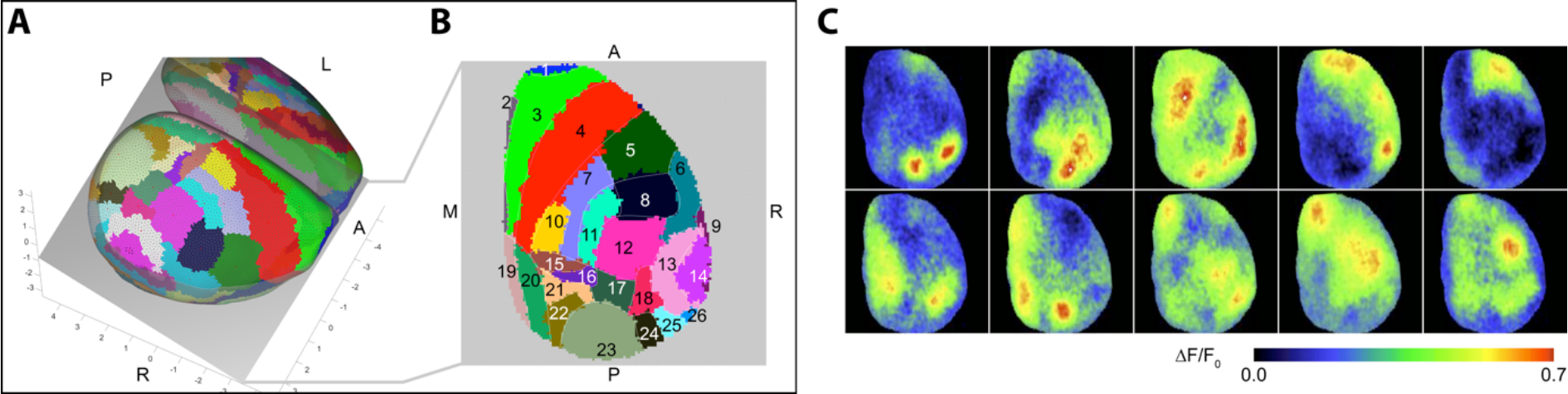
Modeling the focal plane for voltage sensitive dye (VSD) imaging. Panel **A**: The camera setup in the mice experiments was reproduced in the 3 dimensional geometric model of the mouse brain. The focus was first upon the surface of the right isocortex and then adjusted 1 mm inside the cortex so that the depth of field covers most of the right isocortex (for the experimental setup see Mohajerani et al., 2013). The consequence is that the focal plane cuts through the right isocortex (see the gray plane in A) and records brain activity from a wide field of the isocortex. Panel **B**: The visible neural masses are projected onto the focal plane, using the normal vector of the focal plane (i.e., assuming an infinite focal length). The modeled focal plane is sampled by an array of 128 × 128 pixels in accordance with the camera sensor used in the experiments for the VSD imaging recordings. Because, each brain area in the isocortex comprises several neural masses, the areas are colored in both panels **A** and **B**. The areas captured by the VSD imaging are numbered in panel **B** and listed accordingly in Table 6. The labels A, P, L, R, M denote locations: (A)nterior, (P)osterior, (L)eft, (R)ight, and (M)edial. Panel **C**: Experimental VSD imaging of spontaneous brain activity in isoflurane anesthetized mouse show spatial organization (here ten exemplary patterns) demonstrating the qualitative correspondence of experimental and modeled VSD imaging purely based on the ABA parcellation. Note that the (secondary and primary) motor areas (areas 3 in green and 4 in red in the panels **A** and **B**) and such somatosensory areas as the barrel field (area 12 in magenta in the panels **A** and **B**) represent spatially extended areas that are functionally undivided (e.g., by topographic maps) in the ABA. The VSD imaging can indicate activity patterns that are finer grained than the used ABA division of the isocortex into brain areas in the model. The snapshots of the experimental VSD imaging of spontaneous activity in panel **C** are taken and adapted with permission from already published data (Supplementary Video 2) by Mohajerani et al., 2013.

## DISCUSSION

Brain network dynamics can be *systematically* explored only in silico. To explore the mouse brain dynamics in silico, we built a high-resolution whole mouse brain model. The model structure is based on the database Allen Brain Atlas (ABA) comprising tracer studies. We investigated the network behavior by assuming simple oscillatory but canonical dynamics at each brain region, that is, a network node. Each region had the same local dynamics, only differentiated by the ABA connectome from one to the other. We simulated the whole-brain mouse model on a computer using the neuroinformatics platform The Virtual Brain (TVB). To test whether the mouse brain activity can be exclusively inferred from the underlying connection topology, we performed graph theoretic analysis of the model structure. The results show that the overall performance of the 13 graph-theoretic measures is rather poor. The best prediction of the simulated brain responses to focal stimulation and the extracted dynamically responsive networks (DRNs) resulted from comparing the rank of the activity with the rank of connectivity strengths of the direct projection from the stimulation site, where a high correlation indicates that the energy induced by the stimulation spreads into areas of the brain that are directly connected to the stimulation site. The rank correlation is weak but significant for most connectivity parameter configurations. We compared measured and simulated brain activity using the observer model for (virtual) VSD imaging. For exploring the mouse brain dynamics in silico, we systematically varied three model parameters, namely (i) the ratio to which extend brain areas can interact over short-and long-ranges, (ii) the range of short connections that are homogeneous on the isocortex, and (iii) the location of the focal stimulation. The extent to which information is processed over short-or long-range SC is unclear. The results show that both SC types of short-and long-range connections in the mouse model act as spatiotemporal filters of local dynamics in the emergent functional network. The results (Figure 3) do not support evidence for a reorganization of dynamics due to interplay between both types of SC in the mouse as compared to human models (Spiegler et al., 2016). We performed analyses of the brain responses to stimulation. The focal stimulation is closely related to direct electrical, sensory and optical stimulation. We identified consistent spatial motifs of mouse brain activity called dynamically responsive networks (DRNs) using dimension reduction methods. We found the experimentally known functional networks back in the DRNs. Interestingly, the salience and the default mode network spanned several motifs indicating the salience and default mode network to be a junction for functional networks. We furthermore investigated the spatiotemporal organization of brain activity following sensory pathways. The model structure reflects the sensory systems, namely: the auditory pathways, the visual pathways, the whisker pathways and the pathways for the limbs. Finally, with the developed large-scale mouse brain network model we had, for the first time, the chance to systematically investigate sensory pathways and, especially, the processing of sensory data. We showed that the in-silico brain responses resemble in-vivo brain dynamics after stimulation. In the following we discuss the results in more detail.

### Modeling results

The structure of the model (Figure 1) based on the database of the Allen Brain Atlas comprising tracer studies was coregistered with the anatomy of the mouse brain (see Oh et al., 2014). We spatially sampled the brain volume given by the ABA and obtained a network model composed of 512 distinct brain areas. The model incorporates the smooth surface of the isocortex that we reconstructed from the ABA using a regular mesh of 27,554 triangles with 41,524 edges between 13,972 vertices (Figure 11) and the spatial extent of the 84 cortical areas given by the ABA. The 512 distinct brain areas contain the 84 isocortical areas (42 per hemisphere) and 428 non-isocortex areas (see Table 1 for the division, Table 2 and Table 3 for a complete list of area names). These 428 sub-isocortical areas are lumped in the model to a point in physical space (i.e., centroid). From these numbers, it is evident that the isocortex constitutes only a small part of brain areas considered in the model (and in the ABA). However, the isocortex is the structure from which the here considered wide field VSD imaging captures activity from 26 isocortical areas in the ABA. Thus we often observe only from a small fraction of interconnected areas (here 26 observables vs. 512 brain areas in the model). Here it is important that the spatial resolution in the model coincides with the resolution of the model. Although the resolution of the brain geometry (e.g., the isocortical mesh) agrees with the measurements (e.g., VSD imaging), the quality of the modeling highly depends on the division of the brain into areas (parcellation) and with that the resolution and reconstruction of connections (Proix et al., 2016). The version of the ABA that we have used to construct the model, for instance, the barrel field of the primary somatosensory areas is summarized by one single area (see Table 2). A distinction of the barrels in the barrel field of the primary somatosensory areas would improve to distinguish lower and upper limb (especially hind and forelimb), see Figure 6 panel E. The model can be improved by using not only the new ABA model (https://www.biorxiv.org/content/early/2018/04/01/293019) but also by incorporating other databases (e.g., http://www.mouseimaging.ca/research/mouse_atlas.html; Dorr et al., 2008; www.MouseConnectome.org).

### Modeling brain dynamics and limitations

We assumed a natural frequency of 42 Hz at each network node and 1 m/s of conduction speed - both within biologically plausible ranges (see section ‘Conduction speed and transmission delays’ and section ‘Local areas and their dynamics’ in the Results). By this simple dynamics at each node we probed the experimentally derived large-scale brain network to produce functionally relevant organizations in the activity by stimulation. If the model equipped with this simple dynamics does not show functionally relevant organizations in the simulated brain activity then the model misses relevant complexity. The experimentally derived large-scale brain network may miss essential features (perhaps the average brain structure is not sufficient and averages out important components) and better data needs to be provided (better estimates for the connectivity strengths). The behavior in the large-scale brain signals (e.g., M/EEG and functional MRI) associated with brain function and dysfunction are perhaps epiphenomena at the large scale of the brain and actually originate from processes on a micro-and meso-scale (rather than throughout the interaction of brain areas) that are reflected onto the large scale of the whole brain and its functional measurements. Another point is the local dynamics itself. The local model for the simple dynamics that we use here incudes a surrogate parameter for the ratio of excitation and inhibition. However, this model is generic and does not represent a particular local structure. If this model is not sufficient, biologically informed models are available. The local dynamic models can be biologically informed about the local circuitry and specific activities (e.g., certain rhythms). Local circuitries are sketches of characteristic elements (e.g., projection neurons, interneurons) and their connections involved in relevant processes (e.g., excitation/inhibition) based on morphological and physiological arguments. Using such biologically informed models increase the repertoire of dynamics at a node by their local circuitry and are thus meso-and microscopic descriptions. It is worthwhile to mention that the complexity in the behavior is linked to the complexity of the structure. Complex behavior (e.g., occurrence of spike-wave complexes or epileptic discharges in distant brain areas) indicates the complexity of systems and therefore cannot be embedded in simple systems. If brain dynamics are provided large-scale models are able to provide plausible than the large-scale network provides entry points for interventions, useful for diagnostics and therapy among others.

### DRNs and functional networks

We performed a systematic exploration of brain dynamics via focal stimulation to extract the DRNs. The DRNs are constrained by the local model dynamics and the connection topology. The interplay of these factors imposes conditions on the dynamic repertoire of a given network, independent on the details of how these factors are physiologically realized. To be specific, there is a multitude of ways to generate a particular behavior of a neural population (Marder & Taylor 2011) such as an oscillation, but once established, then it does communicate via the connectivity with other regions and the actual physiological underpinnings play no role anymore (unless new factors arise such as spike rate adaptation, for instance). From this perspective, the DRNs fundamentally reflect the functional dynamic repertoire of a network given a particular connectivity. The DRNs and the mapping to stimulation allows for a coordinated interaction with functional networks using different types of stimulation (e.g., sensory and transcranial). The DRNs show characteristic motifs. Each motif is a spatial distinct organization of brain activity and composed by three parts. We identified five different spatial patterns in the parts that entangle the motifs. The presence of a pattern in several motifs suggests that the brain is able to phrase more than one motif (thus more than one DRN) at a time. The experimentally known functional networks, especially the default mode network (and the salience network) are reflected in parts of these motifs.

With the present modeling study, we extracted the brain motifs, the possible expressions, and how it is organized. The spatial patterns tell us which brain area is active. We also describe the themes by relating the patterns to experimentally known functional networks. The used local model is simple and appropriate for investigating the large-scale spatiotemporal organization of brain responses, which is the scope of this work. As a consequence of the mono-rhythmic behavior and the uniform parameterization of the local dynamic model, the simulated network dynamics show limitations in describing the various rhythms involved in the brain organization (i.e., neuronal firing and brain rhythms alpha to gamma). For the purpose of describing the broadband frequency behavior (as observable in EEG), the network model has to allow the generation of more complex activity, which could be potential subject to subsequent studies. The modeling is challenging and goes hand in hand with the debate about the generation of brain activity such as rhythms and especially its spatial extent. The risk in this kind of modeling work based on large-scale data (as with modeling based on microscopy data, see Marder & Taylor, 2008) lies in the overfitting (additional state variable increase the degree of freedom and may increase the complexity of the model). One approach would be to use the same simple local dynamic model for each node as we have done in this work but then to parameterize nodes differently accounting for the spatial differences in natural frequencies.

### Sensory pathways

The sensory networks in Figure 6 are mainly based on textbook descriptions (e.g., Watson et al., 2011) and include the relevant structures given by the ABA. The sensory pathways can be described using the ABA. However, the ABA needs refinements to distinguish lower and upper limb (especially hind and forelimb), see panel E. Neither does the ABA include a division of the barrel field of the primary somatosensory areas. The networks in Figure 6 indicate information flows to areas that do not necessarily terminate in the primary sensory areas of the isocortex, such as the hypothalamic and midbrain targets of the retinal ganglion cells in in Figure 6, panel **A**, and the midbrain nuclei related to whisker movements in Figure 6, panel **D**. These connections are well known and usually not discussed regarding sensory processing in textbooks. As a result, visual system and whisker system meet in the sensory related superior colliculus regarding eye and whisker movements. It is worth mentioning that the model is agnostic about the (sensory) information, meaning that nuclei may be activated but the object of processing (information) is less defined. The vast amount of studies about the physiology of nuclei allows attaching meaning to the modeling results. For instance, the hypothalamic nuclei and the nuclei in the midbrain (nucleus of the optic tract, olivary and posterior pretectal nucleus) receive input from the retina but are not directly part of the visual pathways running from the retina to in the primary visual cortices, see Figure 6, panel **A**. The hypothalamic nuclei play however a role in the circadian timing system and the nuclei in the midbrain are known to be involved in eye movement coordination and reflexes.

Certain patterns in the auditory and visual pathway belong to the same cluster. This is most likely because of the spatial proximity of the inferior colliculus to the superior colliculus (sensory related). The resolution of the mouse tracer studies may be not good enough because of leakage.

There is also overlap between whisker and limbs pathways. The barrel cortex (i.e., Left Primary somatosensory area, barrel field (area 5)) is only separated from the limb areas (i.e., Left Primary somatosensory area, lower limb (area 6) and Left Primary somatosensory area, upper limb (area 8)) by an unassigned area, that is, Left Primary somatosensory area, unassigned (area 10). Consequently, the parcellation might be misleading taking into account that we are dealing with an average mouse brain compiled by numerous different brain of (genetically the same mouse type). This may explain the uncertainty in the matching by errors in the allocations of brain tissues.

Ventral posteromedial nucleus, Ventral posterolateral nucleus, and Posterior complex are neighboring areas and part of the posterior nuclei of the thalamus. The grouping may indicate an uncertainty and limitation of the data because of the proximity and the diffusion of the tracer locally during the measurement technique and the agglomeration of different experiments.

### Translation Human Animal

With this study we adapted the focal stimulation protocol that we have used previously in-silico for exploring human brain dynamics. Although the mouse brain is way smaller than the human brain, the models are comparable in terms of number of nodes used to describe the brain network (in 16,500 human vs. 14,000 in mouse). However, because of the different brain sizes, brain areas, especially the subcortical areas are described in more detail in the mouse brain than in the human (76 cortical areas and 116 thalamic areas in the human model vs. 84 isocortical areas and 428 subisocortical areas in the mouse model). The mouse brain model used in this study is most detailed in the subisocortical structures, namely cerebellum, medulla, pons, midbrain, hypothalamus, thalamus, and cerebral nuclei). It does provide an excellent validation framework for the large-scale brain network modelling approach.

One of the main questions was to which extent information is processed via short-and long-range connections. The result in human was that the short-range connectivity is relevant for explaining experimentally known functional networks. By varying the spatial extent of short-range connections and varying the ratio of short-to long-range connections we observed a reorganization of the DRNs and the brain dynamics. This reorganization of brain dynamics due to changes in the connectivity could not be found in the mouse model. The model suggests that the mouse brain dynamics are organized around 12 motifs (see Figure 5). These motifs are fairly stable in the mix of short-range and long-range connections (see Figure 3).

The motifs of the DRNs in both the mouse and the human model reflect functional networks. Whereas the DRNs in human span a variety of functional networks (e.g., default-mode, memory, attention), the DRNs in mice cover mainly functional networks at rest such as salience, default mode, medial, and lateral networks and less the sensorimotor networks (only the network of the frontal eye field).

Application of stimulation techniques is limited in human to transcranial and sensory stimulation and intracranial stimulation in patients. In animals a battery of interventions is available such as optogenetic tools and electrophysiology. These tools make validation feasible and, moreover, the translation to human through modeling so highly important.

### In-vivo sensory and focal stimulation in the mouse brain

Stimulation paradigms are critical for the validation of any model paradigms, because conceptually they take the system out of its attractor states and allow the sampling of their dynamic neighborhood. The thus obtained information is significantly more predictive than any resting state paradigms.

The simulated data after focal stimulation of the visual and retrospleial cortices match better with the empirical data. We suggest that it is because the tracing experiments provided by the ABA describe the structural connectivity of the visual network more accurately than that of the sensory one. We believe that by incorporating a more accurate structural connectivity of cortical networks, we can achieve a more realistic model, which is able to produce patters of activity resembling the empirical data.

Beyond connectivity, there are other significant simplifications present in the network model affecting its validity, when confronting with empirical data. The most obvious one, from the network perspective, will be the so far assumed homogeneity of local network node dynamics. It is well established that local brain region dynamics varies across the network in terms of excitability (physiological rhythms measured in intracranial human EEG vary across brain regions, see Bartolomei et al., 2017), frequency (posterior to anterior gradient of eigenfrequency ranging from 10 to 40-50 Hz (e.g., Rosanova et al., 2009), and rhythmic organization (burst activity such as in the reticular nucleus of the thalamus (see Marlinski & Beloozerova, 2014; Shermann, 2001); harmonic activity (for example, the alpha waves originating from the occipital lobe, see Niedermeyer, 1997), others behavior such as the activity change in suprachiasmatic nucleus of the hypothalamus course of the circadian rhythm (see Welsh et al., 1995)). These factors are well known to determine the synchronization behavior of networks, although their influence on spatiotemporal energy dissipation and the creation of DRNs may be limited, because of the large-scale nature of network propagation considered here. These are doubtlessly important factors, but connectivity will remain the most critical one, thereby leaving it open the extent to which connectivity details are functionally expressed.

### Conclusion

The full-brain network perspective is very particular. In its essence, it decomposes the brain into a system composed of nodes and links, which is capable of spatiotemporal pattern formation. This view allows us to ask fundamental questions about stimulated pattern propagation, which is at the heart of information processing in the brain. As unsatisfactory the answers are physiologically (due to the nature of the approach), as astonishing is the validity and degree of detail, to which signal propagation following stimulation (sensory or focal) is explained without the need of fine tuning of parameters, further underwriting the fundamental nature of the insights gained here. We have essentially mapped out all the network signal propagation pathways supported by the mouse brain.

## MATERIALS AND METHODS

In order to perform exploration of brain dynamics in silico we model the mouse brain using the large-scale brain model, consisting of the geometry and the structural connectivity combined with local dynamics. We performed simulations of the whole-brain network model, decomposed the brain response after focal brain stimulation, and extracted the dynamically responsive networks (DRNs). By performing statistics, we tested the predictability of the DRNs by the structural connectivity and compared experimentally known functional networks with the DRNs. Furthermore, we investigates the integration of stimulation and response along sensory pathways into the whole brain network. Finally we compared simulated data and voltage sensitive dye imaging and during in-vivo sensory and optogenetic stimulation in the mouse brain. The following sections provide a detailed description of each step:

### Modeling the mouse brain

Using *The Virtual Brain* platform (Melozzi et al., 2017; Sanz-Leon et al., 2013, 2015; http://thevirtualbrain.org/), we triangulate the smooth surface of the isocortex using a regular mesh of 27,554 triangles with 41,524 edges (88 μm is the length of the shortest edge, 190 μm is the longest edge, with a mean edge length of 125 μm) between 13,972 vertices (Figure 1, Panel **A–B**, and Figure 11), distributed across 84 cortical areas given by the Allen Brain Atlas (ABA; http://connectivity.brain-map.org) (Figure 1, Panel **C**). Each area of the isocortex contains between 69 and 667 nodes in the mesh (Table 1 to Table 3) and, because the mesh is regular, the number of nodes indicates the size of an area in the ABA. In addition to the areas in the isocortex, the model includes 428 subisocortical areas. The division of the reconstructed isocortical surface into areas, that is, the parcellation was performed by (i) extending the normal vectors of each vertex to about 200 μm inside the volume of the isocortex given by the ABA, (ii) determining the 5 closest injection sites, and (iii) evaluating their classification to an area of the isocortex in the ABA. The isocortical parcellation was furthermore corrected for holes and isolated/mismatched vertices.

To connect nodes with each other, we distinguish homogeneous from heterogeneous SC (Figure 1, Panels **D**–**F**). The homogenous SC (of short-range connections) links nodes within an area, and between areas if they are spatially close from one another with a connection probability decreasing with distance (Braitenberg and Schüz, 1991, 1998) (Figure 1, Panels **D** and **F**). The heterogeneous SC (of long-range white matter tracts) links all the nodes of an area with the nodes of another area (Figure 1, Panels **D** and **E**), based on known anatomical atlas of the mouse brain, that is, ABA. Neighboring areas are able to exchange information via the homogeneous SC within the cortex and via the white matter tract, that is, heterogeneous SC (e.g. Area 2 with Areas 1 and 3 in Figure 1, Panel **D**).

Each vertex point is a network node holding a neural mass model connected to other nodes via the homogeneous SC and heterogeneous SC. When an area is stimulated, all the nodes of this area are simultaneously activated and then the stimulation-induced activity in each node decays differently according to the activity in the surrounding via short-range connections (i.e., homogeneous SC) and remote nodes via long-range connections (i.e., heterogeneous SC). The ability to drive the network does not depend on the number of nodes within an area, because the heterogeneous SC transfers the mean of the activity in all the nodes within an area to all the nodes in another areas.

We consider this ratio of homogeneous SC to heterogeneous SC as a degree of freedom and perform a parametric study (see Jirsa and Kelso, 2000; Qubbaj and Jirsa, 2007, 2009 for systematic studies with two-point connection). The ratio has been estimated. For instance, Braitenberg and Schüz (1998) assessed that pyramidal cells have synapses in equal shares from long-range and local axons. However, the ratio of homogeneous SC to heterogeneous SC mainly depends on the resolution of the used geometrical model of the cortex, with that the representation of the SC, and the network node description (e.g., canonical model, neural mass model), which is able to incorporate local connectivity (see, for example, Spiegler and Jirsa, 2013 for more detail). At the extremes, (i) 0% of heterogeneous SC (thus 100% of homogeneous SC gives two unconnected cerebral hemispheres with locally but homogeneously connected nodes) only allows activity to propagate locally from a cortical stimulation site, and (ii) 100% of heterogeneous SC (thus 0% of homogenous SC gives 512 purely heterogeneously connected brain areas with locally unconnected nodes) only allows activity to travel long distances with time delays via white matter fiber tracts.

Furthermore, since the spatial range of homogeneous SC is not known (Spiegler and Jirsa, 2013), we also consider it as a parameter varying between 500 μm and 1000 μm. We then systematically stimulate each of the 512 areas with a large range of parameter values (for the ratio and the spatial range), resulting in a total of all 18,432 simulation trials.

Brain dynamics at rest have been found to operate near criticality (Ghosh et al., 2008; Deco et al., 2011, 2013, Spiegler et al., 2016). Near-criticality is defined as a system that is on the brink of a qualitative change in its behavior (Shew and Plenz, 2013). The proximity to criticality predicts that the brain’s response to stimulation will primarily arise from structures and networks that are closest to instability. Activities in those networks require the most time to settle into equilibria after stimulation, and are associated with large-scale dependencies and scale invariance (Haken, 1978). This would be consistent with the center manifold theorem, which states that a high-dimensional system in a subcritical state will converge on a lower dimensional manifold (here few networks) when the system is stimulated. Consequently, we equally set each node in the brain network model to operate close to its critical point, where the network shows no activity without stimulation. We use the stable regimen of each network node (i.e., stable focus) to stimulate a given area in the direction of its instability point (i.e., supercritical Andronov-Hopf bifurcation) and induce characteristic energy dissipation through the brain network. The dissipation of energy will be constrained by the homogeneous SC and heterogeneous SC, the associated signal transmission delays, and the local dynamics at the network nodes. In the network model, the operating point of every node, when disconnected from the network, is at the same distance from its critical point, that is, the supercritical Andronov-Hopf bifurcation (Figure 2, Panel **A**). If the critical point is reached, the node enters into a constant oscillatory mode. In the network, the SC (incl. time delays) determine the alteration of the working distance to the critical point at each node in time by weighting and delaying the incoming activity from other nodes in the network. Hence, network metrics of the SC such as the in-strength, that is, the sum of weights of incoming ties to a node may indicate the distance of a node’s operating point to its critical point and thus criticality (Kunze et al., 2016). The network model, however, is set so that criticality is never reached, by normalizing the SC to unity maximum in-strength so that activity cannot be amplified through the SC. As a result, when a node is stimulated, the node operates closer to the critical point and the response is in the form of a damped oscillation (Figure 2, Panel **A**). The closer a node operates to the critical point, the stronger the node’s responses with high amplitude and long decay time (Figure 2, Panel **A**). The nodes are working near criticality (i.e., they get close to a change in behavior, which would be here a switch to a constant oscillatory mode, but never reaching it). Thus the response to the stimulation is transient, lasting a few milliseconds. The damped oscillation generated in one stimulated node is then sent via its efferent connections to its target nodes, triggering there, in turn, a damped oscillation (Figure 2, Panel **B**). If the network were mainly based on nodes connected in series, activity would decay very fast after the stimulation (Figure 2, Panel **B**). However, since the outgoing activity of a node can influence the nodes projecting back to it, recurrent systems appear (Figure 2, Panel **C–D**), which allow activity to dissipate on a much longer time scale. The evoked activity, after the initial decay, thus persists in the so-called responsive networks (**Fig. 2c, d**), which may reflect feedback loops and re-entry points in the SC. A dynamically responsive network acts on changes, for instance, due to sensory stimuli and random fluctuations in the network (flexibility), and outlasts the stimulation (criticality).

The described network properties are illustrated in Figure 11 to Figure 14 as well as Table 4 and Table 5. The differences in the response stem from the proximity to criticality, which depends upon the SC (in particular the extent of recurrent networks), comprising the synaptic weights and the time delays (Figure 1). This behavior is predicted by the center manifold theorem, which is the mathematical basis for criticality (Haken, 1978).

The directionality of the heterogeneous SC derived from the tracer studies provided by the ABA is assessed by two measures estimating the symmetry in the connectivity weight matrix. The first measure, *Q*_0_ = [0,1], gives 0 if symmetric and 1 if asymmetric and reads:

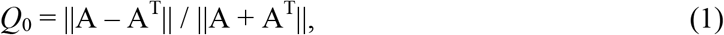

where A is the square matrix containing connection weights between the 512 areas, and ‖A‖ is the matrix norm. The second measure, *Q*_1_ = [0,1] is defined as follows:

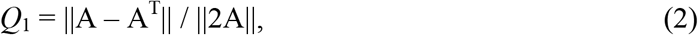

and gives *Q*_1_ = 0 if the matrix A is symmetric and *Q*_1_ = 1 if anti-symmetric. Both measures indicate a high symmetry in the weight matrix of the ABA SC and therefore a dominant bi-directionality of connectivity.

### Large-scale brain model

Dynamics of a vector field Ψ (*x*, *t*) at time *t* ∈ ℝ^1^ and position *x* ∈ ℝ^3^ in space Ω are described by a delay-integro-differential equation:

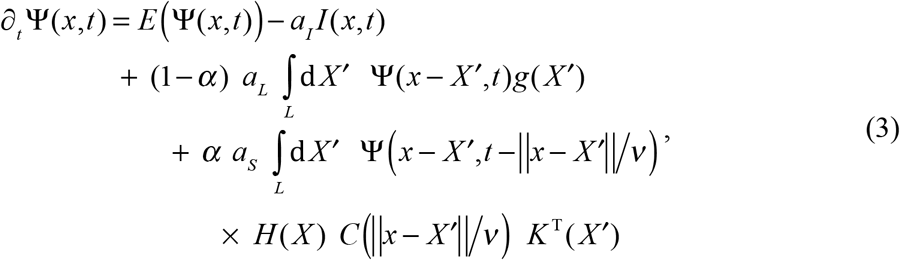

were ∂*_t_* is the derivative with respect to time, *t*. The input *I* (*x*, *t*) allows the stimulation dynamics to intervene on a node. The operator *E* (Ψ (*x*, *t*)) locally links variables of the vector field. The scalar *α* balances the effect of the homogeneous SC and the heterogeneous SC (first and second integral) on the vector field. The vectors *a*_*I*_, *a*_*L*_, and *a*_*S*_ of factors relate the input *I*, and both types of SC to the vector field Ψ (*x*, *t*). The kernel *g* (*x*) describes the homogeneous SC. The field is time delayed due to a finite transmission speed *v* via the heterogeneous SC given by matrix *C* (*x*). The vectors *H* (*x*) and *K* (*x*) establish the links between the heterogeneous SC and the targets and the sources. Note that the transmission speed enters the second integral concerning heterogeneous SC. We assumed the transmission via the homogeneous SC (first integral) to be instantaneous, which reduces the computational expenses, in order to perform the parameter study. The spatial and temporal aspects of the model are described in more detail in the following two subsections.

### Geometry and structural connectivity (SC)

The spatial domain Ω = {*L*_1_ ∪ *L*_2_ ∪ *S*} separates both cerebral hemispheres *L* = {*L*_1_ ∪ *L*_2_ }: left, *L*_1_ and right, *L*_2_, from subisocortical areas *S*, that is, ∩Ω = Ø. Two open surfaces describe the geometry of the left and right isocortex (*L*_1_ and *L*_2_) by a regular mesh of 27,554 triangles with 41,524 edges between 13,972 vertices. The homogeneous SC follows a Gaussian distribution *g* (*x*) = exp (−*x*^2^ / (2*σ*^2^)), which is invariant under translations on *L* (Spiegler and Jirsa, 2013). Each open surface, *L*_1_ and *L*_2_, is divided into *m* = 42 areas, that is, *L*_1_ = ∪_*r* ∈ *R*1_ *A_r_* and *L*_2_ = ∪_*r* ∈ *R*2_ *A_r_* with *R*_1_ = *R*(*m*), *R*_2_ = *R*_1_ + *n*: *R*(*λ* ∈ ℕ) = {*r* | *r* ℕ, *r* ≤ *λ*}, where *n* = 428 is the number of subcortical areas. The division of the surfaces into areas follows the classification of volumes with regards to the tracer studies provided by the ABA, *A_r_* = *A*(r ∈ ℕ) ∈ Ω : ℕ → ℝ^3^ onto space Ω for introducing heterogeneous SC. Each of the *n* = 428 considered subisocortical areas is lumped to a single point in space *S* = ∪_*r* ∈ *R*3_ *A_r_* with *R*_3_ = *R*(*n*) + 2*m*. The heterogeneous connections, *C* transmit mean activities of sources to target areas, *H* (*x*) and *K* (*X′*) with a finite transmission speed, *v* = 1 ms^−1^. The square matrix, *C* (‖ *x* − *X′* ‖ / *v*) contains (2*m* + *n*)^2^ weights, *c_ij_* (‖ *x* − *X′* ‖ / *v*) : *i*, *j* = 1, …, 2*m* + *n* taken from the ABA. The row vectors *H* (*x*) and *K* (*X′*) contain 2*m* + *n* operations, *h_i_* (*x*) and *k_j_* (*X′*) on the targets and sources, respectively. The operations are *h_i_* (*x*) = *δ_x_* (*A_i_*) and *k_j_* (*X′*) = *δ_X′_* (*A_i_*) / |*A_j_*| with the Dirac measure *δ*_Ω_ (*A*) on Ω and the cardinality |*A_r_*| of the set *A_r_*.

The description of the large-scale brain network model (Equation 1) is fully compatible with previous TVB descriptions (Sanz-Leon et al., 2015; Spiegler et al., 2013, 2016). Note that the set notation is used here to describe brain areas and the division of homogeneously distributed and connected network nodes on both isocortices into cerebral areas.

### Local dynamics

The vector field describes a two-dimensional flow (Stefanescu and Jirsa, 2008) linking two variables Ψ (*x*, *t*) = (*Ψ*_1_ *Ψ*_2_)^T^ (*x*, *t*) in (1) as follows

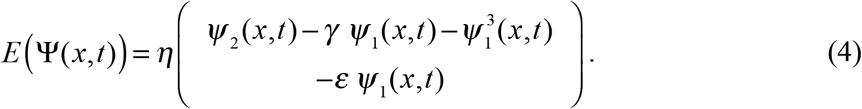

The parameterization: *γ* = 1.21 and *ε* = 12.3083 sets an isolated brain area close to a critical point, that is, an Andronov-Hopf bifurcation (sketched in Figure 2) with a natural frequency around 42 Hz using a characteristic rate of *η* = 76.74 s^−1^. This rhythm in the gamma band accounts for local activity such as a coordinated interaction of excitation and inhibition (Buzsáki and Wang, 2012) that is not explicitly modeled here but represented by the surrogate parameter *γ*. The Dirac delta function is applied to a brain area, *I*_*r*_ (*x*, *t*) = −5*η δ_x_* (*A*_*r*_) *δ* (*t*). The connectivities and the input act on the first variable *Ψ*_1_ (*x*, *t*) in (1) by *a_L_* = *a_S_* = (*a_I_*)^T^ = (*η* 0). The connectivity-weighted input determines criticality by working against inherent energy dissipation (i.e., stable focus) towards the bifurcation. So that the bifurcation was not passed, both homogeneous and heterogeneous SC, *g* (*x*) and *C* (‖ x − *X′* ‖ / *v*) are normalized to unity maximum in-strength across time delays by: (i) ∫ d*x g* (*x*) = 1, and (ii) 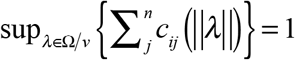

### Simulation

To simulate the model on a digital computer, physical space and time are discretized. The folding of the human cortex presents a challenge for sampling. The surface of the left isocortex *L*_1_ is evenly filled with 7,020 nodes and the surface for the right isocortex *L*_2_ is evenly filled with 6,952 nodes. Subcortical structures in *S* remain unaffected by the discretization. The geometry of the brain is captured in physical space, Ω by a net of 14,000 nodes (i.e. 13,972 cortical and 426 subisocortical nodes). The spatial integrals in (1) are rewritten as matrix operations, where the heterogeneous SC remains the same and the homogeneous SC is spatially sampled on the cerebral surfaces (Spiegler and Jirsa, 2013). The system of difference equations are then solved using Heun’s method with a time step of 40 μs for 1 second per realization of one of the following factors: each of the 512 stimulation sites, SC-balance, α = {0.0, 0.2, 0.4, 0.6, 0.8, 1.0}, and homogeneous spreading, σ / μm ∈ ℕ: 500 ≤ σ / μm ≤ 1000. The implementation is verified by the algebraic solution of an isolated node (i.e., no connections), and by the field properties (e.g., compact solutions spreading radially around a stimulation site) of the homogeneously linked cerebral nodes.

The lower bound of the spatial range of σ = 500 μm results from the used geometrical model for the isocortex. A nearly regular mesh of triangles approximates each cerebral hemisphere with a finite edge length of 125 μm on average (see Figure 11). The used Gaussian kernel for the homogeneous SC is sampled in the model through the cortical mesh. Because of the finite edge lengths in the mesh, the spatial range of the homogeneous SC should not fall below 207.78 μm for −3 dB cutoff of spatial frequencies with respect to their magnitude (Spiegler and Jirsa, 2013). The lower bound of the spatial range of σ = 500 μm for the homogeneous Gaussian connectivity kernel causes a loss of at least 4.55% of spatial information (mainly short-range), which corresponds to −17.37178 dB cutoff (see Spiegler and Jirsa, 2013).

### Stimulation and Decomposition

All network nodes of a brain area are constantly stimulated for a period of the characteristic time of the nodes, *η*^−1^ to evoke damped oscillations with a maximum magnitude of one. The stimulation response of an isolated node is subtracted from the response of stimulated nodes in the network. A *Principal Component Analysis* (PCA) was performed using the covariance matrix among the 16,500 nodes. The period of 0.5 s data after 0.2 s of stimulus onset was decomposed (similar to Spiegler et al., 2016). For further analysis, up to three principal components (i.e., orthogonal) are considered that cover more than 99% of variance across conditions.

### Extracting dynamically responsive networks (DRNs)

The dot product of the normalized eigenvectors from the decomposition the stimulation response was used to measure the similarity of the dissipation across different stimulation sites for a range of values of the balance of the SC and a spatial range of the homogeneous SC. The eigenspaces are clustered based on the similarity measure using k-means for each SC-balance and each range of the homogeneous SC. The number of clusters is estimated via the gap statistic (Tibshirani et al., 2001). For each cluster, the eigenspaces are rotated to the basis of the one with the highest similarity among all in the cluster, using the singular value decomposition and calculating the optimal rotation matrix (Kabsch, 1978). Averaging the aligned basis vectors in a cluster (across eigenspaces) gives the set of eigenvectors for each cluster. Each resulting eigenvector indicates the contribution of each network node (e.g., belonging to a cortical or a subcortical structure) to a DRN.

### Statistics on the DRNs and Prediction of the DRNs by the structural connectivity

To performed statistics on the catalogue of DRNs we pair-wise correlated each of the three parts of each of the 24 motifs, that is, twelve motifs by stimulation of left brain areas and twelve motifs by stimulation of right areas (makes a total of 72 parts composing the 24 motifs). The significance level of 1% was Bonferroni corrected by the total number of components. In addition, three thresholds are applied to the correlation values *C* in order to find strong (*C* > 0.8), good (0.6 < *C* ≤ 0.8), and moderate agreements (0.5 < *C* ≤ 0.6) among the 72 parts of the 24 motifs. The analysis reveals a total of five significantly distinct spatial patterns in the motifs of the DRNs, indicating their entanglements.

To test to which extent the topology in the structural connectivity predicts the brain activity due to stimulation, we calculated the following graph-theoretic measures on the long-range SC: in-, out-, total-degree; in-, out-, total-strength; and clustering coefficient (Rubinov and Sporns, 2010). Incoming, outgoing, or all connected ties to an area are measured in terms of (i) their numbers, and (ii) their weights. By counting the connections we obtain the in-, the out-, and the total-degree. By calculating the sum of connection weights we obtain the in-, the out-, and the total-strength. The clustering coefficient measures the degree to which areas in a graph tend to group together. We correlated the activity at the network nodes after stimulation with the graph theoretical characterization of the network nodes. Furthermore, we correlated the activity at the network nodes after stimulation with the connectivity strengths of the direct afferent (incoming connection to an area) and efferent connections (outgoing connection from an area). Each nine measures of the brain areas in the heterogeneous SC are then compared with the parts of each DRN (i.e., the eigenvector). For the correlation we used the Pearson correlation, the rank correlation (Kendall’s tau) and the Bhattacharyya coefficient. To test statistical significance, the same permutation test is used as for the comparison of the DRNs with the functional networks. We Bonferroni corrected the significance level of 5%.

### Sensory pathways

If not explicitly cited, the main reference for the description of the sensory pathways of the mouse brain is the book authored by Watson, Paxinos, and Puelles (2011). The visual pathways run from the retina through the hypothalamus, midbrain, and thalamus to terminate in the primary visual cortices. The retinal ganglion cells bilaterally connect to the hypothalamus and all of them project to the (sensory related) superior colliculus according to the ipsilateral visual hemifield. The (sensory related) superior colliculus projects to the to areas of the cerebral cortex that are involved in controlling eye movements through the pulvinar and lateral posterior nucleus of the thalamus (Stein and Meredith, 1991). Pulvinar nuclei are virtually nonexistent in the rat, and grouped as the lateral posterior-pulvinar complex with the lateral posterior thalamic nucleus due to its small size in cats. In humans it makes up roughly 40% of the thalamus making it the largest of its nuclei (LaBerge, 1999). Furthermore, the (sensory related) superior colliculus also receives activity from other sensory systems such as whisker movements (Stein and Meredith, 1991). The retinal ganglion cells have ramifications to contralateral structures in the midbrain, namely nucleus of the optic tract, olivary pretectal and posterior pretectal nucleus (Pak et al., 1987) and ramifications that run into the primary visual cortex through the thalamus, namely intergeniculate leaflet as well as ventral and dorsal parts of the lateral geniculate complex according to the ipsilateral visual hemifield. Intergeniculate leaflet of the lateral geniculate complex seems to play an important role in mediating phases of the circadian rhythms that are not involved with light, as well as phase shifts that are light-dependent (Harrington, 1997; Edelstein and Amir, 1999). The network with the specific visual hemifield connections is summarized in Figure 6, Panel **A**. The auditory pathways originate from the cochleae to the primary auditory cortices through the dorsal and ventral cochlear nucleus of the medulla, nucleus of the lateral lemniscus of the pons, inferior colliculus, and nucleus of the brachium of the inferior colliculus of the midbrain, and the medial geniculate complex of the thalamus. The specific contra- and ipsilateral connections are summarized in Figure 6, Panel **B**. The pathways from the facial whiskers to the barrel fields run trough the medulla via the trigeminal nerve, the pons, the midbrain, and the thalamus to terminate in the somatosensory areas of the isocortex. In the different brain structures, each individual whisker is represented in a discrete anatomical unit, that is, a barrel (see Figure 6, Panel **C**). The major facial whiskers are topographically organized in the parts of the medulla, the pons, the thalamus, and the isocortex. The somatotopy of the whisker follicles mainly comes to the primary and secondary somatosensory cortices (barrel cortex) via the trigeminal nerve nuclei through the principal sensory nucleus of the trigeminal of the pons and caudal part of the spinal nucleus of the trigeminal of the medulla (somatotopic arrangement of barrelets), and the ventral posteromedial nucleus of the thalamus (somatotopic arrangement of barreloids). Specifically, the trigeminal nerve fibers have ramifications to caudal, interpolar, and oral part of the spinal nucleus of the trigeminal (medulla), to the principal sensory nucleus of the trigeminal of the pons, and to the trigeminal nucleus of the midbrain (Bosman et al., 2011). Whisker movements can activate the sensory related parts of the superior colliculus of the midbrain through the trigeminal nucleus (Matesz, 1981; Ndiaye et al., 2000). The superior colliculus has other parts related to the sensory processing such as the visual system and eye movements. The main sensory pathway is conveys somatotopic information from the whiskers to the barrel field of the primary somatosensory areas of the isocortex via the principal sensory nucleus of the trigeminal (pons), and the ventral posteromedial nucleus (thalamus). The other minor pathways are as follows: The principal sensory nucleus of the trigeminal (part of the pons) mainly projects to the dorsomedial part (and less to the head area) of the ventral posteromedial nucleus of the thalamus, and less to the posterior complex of the thalamus. Posterior complex of the thalamus receives connections from the ventral posteromedial nucleus, the interpolar and oral parts of the spinal nucleus of the trigeminal and sends input to the primary and secondary somatosensory cortices as well as the primary motor cortices (Bosman et al., 2011; Lane et al., 2008). The caudal part of the spinal nucleus of the trigeminal sends input to the ventral posteromedial nucleus and to the lateral dorsal nucleus, which send input to the somatosensory areas. The specific contra-and ipsilateral connections are summarized in Figure 6, Panel **D**. The pathways from the fore and hind limbs pass via dorsal root ganglion to the ipsilateral dorsal column nuclei of the medulla, comprising the cuneate nucleus and the gracile nucleus. The upper limb, and especially the forelimb, connects through the cuneate nucleus, whereas the lower limb, and especially the hind limb, connects through the gracile nucleus. The nuclei of the medulla connect via the medial lemniscus to the contralateral ventral posterolateral nucleus of the thalamus, which has ipsilateral ramifications into the primary somatosensory cortex (S1). The specific contra-and ipsilateral connections are summarized in Figure 6, Panel **E**.

We have quantified the quality of the sensory pathways represented in the ABA connectome as follows: A particular set of anatomical connections (structural connectivity), which are involved in the processing of a particular brain function, is embedded in a large-scale brain network forming sub-networks. Large-scale brain network of directed connections, which weights are assumed to be positively-defined, that is, between network nodes *i* and *j*: *c_ij_* ≠ *c_ji_*!, and *c_ij_* ≥ 0 (the first index is the target and the latter is the source). A particular network comprises a set **R** of edges aka connections **R** ⊆ **r***_n_* between index vertices *u* ∈ ℕ and *ν* ∈ *ℕ* aka nodes in physical space Ω, **r***_n_* = {*ν* → *u*}. The arrow in the notation for the *n* ∈ ℕ-connection **r***_n_* indicates a directed projection from vertex *ν* onto vertex *u*. The embedding of a given connection can be characterized by its particular in-and out-strength in a given large-scale brain network or alternatively the maximum in-and output value:

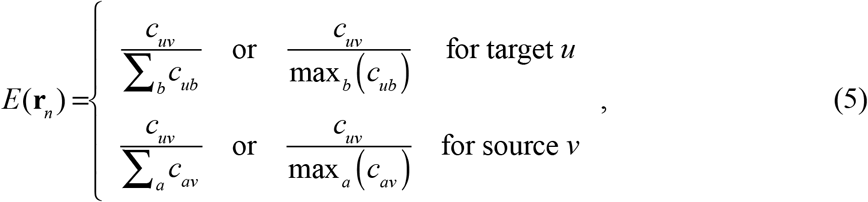

where the maximum income of a target *u* is max*_b_* (*c_ub_*) and the maximum output of a source *v* is max*_a_* (*c_av_*). Using both graph-theoretic measures color the edges **r***_n_* with a positive scalar *c_n_*

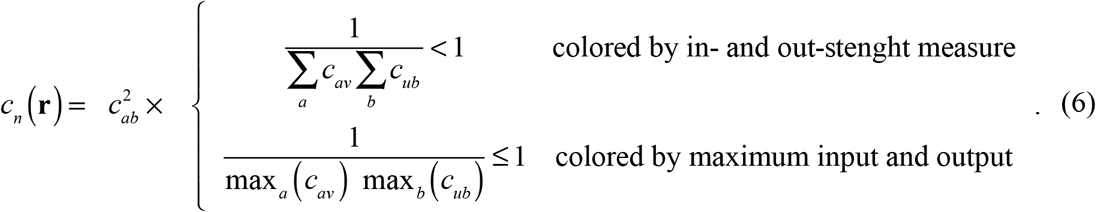

The presence of a subnetwork of interest that is defined by *N* connections **R** ∫ **r***_n_*: *n* = 1, …, *N* in a large-scale network **U** : **R** ∫ **U** can be quantified as follows:

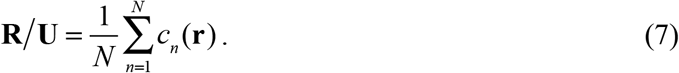

We have used both normalization types to quantify the embedding of sensory pathways. The Pearson correlation and the Bhattacharyya coefficient (Bhattacharyya, 1946) is then used to estimate the amount of overlap (i.e., the square root of the inner product) between a functional network and a DRN, which elements are essentially indicated by an eigenvector (part of a motif). The square of each eigenvector element is taken and summed up within each area. The coarse-grained eigenvectors are normalized to unit length. A functional networks and each normalized coarse-grained part of a DRNs are compared. The *p*-values are corrected due to 27 independent multiple comparisons (nine functional networks with three eigenvectors per stimulation site) with a significance level of 5%.

### Comparing voltage sensitive dye imaging and simulated data

We preprocessed the VSD imaging data by calculating the ratio of the deviation of the optical signal from the baseline and baseline (Δ*F*/*F*_0_) for each pixel separately. Then we converted this ratio to percentage by multiplying it by 100. We defined the baseline as the pre-stimulus level of the optical signal for each pixel. Therefore, a signal level of 0.2% in a particular pixel means that the optical signal captured from that pixel is 0.2% of the baseline for that pixel. Then, we used the Δ*F*/*F*_0_ signal to identify the activated pixels at each time point after the stimulation was applied. We defined the onset of activation for a pixel as the timestamp at which its activity surpasses the 20% of its peak activity. By calculating the onset of the activation for all the cortical regions in our imaging window, we obtained the temporal order of activation of these regions. Moreover, we calculated the maximum of the Δ*F*/*F*_0_ signal for each pixel around a temporal neighborhood of the stimulation time point and obtained the maximum intensity maps in the panels **C** in Figure 8 to Figure 10. These maps ease the qualitative investigation of the overlap of activated regions in the simulated and empirical data.

### In-vivo VSDI and optogenetic stimulation in the mouse brain

To perform the VSD imaging experiments, animals were anesthetized with 0.5% isoflurane, and a large portion of the dorsal skull on the right hemisphere as well as the dura matter were removed. Then, the exposed brain was incubated for 60 – 90 minutes with the dye RH1692 solution prepared with the HEPES-buffered saline (1 mg/ml). After the unbounded dye solution was washed out from the brain, the stained brain surface was covered with the 1.5% agarose dissolved in HEPES-buffered saline and sealed with a glass coverslip. The VSD data was collected as 12-bit images using a CCD camera (1M60 Pantera, Dalsa, Waterloo, ON) and EPIX E4DB frame grabber with XCAP 3.1 imaging software (EPIX, Inc., Buffalo Grove IL). The focal plane was set to about 1 mm bellow the cortical surface in order to reduce the potential brain surface pulsation artifacts. VSD was excited with the red light emitted from a LED (Luxeon K2, 627 nm center) and filtered in the range of (630 ± 15) nm. VSD fluorescence was filtered using a 673–703 nm optical bandpass filter (Semrock, New York, NY) before being captured by the CCD camera.

To identify the coordinates of the sensory areas, periphery stimulation combined with VSD imaging was performed. The peripheral somatosensory, visual, barrel, and auditory stimulation were performed by delivering 1 mA electrical current to the limbs, delivering a short-duration pulse of a green light to the left eye, shaking the whiskers using a pizo-coupled arm, and playing a sound clique next to the left ear, respectively.

The output of a diode pumped solid-state laser emitting 473 nm light (CNI Optoelectronics, Changchun, China), was used to stimulate ChR2-expressing neurons. The laser beam was directed to pre-defined coordinates on the cortical surface using custom software written in IGOR PRO (Portland, OR), which controlled galvanometer scan mirrors (Cambridge Tech, Lexington, MA), via analog output voltage from PCI-6110 DAQ (National Instruments, Austin, TX). Low amplitude and short duration single laser pulses were applied to the cortical surface to ensure sufficient activation, low laser stimulus artifact, and low photo-bleaching.

## SUPPLEMENTARY MATERIALS

**Supplementary Figure 1.**
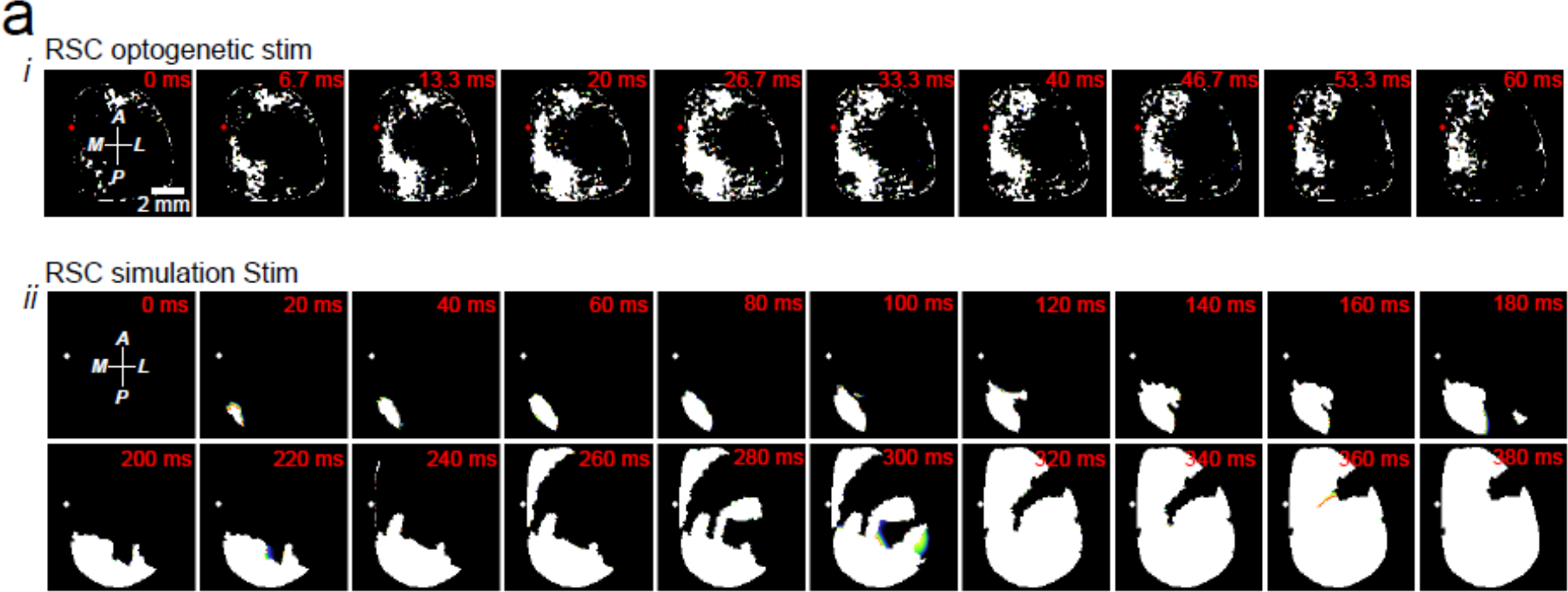
The spatiotemporal pattern of activation of cortical regions after optogenetic (i) and simulated (ii) stimulation of RSC. A region is defined as activated when its activity surpasses 20 percent of its peak activity after stimulation occurred. The white color represents the activated pixels at each frame (timestamp). optogenetic (i) and simulated (ii) stimulation of V2L. A region is defined as activated when its activity surpasses 20 percent of its peak activity after stimulation occurred. The white color represents the activated pixels at each frame (timestamp). its activity surpasses 20 percent of its peak activity after stimulation occurred. The white color represents the activated pixels at each frame (timestamp).

**Supplementary Figure 2.**
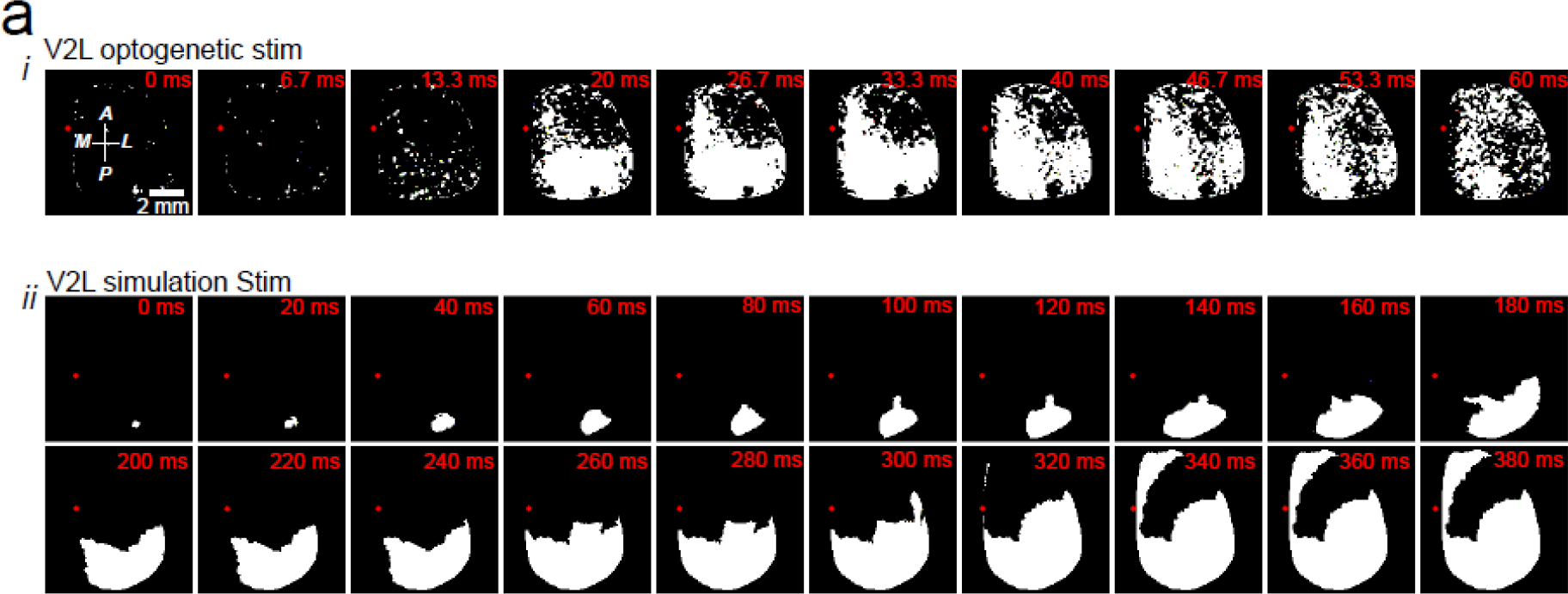
The spatiotemporal pattern of activation of cortical regions after optogenetic (i) and simulated (ii) stimulation of V2L. A region is defined as activated when its activity surpasses 20 percent of its peak activity after stimulation occurred. The white color represents the activated pixels at each frame (timestamp).

**Supplementary Figure 3.**
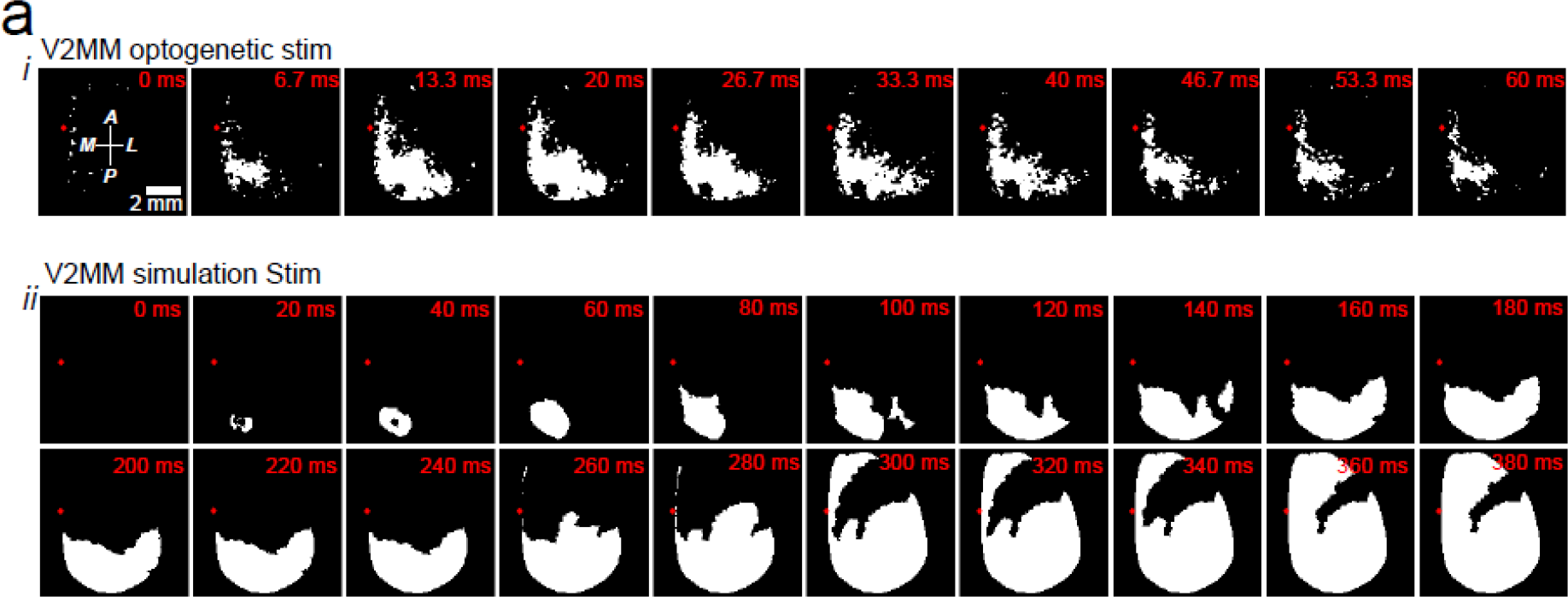
The spatiotemporal pattern of activation 1877 of cortical regions after optogenetic (i) and simulated (ii) stimulation of V2M. A region is defined as activated when its activity surpasses 20 percent of its peak activity after stimulation occurred. The white color represents the activated pixels at each frame (timestamp).

## REFERENCES

http://connectivity.brain-map.org

http://www.mouseimaging.ca/research/mouse_atlas.html

www.MouseConnectome.org

http://thevirtualbrain.org/

Bartolomei F, Lagarde S, Wendling F, McGonigal A, Jirsa V, Guye M, Bénar C (2017) Defining epileptogenic networks: Contribution of SEEG and signal analysis. Epilepsia 58:1131–47, https://doi.org/10.1111/epi.13791

Braitenberg V, Schüz A (1991) Anatomy of the cortex: statistics and geometry. Berlin/Heidelberg: Springer. http://dx.doi.org/10.1007/978-3-s662-02728-8

Bhattacharyya A (1946) On a measure of divergence between two multinomial populations. Sankhya 7:401–6. http://www.jstor.org/stable/25047882

Bosman LW, Houweling AR, Owens CB, Tanke N, Shevchouk OT, Rahmati N, Teunissen WH, Ju C, Gong W, Koekkoek SK, De Zeeuw CI (2011) Anatomical pathways involved in generating and sensing rhythmic whisker movements. Front Integr Neurosci 5:53, https://doi.org/10.3389/fnint.2011.00053

Braitenberg V, Schüz A (1998) Cortex: statistics and geometry of neuronal connectivity. Berlin/Heidelberg: Springer. http://dx.doi.org/10.1007/978-3-662-03733-1

Breakspear M (2017) Dynamic models of large-scale brain activity. Nat Neurosci 20:340–52, https://doi.org/10.1038/nn.4497

Buzsáki G, Anastassiou CA, Koch C (2012) The origin of extracellular fields and currents - EEG, ECoG, LFP and spikes. Nat Rev Neurosci 13:407–20, https://doi.org/10.1038/nrn3241

Buzsáki G, Wang XJ (2012) Mechanisms of gamma oscillations. Annu Rev Neurosci 35:203–25. http://dx.doi.org/10.1146/annurev-neuro-062111-150444

Calvert GA, Spence C, Stein BE (2004) The Handbook of Multisensory Processes. Cambridge, MA: MIT Press, ISBN 9780262033213

David O, Kiebel SJ, Harrison LM, Mattout J, Kilner JM, Friston KJ (2006) Dynamic causal modeling of evoked responses in EEG and MEG. Neuroimage 30:1255–72, https://doi.org/10.1016/j.neuroimage.2005.10.045

Deco G, Jirsa VK, McIntosh AR (2011) Emerging concepts for the dynamical organization of resting-state activity in the brain. Nat Rev Neurosci 12:43–56, http://dx.doi.org/10.1038/nrn2961

Deco G, Jirsa VK, McIntosh AR (2013) Resting brains never rest: computational insights into potential cognitive architectures. Trends Neurosci 36:268–74, https://doi.org/10.1016/j.tins.2013.03.001

Dehaene S, Meyniel F, Wacongne C, Wang L, Pallier C (2015) The Neural Representation of Sequences: From Transition Probabilities to Algebraic Patterns and Linguistic Trees. Neuron 88:2–19, https://doi.org/10.1016/j.neuron.2015.09.019

Denk W, Delaney KR, Gelperin A, Kleinfeld D, Strowbridge BW, Tank DW, Yuste R (1994) Anatomical and functional imaging of neurons using 2-photon laser scanning microscopy. J Neurosci Methods 54:151–62, https://doi.org/10.1016/0165-0270(94)90189-9

Dorr AE, Lerch JP, Spring S, Kabani N, Henkelman RM (2008) High resolution three-dimensional brain atlas using an average magnetic resonance image of 40 adult C57Bl/6J mice. Neuroimage 42(1):60–9, https://doi.org/10.1016/j.neuroimage.2008.03.037

Fagerholm ED, Lorenz R, Scott G, Dinov M, Hellyer PJ, Mirzaei N, Leeson C, Carmichael DW, Sharp DJ, Shew WL, Leech R (2015) Cascades and Cognitive State: Focused Attention Incurs Subcritical Dynamics. J Neurosci 35:4626–34, https://doi.org/10.1523/JNEUROSCI.3694-14.2015

Edelstein K, Amir S (1999) The role of the intergeniculate leaflet in entrainment of circadian rhythms to a skeleton photoperiod. J Neurosci 19:372–80, http://www.ncbi.nlm.nih.gov/pubmed/9870966

Foran DR, Peterson AC (1992) Myelin acquisition in the central nervous system of the mouse revealed by an MBP-Lac Z transgene. J Neurosci 12:4890–7. http://www.jneurosci.org/cgi/pmidlookup?view=long&pmid=1281497

Friston KJ, Harrison L, Penny W (2003) Dynamic causal modelling. Neuroimage 19:1273–302, https://doi.org/10.1016/S1053-8119(03)00202-7

Ghosh A, Rho Y, McIntosh AR, Kötter R, Jirsa VK (2008) Noise during rest enables the exploration of the brain’s dynamic repertoire. PLoS Comput Biol 4:e1000196, http://dx.doi.org/10.1371/journal.pcbi.1000196

Haken H (1978) Synergetics: an introduction nonequilibrium phase transitions and self-organization in physics, chemistry and biology. Berlin/Heidelberg: Springer, http://dx.doi.org/10.1007/978-3-642-96469-5

Harrington ME (1997) The ventral lateral geniculate nucleus and the intergeniculate leaflet: interrelated structures in the visual and circadian systems. Neurosci Biobehav Rev 21:705–27, https://doi.org/10.1016/S0149-7634(96)00019-X

Huys R, Perdikis D, Jirsa VK (2014) Functional architectures and structured flows on manifolds: a dynamical framework for motor behavior. Psychol Rev 121:302–36, https://doi.org/10.1037/a0037014

Krakauer JW, Ghazanfar AA, Gomez-Marin A, MacIver MA, Poeppel D (2017) Neuroscience Needs Behavior: Correcting a Reductionist Bias. Neuron 93:480–90, https://doi.org/10.1016/j.neuron.2016.12.041

Jirsa and Kelso, 2000 Jirsa VK, Kelso JAS (2000) Spatiotemporal pattern formation in neural systems with heterogeneous connection topologies. Phys Rev E Stat Phys Plasmas Fluids Relat Interdiscip Topics 62:8462–5, http://dx.doi.org/10.1103/PhysRevE.62.8462

Jirsa VK, Stacey WC, Quilichini PP, Ivanov AI, Bernard C (2014) On the nature of seizure dynamics. Brain 137:2210–30, https://doi.org/10.1093/brain/awu133

Kabsch W (1978) A discussion of the solution for the best rotation to relate two sets of vectors. Acta Crystallogr A 34:827–8, http://dx.doi.org/10.1107/S0567739478001680

Kunze T, Hunold A, Haueisen J, Jirsa V, Spiegler A (2016) Transcranial direct current stimulation changes resting state functional connectivity: a large-scale brain network modeling study. Neuroimage 140:174–87, http://dx.doi.org/10.1016/j.neuroimage.2016.02.015

LaBerge, D. (1999). Attention pp. 44-98. In Cognitive science (Handbook of Perception and Cognition, Second Edition), Bly BM, Rumelhart DE. (edits). Academic Press ISBN 978-0-12-601730-4 p. 73

Lane RD, Pluto CP, Kenmuir CL, Chiaia NL, Mooney RD (2008) Does reorganization in the cuneate nucleus following neonatal forelimb amputation influence development of anomalous circuits within the somatosensory cortex? J Neurophysiol 99:866–75, https://doi.org/10.1152/jn.00867.2007

Marder E, Taylor AL (2011) Multiple models to capture the variability in biological neurons and networks. Nat Neurosci 14:133–8, https://doi.org/10.1038/nn.2735

Marlinski V, Beloozerova IN (2014) Burst firing of neurons in the thalamic reticular nucleus during locomotion. J Neurophysiol 112:181–92, https://doi.org/10.1152/jn.00366.2013

Matesz C (1981) Peripheral and central distribution of fibres of the mesencephalic trigeminal root in the rat. Neurosci Lett 27:13–7, https://doi.org/10.1016/0304-3940(81)90198-1

Melozzi F, Woodman MM, Jirsa VK, Bernard C (2017) The Virtual Mouse Brain: A Computational Neuroinformatics Platform to Study Whole Mouse Brain Dynamics. eNeuro:4, https://doi.org/10.1523/ENEURO.0111-17.2017

Michel CM, Koenig T (2018) EEG microstates as a tool for studying the temporal dynamics of whole-brain neuronal networks: A review. Neuroimage 180:577–593, https://doi.org/10.1016/j.neuroimage.2017.11.062

Mohajerani MH, Chan AW, Mohsenvand M, LeDue J, Liu R, McVea DA, Boyd JD, Wang YT, Reimers M, Murphy TH (2013) Spontaneous cortical activity alternates between motifs defined by regional axonal projections. Nat Neurosci 16:1426–35. https://doi.org/10.1038/nn.3499

Ndiaye A, Pinganaud G, VanderWerf F, Buisseret-Delmas C, Buisseret P (2000) Connections between the trigeminal mesencephalic nucleus and the superior colliculus in the rat. Neurosci Lett 294:17–20, https://doi.org/10.1016/S0304-3940(00)01519-6

Niedermeyer E (1997) Alpha rhythms as physiological and abnormal phenomena. Int J Psychophysiol 26:31–49, https://doi.org/10.1016/S0167-8760(97)00754-X

Oh SW, Harris JA, Ng L, Winslow B, Cain N, Mihalas S, Wang Q, Lau C, Kuan L, Henry AM, Mortrud MT, Ouellette B, Nguyen TN, Sorensen SA, Slaughterbeck CR, Wakeman W, Li Y, Feng D, Ho A, Nicholas E, Hirokawa KE, Bohn P, Joines KM, Peng H, Hawrylycz MJ, Phillips JW, Hohmann JG, Wohnoutka P, Gerfen CR, Koch C, Bernard A, Dang C, Jones AR, Zeng H (2014) A mesoscale connectome of the mouse brain. Nature 508:207–14. https://doi.org/10.1038/nature13186

Otnes RK, Enochson L (1972) Digital time series analysis. (1^st^ edition) New York: Wiley. ISBN 0471657190

Pak MW, Giolli RA, Pinto LH, Mangini NJ, Gregory KM, Vanable JW Jr. (1987) Retinopretectal and accessory optic projections of normal mice and the OKN-defective mutant mice beige, beige-J, and pearl. J Comp Neurol 258:435–46, https://doi.org/10.1002/cne.902580311

Palmigiano A, Geisel T, Wolf F, Battaglia D (2017) Flexible information routing by transient synchrony. Nat Neurosci 20:1014–22, https://doi.org/10.1038/nn.4569

Paxinos G, Franklin K (2001) The Mouse Brain in Stereotaxic Coordinates. (2^nd^ edition) San Diego: Academic Press. ISBN 0125476361

Pfurtscheller G, Lopes da Silva FH (2001) Event-related EEG/MEG synchronization and desynchronization: basic principles. Clin Neurophysiol 110:1842–57, https://doi.org/10.1016/S1388-2457(99)00141-8

Pillai AS, Jirsa VK (2017) Symmetry Breaking in Space-Time Hierarchies Shapes Brain Dynamics and Behavior. Neuron 94:1010–26, https://doi.org/10.1016/j.neuron.2017.05.013

Proix T, Bartolomei F, Chauvel P, Bernard C, Jirsa VK (2014) Permittivity coupling across brain regions determines seizure recruitment in partial epilepsy. J Neurosci 34:15009–21, https://doi.org/10.1523/JNEUROSCI.1570-14.2014

Proix T, Spiegler A, Schirner M, Rothmeier S, Ritter P, Jirsa VK (2016) How do parcellation size and short-range connectivity affect dynamics in large-scale brain network models? Neuroimage 142:135–149, https://doi.org/10.1016/j.neuroimage.2016.06.016

Qubbaj MR, Jirsa VK (2007) Neural field dynamics with heterogeneous connection topology. Phys Rev Lett 98:238102, http://dx.doi.org/10.1103/PhysRevLett.98.238102

Qubbaj MR, Jirsa VK (2009) Neural field dynamics under variation of local and global connectivity and finite transmission speed. Physica D 238:2331–46, http://dx.doi.org/10.1016/j.physd.2009.09.014

Rabinovich M, Huerta R, Laurent G (2008) Neuroscience. Transient dynamics for neural processing. Science. 321:48–50, https://doi.org/10.1126/science.1155564

Raue A, Kreutz C, Maiwald T, Bachmann J, Schilling M, Klingmüller U, Timmer J (2009) Structural and practical identifiability analysis of partially observed dynamical models by exploiting the profile likelihood. Bioinformatics 25:1923–9, https://doi.org/10.1093/bioinformatics/btp358

Rosanova M, Casali A, Bellina V, Resta F, Mariotti M, Massimini M (2009) Natural frequencies of human corticothalamic circuits. J Neurosci 29:7679–85, https://doi.org/10.1523/JNEUROSCI.0445-09.2009

Rubinov M, Sporns O (2010) Complex network measures of brain connectivity: uses and interpretations. Neuroimage 52:1059–69, http://dx.doi.org/10.1016/j.neuroimage.2009.10.003

Salami M, Itami C, Tsumoto T, Kimura F (2003) Change of conduction velocity by regional myelination yields constant latency irrespective of distance between thalamus and cortex, Proc Natl Acad Sci USA, 100(10): 6174–9, http://dx.doi.org/10.1073/pnas.0937380100

Sanz-Leon P, Knock SA, Woodman MM, Domide L, Mersmann J, McIntosh AR, Jirsa V (2013) The virtual brain: a simulator of primate brain network dynamics. Front Neuroinform 7:10, http://dx.doi.org/10.3389/fninf.2013.00010

Sanz-Leon P, Knock SA, Spiegler A, Jirsa VK (2015) Mathematical framework for large-scale brain network modeling in the virtual brain. Neuroimage 111:385–430, http://dx.doi.org/10.1016/j.neuroimage.2015.01.002

Schüz A, Chaimow D, Liewald D, Dortenman M (2006) Quantitative Aspects of Corticocortical Connections: A Tracer Study in the Mouse, Cereb Cortex 6(10):1474–86, https://doi.org/10.1093/cercor/bhj085

Sforazzini F, Schwarz AJ, Galbusera A, Bifone A, Gozzi A (2014) Distributed BOLD and CBV-weighted resting-state networks in the mouse brain, NeuroImage, 87:403–15, https://doi.org/10.1016/j.neuroimage.2013.09.050

Shah AS, Bressler SL, Knuth KH, Ding M, Mehta AD, Ulbert I, Schroeder CE (2004) Neural dynamics and the fundamental mechanisms of event-related brain potentials. Cereb Cortex 14:476–83, https://doi.org/10.1093/cercor/bhh009

Sherman SM (2001) Tonic and burst firing: dual modes of thalamocortical relay, Trends Neurosci 24:122–6, https://doi.org/10.1016/S0166-2236(00)01714-8

Shew WL, Plenz D (2013) The functional benefits of criticality in the cortex. Neuroscientist 19:88–100, https://doi.org/10.1177/1073858412445487

Spiegler A, Jirsa V (2013) Systematic approximations of neural fields through networks of neural masses in the virtual brain. Neuroimage:83:704–25. https://doi.org/10.1016/j.neuroimage.2013.06.018

Spiegler A, Hansen EC, Bernard C, McIntosh AR, Jirsa VK (2016) Selective Activation of Resting-State Networks following Focal Stimulation in a Connectome-Based Network Model of the Human Brain. eNeuro:3, https://doi.org/10.1523/ENEURO.0068-16.2016

Stefanescu RA, Jirsa VK (2008) A low dimensional description of globally coupled heterogeneous neural networks of excitatory and inhibitory neurons. PLoS Comput Biol 4:e1000219, http://dx.doi.org/10.1371/journal.pcbi.1000219

Stein BE, and Meredith MA. Functional organization of the superior colliculus. In Leventhal AG ed. The neural basis of visual function, vol 4. Basingstoke: Macmillan; 1991.

Svoboda K, Yasuda R (2006) Principles of two-photon excitation microscopy and its applications to neuroscience. Neuron 50:823–39, https://doi.org/10.1016/j.neuron.2006.05.019

Swadlow HA, Waxman SG (2012) Axonal conduction delays, Scholarpedia, 7(6):1451, http://dx.doi.org/10.4249/scholarpedia.1451

Tibshirani R, Walther G, Hastie T (2001) Estimating the number of clusters in a data set via the gap statistic. J R Stat Soc Series B Stat Methodol 63:411–23, http://dx.doi.org/10.1111/1467-9868.00293

Van de Ville D, Britz J, Michel CM (2010) EEG microstate sequences in healthy humans at rest reveal scale-free dynamics. Proc Natl Acad Sci U S A 107:18179–84, https://doi.org/10.1073/pnas.1007841107

Wang SS, Shultz JR, Burish MJ, Harrison KH, Hof PR, Towns LC, Wagers MW, Wyatt KD (2008) Functional Trade-Offs in White Matter Axonal Scaling, J Neurosci 28(15): 4047–56, https://doi.org/10.1523/JNEUROSCI.5559-05.2008

Watson C, Paxinos G, and Puelles L (2011) The Mouse Nervous System. (1^st^ edition) San Diego: Academic Press Inc. ISBN 9780123694973

Welsh DK, Logothetis DE, Meister M, Reppert SM (1995) Individual neurons dissociated from rat suprachiasmatic nucleus express independently phased circadian firing rhythms. Neuron 14:697–706, https://doi.org/10.1016/0896-6273(95)90214-7

Zingg B, Hintiryan H, Gou L, Song MY, Bay M, Bienkowski MS, Foster NN, Yamashita S, Bowman I, Toga AW, Dong HW (2016) Neural networks of the mouse neocortex. Cell 156:1096–111, https://doi.org/10.1016/j.cell.2014.02.023

